# Enzymatic carbon-fluorine bond cleavage by human gut microbes

**DOI:** 10.1101/2024.07.15.601322

**Authors:** Silke I. Probst, Florian D. Felder, Victoria Poltorak, Ritesh Mewalal, Ian K. Blaby, Serina L. Robinson

## Abstract

Fluorinated compounds are used for agrochemical, pharmaceutical, and numerous industrial applications, resulting in global contamination. In many molecules, fluorine is incorporated to enhance the half-life and improve bioavailability. Fluorinated compounds enter the human body through food, water, and xenobiotics including pharmaceuticals, exposing gut microbes to these substances. The human gut microbiota is known for its xenobiotic biotransformation capabilities, but it was not previously known whether gut microbial enzymes could break carbon-fluorine bonds, potentially altering the toxicity of these compounds.

Here, through the development of a rapid, miniaturized fluoride detection assay for whole-cell screening, we discovered active gut microbial defluorinases. We biochemically characterized enzymes from diverse human gut microbial classes including Clostridia, Bacilli, and Coriobacteriia, with the capacity to hydrolyze (di)fluorinated organic acids and a fluorinated amino acid. Whole-protein alanine scanning, molecular dynamics simulations, and chimeric protein design enabled the identification of a disordered C-terminal protein segment involved in defluorination activity. Domain swapping exclusively of the C-terminus conferred defluorination activity to a non-defluorinating dehalogenase. To advance our understanding of the structural and sequence differences between defluorinating and non-defluorinating dehalogenases, we trained machine learning models which identified protein termini as important features. Models trained on 41-amino acid segments from protein C-termini alone predicted defluorination activity with 83% accuracy (compared to 95% accuracy based on full-length protein features). This work is relevant for therapeutic interventions and environmental and human health by uncovering specificity-determining signatures of fluorine biochemistry from the gut microbiome.

**Significance:** Humans have introduced carbon-fluorine bonds into numerous manufactured compounds, including pharmaceuticals, leading to the formation of toxic fluorinated byproducts. While the human gut microbiota is known for its ability to metabolize drugs, its encoded capacity to break the strong carbon-fluorine chemical bond was previously unknown. Here we discovered that human gut microbial enzymes are capable of cleaving carbon-fluorine bonds. We developed a 96-well colorimetric fluoride assay amenable to bacterial culture-based screening. We additionally conducted whole-protein alanine scanning mutagenesis and identified through machine learning that flexible C-terminal loop residues were predictive of defluorination. Taken in the context of flexible regions of other enzyme families known to perform fluorine chemistry, this work supports using convergent structural features to predict defluorination specificity.

## Introduction

Over 20% of pharmaceuticals and 50% of agrochemicals released in the past 25 years contain one or more fluorine atom(s) (1, 2). Fluorination often improves bioavailability and extends compound half-life. However, fluorine also has a ‘dark side’ due to the production of fluorinated degradation products which can disrupt host metabolism (3). For example, key degradation products of the widely-used chemotherapeutic 5-fluorouracil (5-FU) including fluorocitrate, α-fluoro-β-alanine, and fluoroacetate, inhibit the citric acid cycle and cause severe neuro-(4, 5) and cardiotoxicity (6, 7).

The gut microbiota is known to modify and thereby inactivate or increase the toxicity of various xenobiotics (8, 9) including fluorinated compounds. Gut microbes increase the cytotoxicity of the fluorinated antineoplastic prodrug fludarabine (10) and modify the fluorinated chemotherapeutic gemcitabine through acetylation (10) or deamination (11). Anti-cancer fluoropyrimidine drugs including capecitabine and 5-FU are inactivated by gut bacteria through a pathway that mimics host metabolism (9, 12). Microbiome-dependent pharmacokinetics (9) have been linked to interindividual differences in the composition and metabolic capabilities of the human gut microbiota (13). This could impact drug activity by increasing toxicity and side effects and may potentially influence the significant interpatient variability in responses to fluorinated compounds. Given the increased use of fluorinated pharmaceuticals and the ingestion of fluorinated xenobiotics from the environment, we hypothesized that microbial carbon-fluorine bond cleavage may also be catalyzed by microbes in the human gut (14). Although there are reports of defluorinating enzymes from bacteria isolated from soil (15), ruminants (16), and activated sludge (17), none were previously associated with the human gut microbiota.

Earlier efforts to identify microbes and enzymes with defluorination activity focused on activity-guided enrichment and isolation-based strategies (16, 18). In addition to the high strength of the carbon-fluorine bond, the product of defluorination activity (fluoride anion) is also toxic to many microbes, presenting evolutionary hurdles to positive selection during enrichment and isolation (19, 20). Even when defluorination activity is detected, isolation of the responsible microbes and identification of the responsible enzyme(s) remains difficult. With the rapid growth of sequencing databases, targeted sequence-guided (15) or structure-guided (21) metagenome mining techniques are emerging as alternative approaches to identify defluorinating enzymes.

Although several different enzyme classes show promiscuous activity for fluorinated substrates (20), only hydrolases are known to cleave fluorinated compounds as their native substrates (22). Enzymes capable of hydrolytic defluorination (23) belong to two different superfamilies: the α/β hydrolase superfamily (24) and, more recently, the haloacid dehalogenase-like (HAD) superfamily (25). Although the majority of HADs actually act on a wide range of organic phosphate substrates (26), a subset of HADs are capable of regio- and stereospecific hydrolysis of halogenated (including fluorinated) carboxylic acids with a broader substrate specificity than the α/β hydrolase superfamily (25). The few defluorinating HADs described, primarily from soil, catalyze carbon-halide bond cleavage via a S_N_2-like mechanism through nucleophilic attack by an aspartate to displace the halide (25, 27). The active sites of defluorinating and dechlorinating HADs are highly similar (25), thus it remains challenging to mine metagenomic datasets to accurately identify HADs with specificity for fluorinated compounds (21). Previously, molecular dynamics (MD) showed that HADs capable of defluorination sampled more compact halide-binding pocket conformations compared to non-defluorinating HADs (25). Here,e we tested this MD-based classification method and found that it did not generalize well to a larger set of defluorinating and non-defluorinating HADs, which us motivated to search for alternative accessible features to predict metabolic defluorination potential from microbiome data.

In this study, we combined metagenome mining with in vitro experimentation to test the activity of over twenty new dehalogenases from the gut microbiome. To overcome the limitations of time-consuming ion chromatography (IC) measurements for fluoride, we developed an easy-to-use microtiter plate screen for defluorination activity. Using this approach, we screened large alanine scanning mutagenesis libraries to identify key amino acids important for defluorination. Notably, we successfully converted a dechlorinating HAD into an active defluorinating HAD by substituting its terminal 41 amino acids with those from a naturally-occurring defluorinating HAD. Molecular dynamics simulations, alanine scanning, and machine learning results converged to support the previously unknown role of the protein carboxyl (C)-terminal region of HADs in defluorination. Swapping this C-terminal region conferred defluorination capability to a non-defluorinating enzyme, thereby providing new molecular features to predict enzyme-substrate specificity for fluorinated compounds from microbiome data.

## Results

### The human gut microbiota encodes HAD-like enzymes

To investigate the potential for defluorination by human gut microbial enzymes, we first tested defluorination activity for fluoroacetate as a simple fluorinated substrate and a known 5 -FU degradation product (28). Notably, prior studies identified the soil bacterium, *Rhodococcus jostii* RHA1, as a degrader of fluoroacetate along with the discovery of its responsible defluorinating HAD superfamily enzyme classified as a L-2-haloacid dehalogenase (15). These enzymes primarily dehalogenate small chlorinated alkanoic acids with only a few demonstrating the capability to cleave the more recalcitrant carbon-fluorine bond (25). We therefore used the *Rhodococcus jostii* RHA1 HAD (Rha0230) for a protein homology search in the HumGut database (29). To our surprise, we detected over 500 HAD homologs encoded in human gut metagenomes (Fig. S1, Table S1).

The identified HAD-encoding genes were predominantly found in bacteria belonging to the phylum Bacillota, formerly named the Firmicutes (30). However, homologs were also prevalent in the Actinomycetota, Pseudomonadota, and Bacteroidota (Fig. 1A, Fig. S1). To further elucidate the relationships among these proteins, we constructed a protein sequence similarity network (Fig. S2), which revealed several distinct HAD clusters (29, 31). Two of these clusters contained characterized members of the L-2-haloacid dehalogenase subfamily (25) within the larger HAD superfamily (26). Other clusters could be assigned to other HAD superfamily family functions such as phosphonate and azetidine hydrolysis (26, 32). For subsequent analysis, we focused on clusters containing the 2-haloacid dehalogenases and curated the dataset through a multiple sequence alignment to eliminate redundant sequences and confirmed the presence of the catalytic residues (Fig. S3). We selected seven new representatives for in vitro experiments (Fig. S2, Table S2). The selected sequences from the human gut microbiota showed low amino acid sequence identity to previously characterized HADs (18% -46%; mean: 29%; Table S1). We also investigated the genomic context by extracting ± six genes on either side of the query HADs (Table S3). No CLC^F^ or Fluc (*crcB*-encoded) fluoride ion transporters or other relevant protein families were detected. Therefore, the role of these HADs in defluorination could not be inferred based on genomic co-localization. The lack of functional prediction based on sequence similarity or genomic context therefore motivated experimental validation of these unknown HADs.

**Figure 1.**
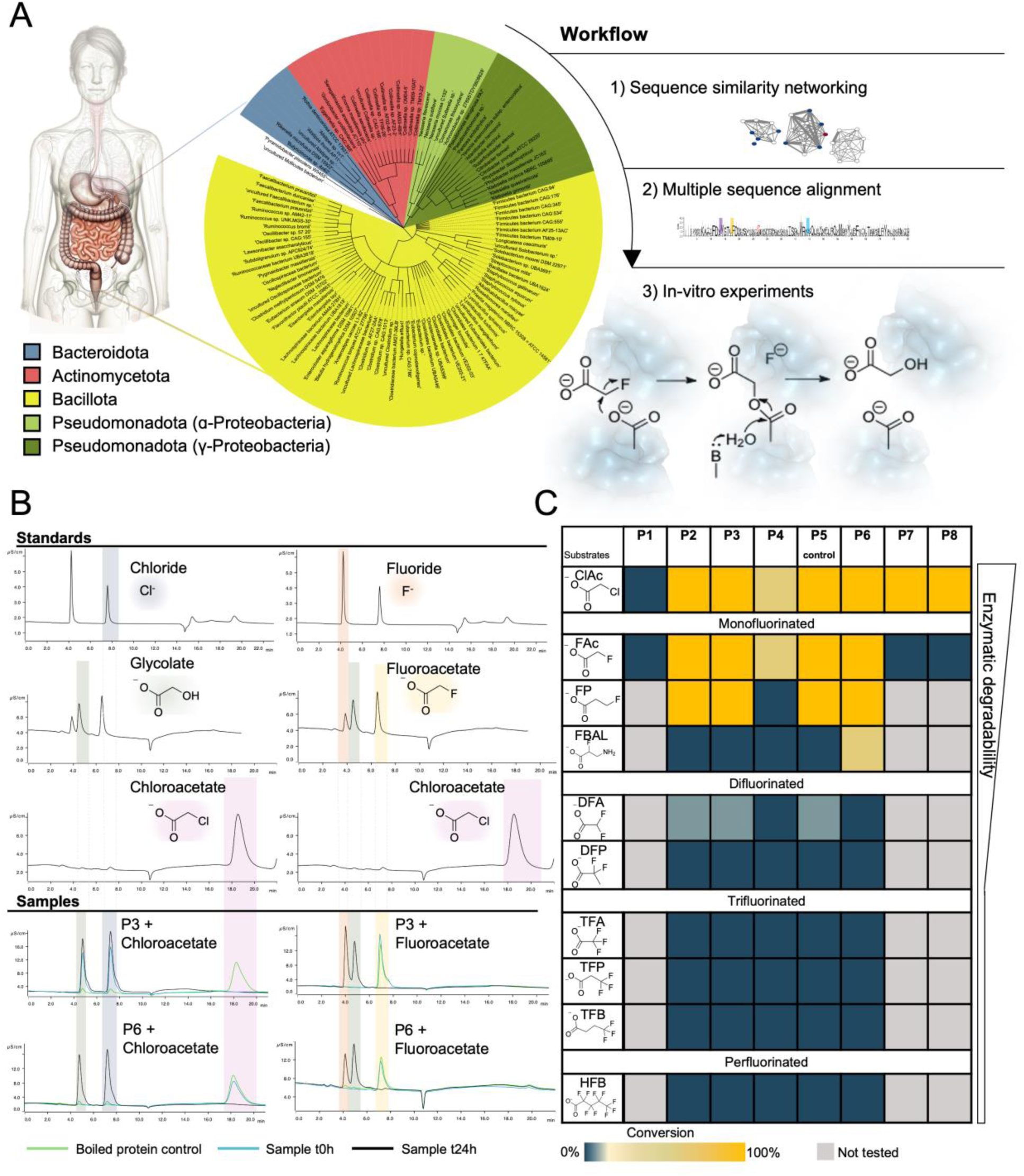
Haloacid dehalogenases are widespread across human gut bacterial phyla and exhibit broad substrate specificity. A) Overview of the identification pipeline for the selected haloacid dehalogenases. We reduced initial homology search hits using sequence similarity networking and multiple sequence alignment, resulting in eight candidates. B) Ion chromatography traces of standards colored for distinction. Blue shade: Chloride; Green shade: Glycolate; Pink shade: Chloroacetate; Red shade: Fluoride; Yellow shade: Fluoroacetate. The last four traces show protein 3 and protein 6 tested with fluoroacetate and chloroacetate. Green line: boiled protein control; Blue line: sample at t0; Black: sample after 24h incubation. The different standard peaks are color coded and dotted lines are extending the colored shading towards the sample peaks. C) Substrates specificity of ten different substrates with the eight selected proteins. The substrate conversion rate ranges from dark blue (0% conversion) to Yellow (100% conversion) (Fig. S5-S12) gray: not tested. ClAc: Chloroacetate; FAc: Fluoroacetate, FP: Fluoropropionate; FBAL: α-fluoro-β-alanine; DFA: Difluoroacetate; TFA: Trifluoroacetate; DFP:Difluoropropionate; TFP: Trifluoropropionate; TFB: Trifluorbutanate; HFB: Heptafluorobutanoate. Predicted enzymatic degradability of the substrates based on literature is indicated by the broadness of the bar.

### Human gut microbial dehalogenases are active in carbon-fluorine bond cleavage

To screen for in vitro defluorination activity, we heterologously expressed, purified, and tested seven selected gut microbial dehalogenases with ten different halogenated substrates (nine fluorinated organic acids and one chlorinated substrate, chloroacetate, Fig. 1, Fig. S4 Table S4). The HAD from the soil bacterium *Rhodococcus jostii* RHA1 (P5, Protein Data Bank (PDB) ID: 3UMG) was included as a positive defluorination control as described previously. All proteins tested except for one (*Enterocloster aldenensis*, P1, Fig. 1C) showed dehalogenase activity with at least one substrate (Fig. S5-S12). Two enzymes hydrolyzed chloroacetate but were not active with any fluorinated substrates (P7 and P8, Fig. 1C). The remaining four enzymes hydrolyzed fluoroacetate, producing fluoride ion and glycolate stoichiometrically, as measured by ion chromatography (Fig. 1B). Compounds with perfluorinated methyl (-CF_3_) or perfluorinated methylene (-CF_2_-) groups including trifluoroacetate, difluoropropionate, trifluoropropionate, trifluoropentanoate, and heptafluorobutanoate were not accepted by any of the eight HADs tested (Fig. 1C). Four of the enzymes, including the control were active with known transformation products of fluorinated xenobiotics, namely, 2-fluoropropionate, α-fluoro-β-alanine, and the polyfluorinated compound, difluoroacetate. These active defluorinating enzymes originate from diverse gut bacterial families including Eggerthellaceae (*Gordonibacter*, P2 and P3), Moraxellaceae (*Acinetobacter*, P4), and Christensellaceae (*Guopingia*, P6).

We further examined the impact of oxygen on one defluorinating enzyme P3 as a representative for our gut microbial HADs. As expected, based on the established L-2-haloacid dehalogenase mechanism (24–26), P3 cleaved fluoroacetate to produce glycolate and fluoride ions under both aerobic and anaerobic conditions (Fig. S13). To test whether we could detect the defluorination activity of the gut bacterium *Gordonibacter pamelaeae* anaerobically in gut microbial medium (GMM) with fluoroacetate and measured fluoride concentrations using a fluoride ion electrode (Table S5, Methods). Defluorination activity was not detectable, perhaps due to challenges in expression, transport, or the preferential use of other substrates by *G. pamelaeae*. Due to the limited genetic tractability of the native gut strains encoding these enzymes, we instead focused efforts on in vitro enzyme characterization. We measured the steady-state enzyme kinetics for P3 with fluoroacetate and compared our results to other defluorinating enzymes including a L-2-haloacid dehalogenase (DeHa2) and fluoroacetate dehalogenases (DeHa4) from *Delftia acidovorans* D4B (33) and the fluoroacetate dehalogenase from *Rhodopseudomonas palustris* (RPA1163, (34)). Briefly, P3 has a *K*_m_ of 59.8 ± 7.7 mM approximately one order of magnitude higher than the *K*_m_ values of DeHa2 and DeHa4 (Table S6), although lower than the *K*_m_ of RPA1163 (86.8 ± 13.7 mM)(34). The *k*_cat_ of 0.25 ± 0.02 s^-1^ is about fourfold lower than the *k*_cat_ of RPA1163 (Table S6). The weak substrate binding and correspondingly high off rate suggested by the *Km* may be a contributing factor to the overall low catalytic efficiency.

### Molecular dynamics of defluorinating and non-defluorinating enzymes reveals range of conformational compactness

Previously, MD simulations showed that HADs with defluorination activity sampled more compact conformations in the halide-binding pocket relative to non-defluorinating enzymes (25). Since the HADs identified here from the human gut microbiota were not tested in earlier studies, we first replicated the molecular dynamics (MD) simulations as described (25). In a subsequent round of simulations, we then increased the simulation time from 300 ns to 1 μs using the Amber99SB-ILDN force field instead of OPLS/A because of better performance for proteins (Fig. S14) (35, 36). Within the past few years, major advancements have been made in protein structure prediction using AlphaFold (AF) (37, 38), even enabling accurate MD simulations using relaxed AF models (39). Thus, for proteins where crystal structures were not available, we used AF to generate models (Fig. S15). In a benchmark comparison with Chan et al. (25), we obtained comparable results in MD simulations of the available X-ray crystal structures and their respective AF models (Table S7, Fig. S16). Simulating all 11 available defluorinating dehalogenases and 7 non-defluorinating dehalogenases in water for 1 μs, while tracking the size of the halide binding site, did not yield any significant differences between the two groups (Fig. 2A). Further analysis on all conserved amino acids of HADs and their relative distances between each other also showed no significant differences (Fig. S17; Table S8).

**Figure 2.**
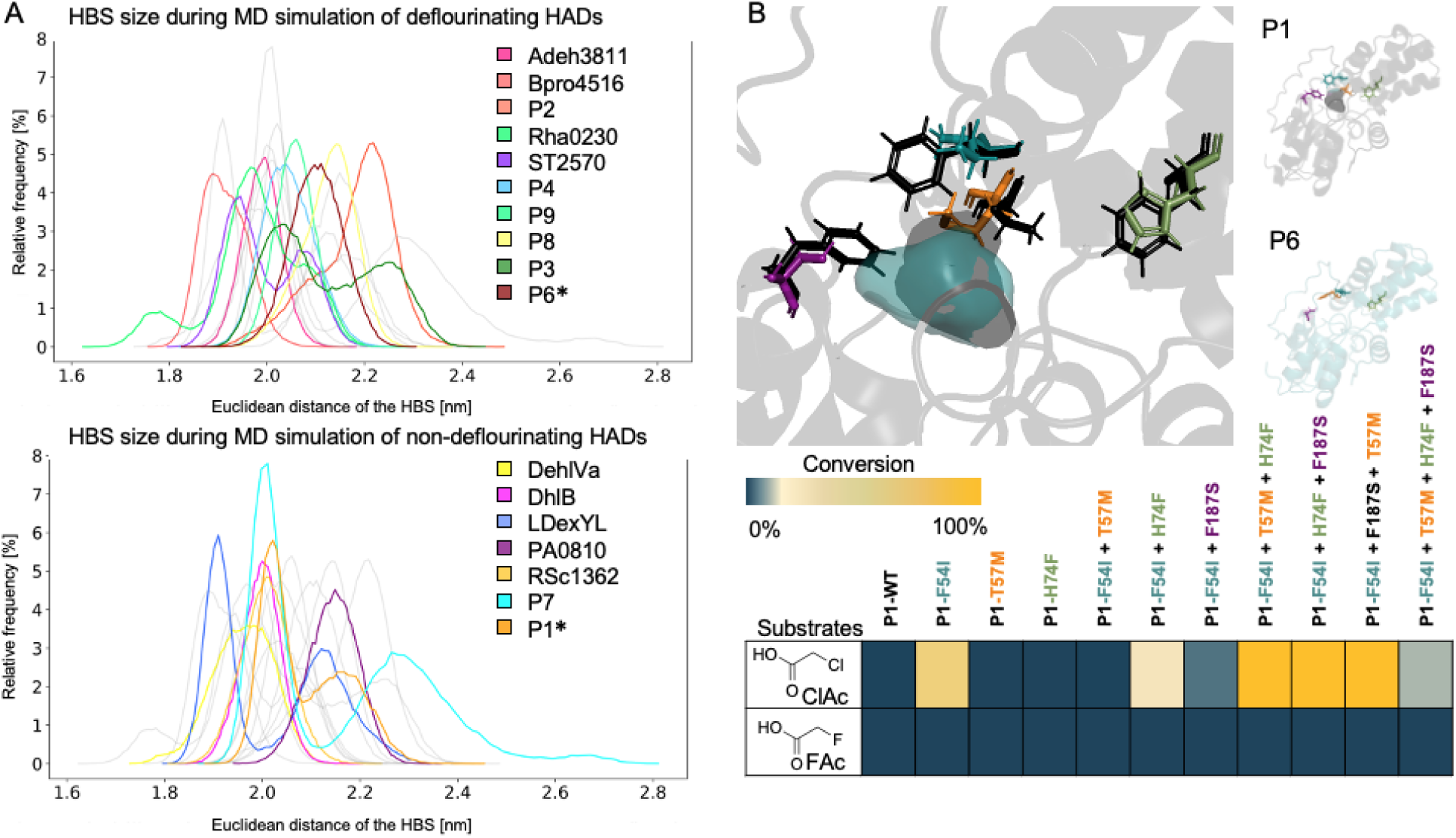
Defluorination capability does not correlate with the active site compactness or amino acid sequence identity. A) Halide binding site (HBS) size in nm during molecular simulations. Distribution of the Euclidean distances of the 3 C_α_ atoms of the HBS during simulation from 500 ns to 1000 ns. *Asterisks indicate the two enzymes used for protein engineering in this study B) Visualization of the active site of the inactive protein P1 (WP_118709078, *E. aldenensis*) black and the defluorinating protein P6 (WP_178618037, *G. tenuis*) teal. The variants introduced in P1 (*E. aldenensis*) are F54I (blue), T57M (orange), H74F (green), and F187S (purple). Substrate specificity of the different protein variants with the two substrates chloroacetate (ClAc) and fluoroacetate (FAc).

We used MD simulations to further investigate the differences between enzymes capable of defluorinating difluoroacetate (DFA). Previous research on DFA defluorination indicated a smaller active site volume in the fluoroacetate dehalogenases with higher DFA activity (24). Therefore, we compared the size of the halide-binding site in the three enzymes (P2, P3, and P5) capable of defluorinating DFA with those in P4 and P6 (Fig. S18). We observed no significant difference in the distribution of halide-binding pocket sizes that could explain the altered substrate preference. We note, however, that the distributions of Euclidean distances for some of the HADs were multimodal, suggesting distinct conformational states which are likely to include both catalytically active and inactive states. This requires more in-depth investigation but does not change the overall conclusions that the MD simulations alone are insufficient to distinguish defluorinating from non-defluorinating HADs.

We conducted additional experiments to confirm that the fluoride ion release is not solely due to fluoroacetate (FAc) contamination in the DFA (Fig. S19). However, DFA activity is weak, and it remains unknown whether the enzyme itself contributes to the second defluorination step, as the intermediate gem-fluoro alcohol is likely to undergo spontaneous HF elimination to form a ketone (40).

### Phylogenetic analysis does not differentiate defluorinating and non-defluorinating enzymes

To further investigate methods to differentiate defluorinating and non-defluorinating enzymes, we generated a phylogenetic tree of the HAD-like family protein sequences (Fig. S20, Table S9). The dehalogenating enzymes originated from eight diverse bacterial and archaeal phyla (Fig. S21) which did not group based on the taxonomy of the source organisms. The tree topology revealed that the defluorinating and non-defluorinating enzymes did not form separate or easily distinguishable clades.

### Rational active site engineering converts an inactive enzyme to a dechlorinating enzyme

As neither MD simulations nor phylogenetic analysis yielded conclusive results to distinguish defluorinating and non-defluorinating HADs, we next sought to experimentally test the role of specific residues in the active site. We used two of the seven proteins tested in vitro sharing the highest sequence and structural similarity but distinct activities from each other (Fig. S21) (54% sequence identity, 68% sequence similarity). P1, a non-dehalogenating enzyme, and P6, a defluorinating enzyme. Based on comparison between the active sites of the two enzymes, there were only four residues that were significantly different in the active site vicinity including the halide-binding pocket and substrate tunnel (Fig. S21). We predicted these residues may provide substrate specificity for defluorination, therefore we first aimed for rational engineering of the non-dehalogenating enzyme P1 (*E. aldenensis*) via site-directed mutagenesis to generate single, double, and triple variant combinations as well as the full quadruple variant P1 (F54I, T57M, H74F, F187S) (Fig. S22-S25).

Among the single variants tested, only one (F54I) showed dehalogenation activity. The F54I variant hydrolyzed chloroacetate but did not hydrolyze fluorinated substrates. Based on structural modeling, F54 is positioned at the entrance channel to the substrate binding pocket. In the non-dehalogenating protein P1 (*E. aldenensis*) the larger phenylalanine may exclude chloroacetate at the binding pocket entrance. In contrast, the dehalogenating protein P6 (*Guopingia tenuis*) features an isoleucine at this position, potentially enabling access for chloroacetate. While we did achieve a functional dehalogenase through the single-point mutation in P1 (*E. aldenensis*), the individual active site changes did not singularly confer defluorination activity. We hypothesized epistatic interactions may promote defluorination. We therefore used the F54I variant as the template for generating double, triple, and quadruple active site variants. In the P1 (F54I, H74F) double variant and all triple variants we achieved improved dechlorination activity (Fig. 2B). Surprisingly, we did not observe defluorination activity in any of the rationally engineered variants, even though the active site residues were 100% identical to the defluorinating reference enzyme P6 (*G. tenuis*). This result suggests that additional second shell interactions or other conformational changes (41) not localized in the active site may be critical for defluorination activity.

### Development of a new microtiter well fluoride detection assay for alanine scanning

In order to test the contributions of a wider set of amino acids in defluorination e.g., peripheral residues in addition to active site residues, we used alanine scanning mutagenesis to generate full alanine variant libraries of the proteins: P6 *(G. tenuis*) and P1 (F54I, T57M, H74F, F187S) (*E. aldenensis*). In total, the library consisted of 474 single alanine variants. To screen the alanine library, we devised a microtiter well plate colorimetric screening method for defluorination to avoid lengthy ion chromatography steps for analysis. Previously, a lanthanum-alizarin assay for microtiter well plates (42) and a xylenol orange assay for an agar plate-based method (43) were reported. We tested the lanthanum-alizarin assay (42) and additionally developed a miniaturized liquid adaptation of the xylenol orange screening method. Both assays yielded a clear change in absorbance in the presence of free fluoride ions as validated by ion chromatography results (Table S10). Due to its enhanced color intensity, we report here the results of screening using the miniaturized xylenol orange assay. As the xylenol orange assay has not been described previously for use in liquid, microtiter well plate format, a detailed methods section and list of interfering substances is included (Fig. S26-S27, Table S11). In our assay, sodium phosphate monobasic was the compound with the highest observed interference in the present fluoride assay which can be reduced to below the limit-of-detection with washing steps (Figure S28, Methods).

To test our miniaturized screening method, we selected eleven candidate dehalogenases spanning eleven additional gut microbial genera (Table S12). Based on the colorimetric screen, we detected an active gut microbial defluorinase from *Priestia megaterium* sharing less than 35% amino acid identity to our reference defluorinase (PDB ID: 3UMG). We purified the *P. megaterium* enzyme to homogeneity and validated its defluorination activity using ion chromatography (Fig. S29). This highlights the potential of microtiter-well assays to rapidly identify defluorinating enzymes.

With our new microtiter defluorination assay in hand, we next screened the alanine scanning library with a dual-assay approach for both defluorination and dechlorination activity (Fig. 3A). The screening was conducted in two independent experimental campaigns each with biological replicates as further described in the Methods section, with and without dilution, and normalization for protein concentration (Experiment 1 and Experiment 2, respectively). Across independent experiments and biological replicates of the alanine scanning library (Methods), 39.8-41.7% of alanine variants lost defluorination activity. A slightly lower proportion of variants (33.0-33.5%) lost dechlorination activity in Experiment 1. In Experiment 2, 31%-37% of alanine variants lost defluorination activity and 39-41% lost dechlorination activity. We identified the top 19 alanine variants which showed dechlorination activity but lost defluorination activity in both experiments (Fig. S32-S34, Table S14-15). Despite variability among replicates within and across Experiments 1 and 2 caused by inherent biological differences in growth and expression—remaining even after protein normalization correction (Fig. S30-31)—both experiments identified the same important regions and residues linked to defluorination (Fig. S35, Table S13). Subsequent machine learning results are reported here for Experiment 1 for clarity (Exp. 2 results are reported in Table S16 and Fig. S36).

**Figure 3.**
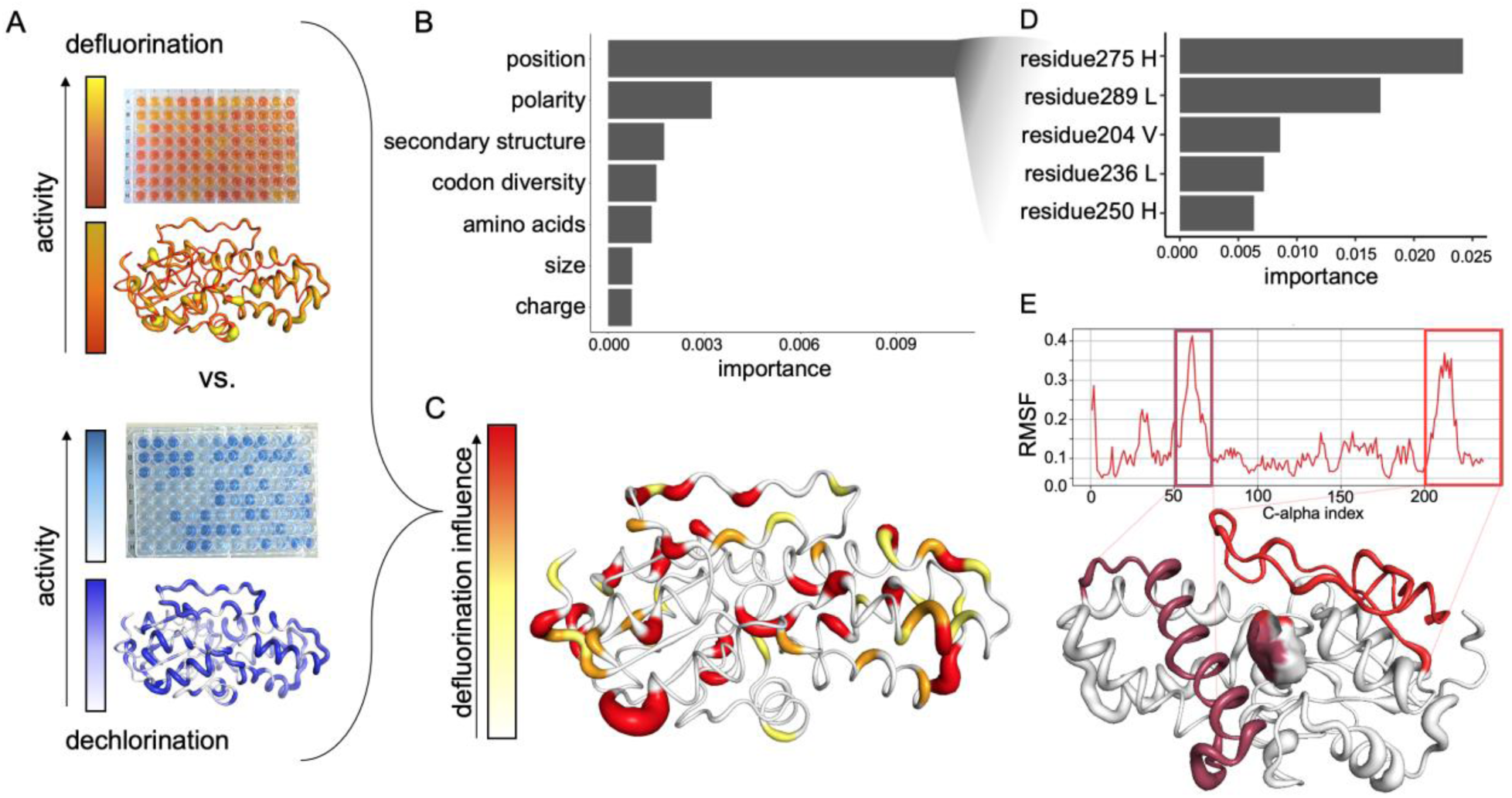
The intrinsically disordered C-terminal region is important for defluorination activity. A) Results of Experiment 1 colorimetric assays for defluorination and dechlorination activity, respectively, mapped onto the P6 protein model. The color gradient from white (low activity) to yellow (medium activity) to red (high activity) and line thickness is used to visualize activity differences based on location in the wild-type protein structure. B) Mean relative feature importance plot extracted from random forest regression model using alanine scanning data to predict defluorination activity. C) Protein ‘hotspots’ important for defluorination and dechlorination based on the alanine scanning data, where line thickness and coloring correspond to a higher importance for defluorination alone. D) Top five residue positions with the highest mean Gini importance in random forest classification for defluorinating vs. non-defluorinating proteins E) Root mean square fluctuation (RMSF, reported in nm) of the C_α_ atoms indicates high movement in the two regions of the protein. The two boxes highlight the more flexible regions in the proteins and lines connect the corresponding amino acids in the protein visualization. Maroon = amino acids 36-69, Red = amino acids 197-235. The surface of the binding pocket location is shaded by proximity to the two regions.

### Machine learning highlights important protein regions for defluorination

To analyze our alanine scanning data, we used the random forest algorithm to train a machine learning model to predict defluorination activity. Random forest is suitable for both regression and classification tasks even under low-data regimes (n=474 data points from the alanine scanning experiments). Our regression model trained on 80% of the alanine scanning data was capable of predicting quantitative changes in defluorination activity as the result of single alanine point mutations with the remaining 20% of the data with a root-mean square error (RMSE) of 0.2 ± 0.02 enzyme activity units (on a min-max normalized activity scale between 0-1 defluorination activity). Based on the relative feature importance, the most predictive feature was the position of each amino acid within the protein sequence rather than the size, charge, identity, codon diversity, or polarity of each amino acid converted to alanine (Fig. 3B). These results showing position carried the highest importance for defluorination activity corresponds with the visible ‘defluorination’ hotspots on the protein structure (Fig. 3C).

To train a more generalizable model with applications beyond our selected alanine scanning libraries P1 (F54I, T57M, H74F, F187S) and P6, we used random forest to classify defluorinating vs. non-defluorinating enzymes based on general amino acid features extendable to untested HAD sequences (Fig. 3D). We obtained 95% training and test set classification accuracy (F1 - score = 0.928 ± 0.028, Table S16). To test the generalizability of our classification model, we used the model to predict defluorination activity for a new set of five uncharacterized HAD homologs (Methods, Fig. S35). We obtained a defluorination probability score of 0.9 for a HAD (WP_139652913.1) encoded in the genome of *Raoultibacter phocaeensis* Marseille-P8396^T^ isolated from human stool samples (44). We heterologously expressed and confirmed positive defluorination activity in the *R. phocaeensis* HAD. The other four HADs in the dataset expressed well in *E. coli* but were not active in defluorination. Our models predicted a low defluorination probability of 0.5 or less (Fig. S37, Table S17), supporting the ability of our models to discern high-probability defluorinating HADs from others.

Although recognizing the limited scope of this model to HAD-like family enzymes, this classification analysis pinpointed residues that were important for defluorination activity (Fig. S35-36). Strikingly, the top five most important residues (Experiment 1) for defluorination activity prediction accuracy were localized in the C-terminus (final 41 amino acids). A simplified model trained on only the 41 C-terminal residues as features still achieved 83% testing and training set prediction accuracy (average F1-score = 0.728 ± 0.129, Experiment 1, Table S16).

Additionally, we computed the root mean square fluctuations (RMSF) from our molecular dynamics (MD) simulations of P6 (*G. tenuis*), allowing us to compare the spatial fluctuations of different regions within the protein structural model. Our analysis identified two regions exhibiting high RMSF values. Both regions are situated at the channel entry leading to the active site (Fig. S38), with one of them located at the C-terminus (Fig. 3E). This finding aligns with the residues implicated in defluorination, as determined by both machine learning predictions and experimental evidence.

### Non-defluorinating enzyme gains defluorination function by C-terminus swap

To test the relevance of the C-terminal region experimentally, we generated chimeric proteins by combining different regions from the quad variant non-defluorinating protein P1 (*E. aldenensis* F54I-T57M-F74H-F187S) with the defluorinating enzyme P6 (*G. tenuis*) (Fig. S39). Strikingly, the single P1 quad variant (*E. aldenensis* F54I-T57M-F74H-F187S) chimera generated by swapping in the 41 amino acid C-terminus of the P6 (*G. tenuis*) enzyme showed slight defluorination activity (Fig. 4A). All other chimeric proteins were active in dechlorination, but not defluorination (Fig. 4A, Fig. S39). Notably, the C-terminal region has not been previously reported for its importance in dehalogenation or fluorinated compound substrate specificity. Analyzing our alanine scanning data based on our chimeric segments (Fig. 4B), we observed a pattern in the activity distribution of defluorination activity (Fig. 4B). Based on our MD simulations, we observed that the C-terminal chimeric segment also had the highest overall movement relative to other protein chimeric segments (Fig. 4C). Overall, the combination of alanine scanning and chimeric mutagenesis indicated that the C-terminal 41 amino acid segment was sufficient to install defluorination activity when fused to a non-defluorinating homologous enzyme.

**Figure 4.**
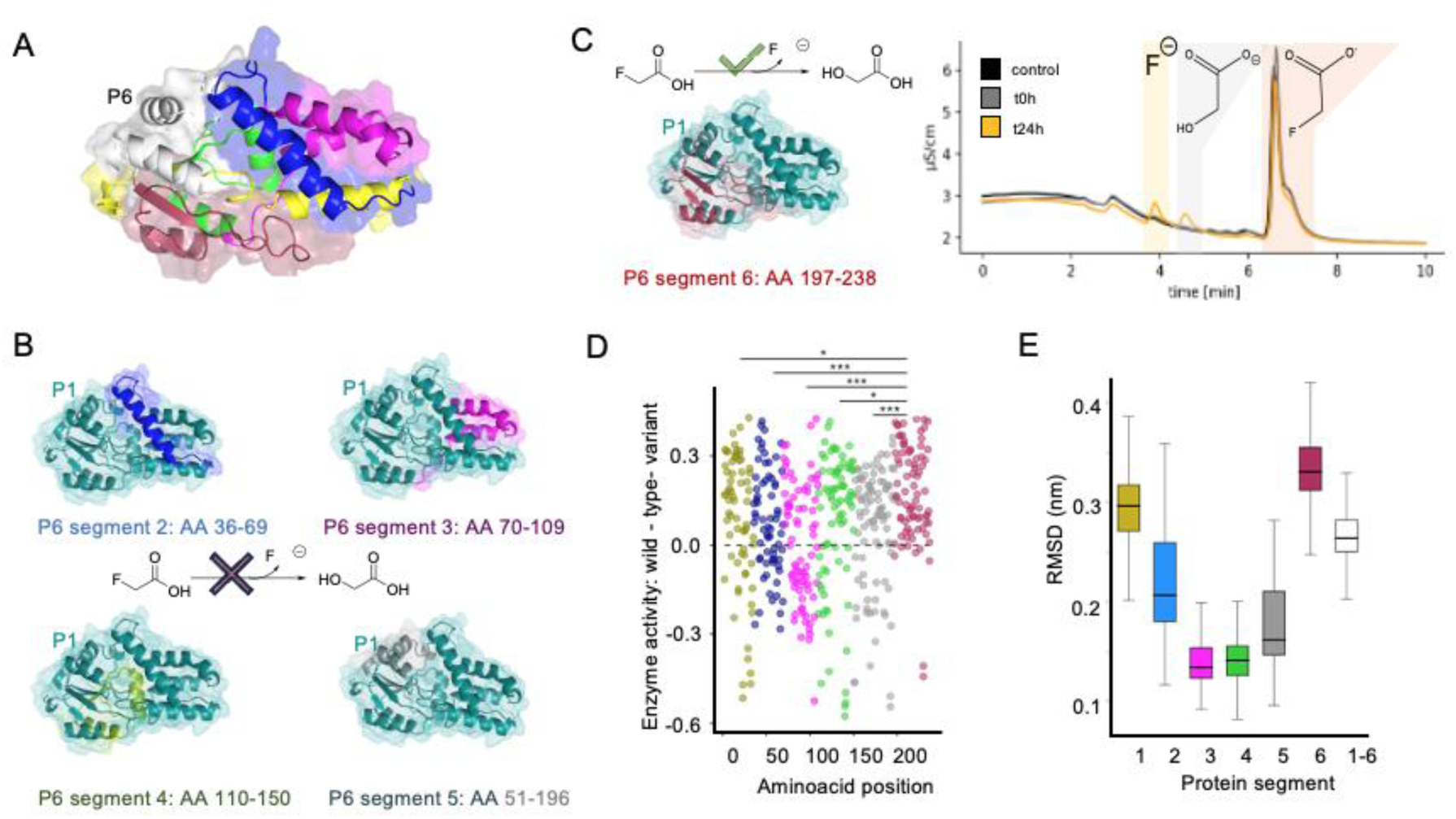
Swapping the C-terminus of a non-defluorinating HAD with the C-terminus of a defluorinating homologous enzyme enables defluorination activity. A) Representation of the defluorinating protein P6. The colored parts represent the exchanged segments used for generating the chimeric protein variants with P1. B) The protein variants with the colored segment from P6 showed no defluorination activity. C) The engineered chimeric protein with a swapped C-terminal segment enables weak conversion of fluoroacetate to glycolate and fluoride ion. D) Protein defluorination activity of the alanine scanning library with all data points shown for two independent biological replicates (Experiment 2 results are visualized, both Experiment 1 and Experiment 2 results are provided in Table S13-14, Table S18). The x-axis indicates the position in P6 and the y-axis indicates the change in min-max normalized activity for each alanine scanning variant relative to wild-type activity (dotted gray line). Points are colored by chimeric segments as depicted in other figure panels. The C-terminal chimeric segment exhibits a significantly different distribution of active to inactive alanine variants from the other chimeric segments of the protein (p<0.05, Kruskal-Wallis test). E) Boxplot of root mean square deviation (RMSD) in nm for different protein segments derived from MD simulations with the C-terminal chimeric segment exhibiting the highest overall movement.

## Discussion

Here we demonstrated that enzymes capable of carbon-fluorine bond cleavage are widely distributed across bacterial phyla common in the human gut. To our knowledge, this is the first report of HADs sourced from the human gut microbiota capable of catalyzing this reaction. Despite sharing sequence and structural similarity, the proteins in this study exhibited variable substrate specificity including activity with fluoroacetate and α-fluoro-β-alanine. Both of these are known toxic transformation products (6, 7) of 5-fluorouracil (5-FU), a widely-used cancer medication on the World Health Organization’s list of Essential Medicines (45). The anticancer mechanism of 5-FU is primarily based on the inhibition of thymidylate synthase (TS) by 5-fluoro-2’-deoxyuridine monophosphate (FdUMP). FdUMP competes with deoxyuridine monophosphate by forming a covalent ternary complex with TS and methylenetetrahydrofolate (46). This mechanism-based inactivation has been a significant driving force behind therapeutic innovations targeting not only human, but also bacterial enzymes, such as α-fluoromethyl ketone bile acid analogs which covalently inhibit gut bacterial bile salt hydrolases (47). Studies on fluorinated substrate analogs that inhibit pyridoxal phosphate (PLP)-dependent alanine racemases (48) as well as other enzyme classes (49), have advanced the discovery and characterization of enzyme catalytic mechanisms and their physiological roles. A notable example is the discovery of a *(S)*-α-fluoromethyltyrosine inhibitor of a microbial PLP-dependent tyrosine decarboxylase. This previously unidentified microbial enzyme converts levodopa, a medication used to treat Parkinson’s disease, into dopamine, thereby contributing to interindividual variation in drug efficacy (50). Collectively, these findings and our study contribute to a growing mechanistic understanding of how diverse gut microbial enzymes interact with fluorinated substrates.

In this study, we also identified four new HADs active on DFA, doubling the number of hydrolytic dehalogenases experimentally shown to defluorinate -CF_2_ groups (24). DFA is a transformation product of various xenobiotics including the refrigerant R1113, chlorotrifluoroethylene, (51) and the anesthetic 1,1,2,2-tetrafluoro-1-(2,2,2-trifluoroethoxy)-ethane (52). DFA is also a byproduct of the photodegradation of penoxsulam, a widely used pesticide (53), thereby exposure to the gut microbiota may occur via the consumption of pesticide residues on food. In general, fluorinated substances seep into groundwater and drinking water, where they are subsequently consumed by humans, thus coming in contact with the gut microbiota, although there is a lack of quantitative exposure data to fully assess the extent and impact of this interaction.Additionally, this study is limited in its scope to overexpressed recombinant HADs sourced from the gut microbiota. In vivo and pure culture experiments remain inconclusive due to unknown conditions for gene expression and interference from complex matrices, requiring further investigation.

Specificity-determining residues or structural features that differentiate defluorinating enzymes from their non-defluorinating counterparts in the L-2-haloacid dehalogenases have remained elusive (25). Alanine scanning proved useful to identify protein variants that retained dechlorination activity but lost their defluorination capability and to pinpoint several amino acids critical for defluorination. Surprisingly, these amino acids were primarily located in the peripheral regions of the protein with a higher percentage of significant residues located in the C-terminal segment of the protein. These observations, along with results from more recent studies (54), support the growing complex understanding of protein function beyond the simplistic lock-and-key model (55). Consequently, our protein engineering efforts shifted towards engineering chimeric proteins by replacing segments of non-defluorinating protein variants with segments from a defluorinating enzyme. We observed chimeric proteins with weak, but robust and reproducible defluorination activity only when the C-terminal segment was exchanged.

The C-terminal regions of proteins have been extensively studied in eukaryotic cells (56, 57), where ‘short linear motifs’ within intrinsically disordered regions (58) have been associated with a broad spectrum of functions, including substrate recognition (56, 59), signaling (60), and disease risk (56, 61). This functional diversity is attributed to the solvent exposure and higher degree of disorder of C-termini, which facilitate protein-protein interactions (62) and oligomerization facilitated through natural C-domain swapping (63). Previously, in a HAD superfamily phosphatase, the flexible C-terminal tail was shown to interface with adjacent monomers to facilitate product release (41). Our MD simulations suggested gut microbial HAD C-termini exhibit greater flexibility than other protein regions. Combined with a ‘gain-of-function’ through the design of a C-terminal chimera, we propose the C-terminal region of HADs plays a significant and previously unreported role in defluorination. This knowledge lays the foundation to engineer gut microbial proteins with expanded substrate ranges for carbon-fluorine bonds, opening new applications in human and environmental health. It adds a prokaryotic dimension to the ‘C-terminome’ to more accurately predict and modulate enzyme functions in microbiomes and across the domains of life.

## Data and code availability

Scripts and data associated with analyses presented in this manuscript are available at: https://github.com/MSM-group/gut_microbe_defluorination_paper

## Acknowledgements

We acknowledge Michael Zimmermann, Mitchell Levesque, and Larry Wackett for their thoughtful discussions and input. We acknowledge René Gall for his technical support with analytics and Till Epprecht for his support in computational analysis. We thank the entire Joint Genome Institute team including Miranda Harmon-Smith, Yasuo Yoshikuni, and many more for their exceptional support. The work (proposal: 10.46936/10.25585/60008420) conducted by the U.S. Department of Energy Joint Genome Institute (https://ror.org/04xm1d337), a DOE Office of Science User Facility, is supported by the Office of Science of the U.S. Department of Energy operated under Contract No. DE-AC02-05CH11231. S.P. was supported by postdoctoral funding from the Peter und Traudl Engelhorn Stiftung. V.P. and S.L.R. acknowledge funding from the Swiss National Science Foundation (grant no. PZPGP2_209124), the Uniscientia Foundation, and the Helmut Horten Foundation (2022-YIG-090).

## Author contribution

Conceptualization, S.L.R and S.I.P.; Methodology, S.L.R, S.I.P. F.F., V.P., R.M. and I.K.B.; Investigation, S.L.R, S.I.P. F.F., V.P., R.M, and I.K.B.; Writing – Original Draft, S.L.R., S.I.P., V.P., and F.F.; Writing – Review & Editing, S.L.R., F.F., I.K.B., R.M., V.P., and S.I.P.; Funding Acquisition, S.L.R. and I.K.B.; Supervision, S.I.P., I.K.B., and S.L.R.

## Declaration of interests

The authors declare no competing interests.

## Methods

### 1. Identification and selection of HADs from the HumGut Database

We retrieved protein sequences of known environmental HADs (2-haloacid dehalogenase subfamily) previously characterized for their defluorination activity (25) from the National Center for Biotechnology Information (NCBI, WP_011593529.1 and WP_011481495.1). The sequences were used as queries for BLAST searches within the HumGut human gut genome collection (Version: February 2023) (29). A BLASTdb was created using the 30,691 HumGut genomes and a tblastn search was performed to identify homologs of the WP_011593529.1 and WP_011481495.1 proteins. The 500 sequence hits with bitscores greater than 40 were subjected for a reverse blast search against the NCBI database. Hits that could be mapped to NCBI database entries with 100% sequence identity and 90-100% query coverage were further curated for redundancies. The resulting list of 114 protein sequences (Table S1) were aligned using Clustal Omega within the EMBL-EBI Framework (64) and the presence of the catalytic active residues were verified. Genomic context for the query sequences were extracted using the genome mining tool RODEO v2.0 (65). Code and query sequences are available at https://github.com/MSM-group/gut_microbe_defluorination_paper. Subsequently, we used the EFI-EST webtool to generate a sequence similarity network (SSN) with default parameters (66). We visualized the 95% amino acid identity representative network using Cytoscape (67) (Fig. S2) and seven SSN cluster representatives were selected for experimental validation (Table S2). Genes were synthesized as described below for in vitro work. In a second ‘validation’ round, 11 new proteins were selected based on high sequence similarity alignment scores for additional experimental validation (Table S12).

### 2. Cloning and heterologous expression of candidate enzymes

The first batch of full-length dehalogenases (n=7) were codon-optimized for *E. coli* codon usage using the Integrated DNA Technologies (IDT) Codon Optimization Tool. The sequences were synthesized as gBlocks by Integrated DNA Technologies. The gBlocks were cloned individually by Gibson Assembly into multiple cloning site 1 of pCFDuet-1 vectors (spectinomycin/streptomycin resistance marker) with tobacco etch virus (TEV) cleavage sites and C-terminal hexahistidine tags (Table S19-S21). Constructs were transformed into *E. coli* DH5α cells and verified by Sanger sequencing (Microsynth, Balgach, Switzerland). Sequence-verified constructs were transformed into *E. coli* T7 Express cells for protein production (New England Biolabs Frankfurt, Germany, NEB C2566I). The second batch of synthesized genes for colorimetric assay validation (n=11) were cloned by the U.S. Department of Energy Joint Genome Institute. These were constructed identically to the first batch with the exception of using the build-optimization software, BOOST (68) for codon optimization and expression in T7 Express *E. coli*. All codon-optimized sequences are available at: github.com/MSM-group/gut_microbe_defluorination_paper/tree/main/protein_sequences_info

Starter overnight cultures (5 mL) were grown in LB with 100 μg/mL spectinomycin overnight at 37°C. From the starter culture, 0.5 mL were added to 200 mL Terrific Broth with 100 μg/mL spectinomycin in a 1 L baffled Erlenmeyer flask and grown to an optical density of 0.5-0.8 at 37°C. Protein expression was induced by adding 0.1 mM isopropyl β-D-1-thiogalactopyranoside (IPTG) and cultures were incubated for 24h at 16°C.

For scale up and purification, induced cultures were poured into 50 mL Falcon tubes and centrifuged at 4000 RCF for 30 min. The supernatant was decanted, and cell pellets were resuspended in 4 mL buffer A (20 mM Tris-HCl, 500 mM NaCl, 20 mM imidazole) per gram cell pellet, 10% glycerol, pH = 8) per gram of pellet weight. Cells were lysed by 3 cycles of sonication using a 6 mm needle, 30% amplitude and 10 x 10 second pulses with a rest time of 10 seconds in between (total sonication time per cycle = 3 minutes). Lysate was centrifuged at 20,000 RCF at 4°C for 45 min. For benchtop purification, supernatants were applied to a column packed with Ni-NTA agarose beads (Qiagen) pre-equilibrated in buffer A. The column was subsequently washed with buffer A and then subjected to two washes with buffer B (40 mM Tris-HCl, 500 mM NaCl, 10% glycerol and 40 mM imidazole). Enzymes were eluted from the column using buffer C (40 mM Tris-HCl, 500 mM NaCl, 10% glycerol, and 500 mM imidazole) The fractions with the highest protein concentration was determined qualitatively by Bradford assay(69). Imidazole and NaCl were removed by running the sample over a PD-10 desalting column (Cytiva) and eluting in buffer (100 mM Tris-SO_4_, pH = 8.5). The concentration of the purified proteins were determined by Qubit™ protein assay and the proteins were flash frozen using liquid nitrogen in 500 μL aliquots and subsequently stored at -80°C.

### 3. Assessment of enzyme-substrate specificity

Stock solutions containing 100 mM of the test substrates listed in Table S5 dissolved in water were prepared. The concentration of the Ni-NTA purified proteins were determined by using the Qubit™ protein assay and the proteins were diluted to a final concentration of 0.5 mg/mL in 100 mM Tris-SO_4_ buffer, pH 8.5. As controls the proteins were boiled for 10 min at 100°C. The test substrates were added to a final concentration of 1 mM and the samples were incubated at 37°C. 50 μL samples were taken for the different timepoints and mixed with 50 μL acetonitrile to quench the reaction. The samples were centrifuged at 13 000 RPM for 5 min and the supernatant transferred into plastic vials for ion chromatography (IC).

An IC system (Metrohm, Switzerland) was used consisting of an IC sampling center 889 coupled to an ion chromatograph (930 Compact IC Flex) with a conductivity detector. Samples were incubated at 10°C in the sample center with 10 μL injection volumes first onto a precolumn (Metrosep A Supp 16, Guard, 4 mm) and subsequently onto a Metrosep A Supp 16 column (250 x 4.0 mm, 5 μm). The temperature in the column was kept constant at 30°C. The eluent consisted of 2.0 mM Na_2_CO_3_ and 5.0 mM NaHCO_3_. The flow rate was set to 0.8 mL/min. IC data processing and quantification was performed using MagIC Net Professional (version 3.3).

All experiments were conducted at least in triplicates.

### 4. AlphaFold and structural comparison

Structural models for the seven initial dehalogenase sequences selected from the HumGut databases, were generated using AlphaFold2 (AF) ColabFold(70). Models with the highest rank for each dehalogenase were prepared with relaxation using the Amber99SB force field(71). Additionally, we used AF to generate models for the HADs PA0810, RSC1362, ST2570 and RHA0230. The accuracy of AF models was compared to known X-ray crystal structures, where available (25).

### 5. Molecular dynamics simulations

Molecular dynamics simulations were performed using the GROMACS package version 2022.5 (72, 73) compiled with CUDA support and run on NVIDIA GeForce RTX 2080 Ti graphic cards. The crystal structures for HADs PA0810, RSC1362, ST2570 and RHA0230 were downloaded from RCSB PDB (PDB IDs: 3UMC, 3UMB, 2W43 and 3UMG, respectively. For PDB files which contained selenomethionine, selenomethionine was exchanged with methionine. Relaxed AF models were also subjected to simulations. The system was prepared with GROMACS tools, parameterized with the force field Amber99SB-ILDN (74), and solvated in explicit water using the default water model TIP3P (75) in a cubic box ∼1.2 nm bigger than the protein in each direction. The protein’s net charge was neutralized with Na^+^ or Cl^-^ ions. Conjugate gradient energy minimization was performed with a minimum of 50 steps and a maximum of 1000 steps or until the energy of the system reaches below 1000 kJ/mol/nm. After a short (200 ps) NVT equilibration and (200 ps) NPT equilibration, the production runs were carried out. The simulations were performed under periodic boundary conditions for a total simulation length of 0.1 µs with ten replicates and 1 µs with two replicates using the leap-frog algorithm with time steps of 2 fs for each simulation. The bonds involving hydrogen atoms were constrained with the LINCS algorithm (76). The cutoff scheme for nonbonded interactions was Verlet with a cutoff of 1.2 nm (for all AMBER force fields). The Coulomb interactions were calculated using the particle mesh Ewald algorithm (77) and a Fourier spacing of 0.14 nm. The temperature was controlled using a modified Berendsen thermostat (78) with a coupling time of 0.1 ps (protein & non-protein coupling groups) and a reference temperature of 300K. The pressure was controlled using the Berendsen barostat (79) an isotropic pressure coupling in x-y-z directions, a coupling time of 1.0 ps, a reference pressure of 1.0 bar, and an isothermal compressibility of 4.5 x 10^-5^ bar^-1^. The variance of the 100 ns simulations with 10 replicates ranged between 0.015 and 0.08 nm for all simulated proteins. For the 1 µs runs the variance between the two replicates ranged between 0.0016 nm and 0.031 nm. The root mean square deviation (RMSD) of the simulated proteins relative to the starting structure (C_α_ and all atoms) was measured using gmx_rms for both whole proteins and for the protein segments with the necessary index files generated with make_ndx. The three active site residues of the halogen binding site (aspartate, asparagine and arginine) were identified and the Euclidean distance between the three C_α_ atoms was tracked during the simulation after an equilibration period of 500 ns with the built-in tool gmx_distance (73).. To measure the Root Mean Square Fluctuation (RMSF) of all C_α_ atoms during the simulation the built-in function gmx_rmsf was used.

### 6. Sequence based comparison of HADs

Multiple sequence alignment and maximum-likelihood phylogenetic analysis (Fig. S20-S21, Table S9) were performed to gain a better perspective on the sequence features of the HAD-like enzymes. We performed these two analyses with a data set of the entire HAD-like family available in UniProt (80) for 248 sequences available as of August 2023 with experimental evidence characterized at the protein level. A multiple sequence alignment of the resulting 248 proteins was performed using the Clustal Omega algorithm (81) with a Biopython wrapper (82) and the phylogenetic tree (Fig. S20-21) was generated using the IQ-TREE algorithm (83, 84). The model selection was performed by IQ-TREE model finder function (83). The best model was selected (substitution model Q.pfam+R8) and a maximum-likelihood tree was constructed with 1000 bootstrap replicates and visualized using ETE 3 (85).

### 7. Rational active site engineering

To identify potentially significant amino acids involved in the defluorination process, we used two of seven in vitro tested proteins sharing high structural and sequential similarity but distinct activities from each other (Fig. S21) (54% sequence identity, 68% sequence similarity). WP_1187090781.1, a non-defluorinating enzyme, and WP_1786180371.1, a defluorinating enzyme. We targeted amino acids within WP_1187090781.1, particularly those situated in or near the active site, for substitution with their counterparts from WP_1786180371.1. Four single variants were constructed: F54I, T57M, F74H, and F187S and their combinations resulting in 14 variants encoded by point mutations in codons for up to four different active site residues. The mutant gene constructs were generated by using mismatch primers (Table S21). A 3 µL Kinase, Ligase & DpnI (KLD) Treatment Reaction containing 0.5 µL of the unpurified PCR product, 1x KLD Reaction Buffer and 1x KLD Enzyme Mix (New England Biolabs) was incubated at room temperature for 10 minutes followed by 10 minutes at 37°C.

### 8. Colorimetric assay for fluoride detection

A colorimetric fluoride screening method was developed here for 96-well plate format based on the agar-plate based colony screening method initially described by Davis et al. (43) Xylenol orange stock solution was prepared with a concentration of 20 mM. To prepare the zirconium oxychloride octahydrate stock solution, 0.3022 g of zirconium(IV)-oxychloride octahydrate was dissolved in 925 μL water and 1.85 mL 12.1 M HCl added and then boiled for 5 min to achieve final concentrations of 500 mM fully depolarized zirconium and 8 M HCl. After cooling, the stock solution was diluted to a concentration of 50 mM zirconium. Shortly before the experiment, the working solution was prepared with 7.8 M HCl, 400 μM xylenol orange and the complex was initiated by the addition of 250 μM zirconium. A list of potential interfering substances and buffers were tested, and results are listed in Table S11. Sodium phosphate monobasic (hereafter phosphate) interferes with the formation of the xylenol orange-fluoride complex at phosphate concentrations above 100 µM. The time dependence of phosphate interference was analyzed over a range of phosphate concentrations between 100 and 1000 µM (Fig. S28).

*E. coli* T7 expression strains carrying the corresponding plasmids were grown in TB medium in a 2 mL deep well plate (Corning, Axygen Inc P-DW-20-C-S) with 100 μg/mL spectinomycin until a optical density of 0.8 was reached The cells were induced with 0.1 mM IPTG and incubated for 24 h. The plate was centrifuged at 4000 RCF for 10 min, the supernatant was decanted and the cells were washed twice with 2 mL 20 mM HEPES (pH 8). After centrifugation at 4000 RCF for 10 min the cells were resuspended in 1 mL 20mM HEPES (pH 8) and fluoroacetate was added to a final concentration of 1mM prior to incubation at 37°C for 24 h. The cells were centrifuged at 4000 RCF for 15 min and 200 μL of the supernatant was transferred to a new 96 well plate and 40 μL of the zirconium/xylenol orange working solution was added. The final concentrations in each well were 1.3 M HCl, 66.7 μM xylenol orange, and 41.7 μM zirconium. Two independent experiments were performed. The first experiment was performed in duplicate for all protein variants and assessed for defluorination activity. The second experiment was conducted in triplicate for all protein variants and tested for defluorination but with normalization to protein concentrations. For the second experiment, protein concentrations were determined using the Pierce^TM^ 660nm Protein Assay kit (Thermo Fisher Scientific, Waltham, MA, USA) following the manufacturer’s instructions.

The absorbances used to calculate fluoride ion concentration was measured at 540 nm and the isosbestic point was measured at 480 nm using an Eon spectrophotometer (Biotek). The approximate fluoride concentration is calculated using the following formula: x=(abs(480 nm)) / (abs(540 nm)) where a ratio above 1.8 indicates a [F^-^] greater than 100 μM. Strongly positive samples (>100 μM [F^-^], x>1.8) appear yellow and negative samples appear red (<10 μM [F^-^], x<1.2). Fluoride calibration curves were prepared by spiking sodium fluoride amounts between 0 µM and 100 µM into cell supernatants taken from negative control samples. The assay shows a 2^nd^ degree polynomial relationship for fluoride concentrations between 0 μM and 100 μM. Fluoride concentrations above 100 µM were diluted 1:3 (Fig. S26) and quantified using a 1:3 diluted calibration curve. It is recommended to measure samples in a span of 15 min after adding the reagent mix, as this time span. Is shown to only slightly alter measurement data even at phosphate concentrations of 100 µM. It is advised to perform the calibration in the sample matrix (negative control) to get reliable concentration data. More information and a spreadsheet to prepare diluted and undiluted calibration curves is available https://github.com/MSM-group/gut_microbe_defluorination_paper/tree/main/Dehalogenation_colorimetric_assays/Experiment2.

### 9. Chloride detection assay

A chloride detection assay kit (QuantiChrom^TM^ Chloride Assay Kit) was obtained from Huber lab and assays were run according to the manufacturer’s instructions with the following changes: Briefly, *E. coli* T7 expression strains carrying the corresponding plasmids were grown in TB medium in a 2 mL deep well plate (Corning, Axygen Inc P-DW-20-C-S) with 100 μg/mL spectinomycin until an optical density of 0.8 was reached. The cells were induced with 0.1mM IPTG and incubated for 24 h. The plate was centrifuged at 4000 RCF for 10 min, the supernatant was discarded, and the cells were washed twice with 2 mL 20 mM HEPES (pH 8). After centrifugation at 4000 RCF for 10 min the cells were resuspended in 1 mL 20 mM HEPES (pH 8) and chloroacetate was added to a final concentration of 30 mM prior to incubation at 37°C for 24 h. The cells were centrifuged at 4000 RCF for 15 min and 2.5 μL of the supernatant was transferred to a new 96 well plate and 100 μL of the colorimetric working solution was added. The chloride concentration was measured at 610 nm with an Eon spectrophotometer (Biotek). The limit-of-quantification was 0.2 mM.Two independent experiments were performed. The first experiment was performed in duplicate for all protein variants and assessed for dechlorination activity. The second experiment was conducted in triplicate for all protein variants and tested for dechlorination but with normalization to protein concentrations.

### 10. Whole-gene alanine-scanning mutagenesis and chimeragenesis

High-throughput alanine-scanning mutagenesis was performed using the Q5® Site-Directed Mutagenesis Protocol (New England Biolabs Inc.) with the following amendments. Briefly, back-to-back forward and reverse primers were designed using NEBaseChanger (https://nebasechanger.neb.com/), where the alanine codon substitution was contained in the center of the forward mutagenic primer. Note that a histidine codon was substituted in place of any native alanine codons. Primers were ordered from Integrated DNA Technologies. The complete set of alanine (n=474) are available on https://github.com/MSM-group/biodefluorination-paper/tree/main/Alanine_scanning_primers. A 6 µL PCR reaction, containing 1x Q5 Hot Start High-Fidelity 2X Master Mix (New England Biolabs), 0.5 µM of the forward and reverse primer, and 10 ng template DNA, was carried out for each variant. P6, WP_178618037.1 (*Guopingia tenuis*) and P1 quadruple variant, WP_118709078.1 F54I-T57M-F74H-F187S (*Enterocloster aldenensis*) cloned into the pCDFDuet-1 vector was used as template DNA. PCR cycles were 98°C for 30 seconds, followed by 30 cycles of 98°C for 10 seconds, 65°C for 20 seconds and 72°C for 2 minutes, followed by a 2-minute final extension at 72°C. A 3 µl KLD Treatment Reaction containing 0.5 µl of the unpurified PCR product, 1x KLD Reaction Buffer and 1x KLD Enzyme Mix (New England Biolabs) was incubated at room temperature for 10 minutes followed by 10 minutes at 37°C. The entire reaction was transformed into NEB® DH5α competent *E. coli* using the manufacturer’s recommended transformation protocol and sequence verified on the Pacific Biosciences Revio platform (Pacific Biosciences, CA) and analyzed using custom pipelines. Plasmid DNA of sequence-verified variants were isolated and transformed into T7 Express cells (New England Biolabs), which was once again sequence verified (Pacific Biosciences). We engineered five chimeric proteins by combining longer sequences from P6, WP_1786180371.1 and P1 quad variant, WP_1187090781.1 F54I-T57M-F74H-F187S (Table S5). Gibson assembly (primers in Table S5) was used for the construction of the plasmids.

### 11. Machine learning

Random forest regression and classification was performed based on scripts and featurization methods published previously (86, 87) with the random forest algorithm as implemented in the ranger package (88). Briefly, for activity prediction using regression based on the alanine scanning data, each point variant (n=474) was featurized using the method of Atchley et al. (89) to a vector representing the position index, size, charge, identity, codon diversity, and polarity of each amino acid which was converted to alanine. These features were subsequently used to predict the change or delta in defluorination enzyme activity from wild-type as a result of each alanine substitution. For the random forest classification task, the sequences of all HAD superfamily enzymes which have been characterized in this study and previously in the literature were aligned using a protein structure-aware alignment method (90). Invariant residues in the alignment were trimmed and the remaining residues were one-hot encoded and used as features to classify HAD sequences as 1: defluorinating or 0: non-defluorinating. For both classification and regression tasks, the data were split by stratified random sampling into 80% training and 20% testing sets. Grid search was used to tune model hyperparameters by 10-fold cross validation repeated in triplicate. All forests were grown to a size of 1000 trees. Hyperparameters that were tuned during cross-validation included the number of variables randomly sampled as candidates at each split, the minimum size of terminal nodes and the splitting rule methods. The variability in training and testing set prediction performances were examined with 1000 different random seeds for training-test splits. To assess the specific importance of the C-terminal region for classification prediction, only the final 41 residues in the HAD alignment were used as features and model performance was compared to models trained on the full-length alignment. Scripts for the machine learning analysis are available at: https://github.com/MSM-group/gut_microbe_defluorination_paper/tree/main/machine_learning/.

### 12. Experimental validation of machine learning model predictions

Random forest classification models were used to predict defluorination probability for HAD homologs in NCBI RefSeq and assembled coding sequences from human gut metagenomes in the Short Read Archive (BioProject ID: PRJNA268964). Five diverse HAD sequences (denoted P20-P24) with defluorination prediction probabilities ranging between 0.38 and 0.85 (Fig. S35) were selected for experimental validation. Genes were codon optimized and synthesized as described above by IDT and cloned by Gibson Assembly into multiple cloning site 1 of pCFDuet-1 vectors. Codon optimized sequences are available here: https://github.com/MSM-group/gut_microbe_defluorination_paper/tree/main/protein_sequences_info/. Constructs were transformed into *E. coli* DH5α cells and verified by Sanger sequencing (Microsynth, Balgach, Switzerland). Sequence-verified constructs were transformed into *E. coli* T7 Express cells for protein production (New England Biolabs Frankfurt, Germany, NEB C2566I). Defluorination activity was then assessed using the colorimetric assay method described above and compared to machine learning predictions for defluorination.

### 13. In vivo validation of P2 from *Gordonibacter pamelaeae* DSM 110924

*Gordonibacter pamelaeae* was inoculated from a cryostock into 8 mL gut microbiota medium (GMM: tryptone peptone 2 g/L; yeast extract 1 g/L; raffinose 10 g/L; L-cysteine 0.5 g/L; meat extract 5 g/L; KH_2_PO_4_ 100 mM; MgSO_4_-7H_2_O 0.002 g/L; NaHCO_3_ 0.4 g/L; NaCl 0.08 g/L; CaCl_2_ 8 mg/L, menadione 1 mg/L; FeSO_4_ 0.4 mg/L ; hematin 1.2 mg/L; histidine 0.2 mM; Tween 80 0.5 mL/L; ATCC vitamin mix 10 mL/L; ATCC trace mineral mix 10 mL/L; resazurin 1 mg/L) in triplicates and incubated anaerobically for 8 days until it reached an OD of 0.1. Fluoroacetate was added to a final concentration of 10 mM and the cultures were incubated overnight. The culture was centrifuged and the supernatant used for fluoride determination before transfer of 2 mL of supernatant into a small beaker with 10 mL TISAB-II-solution (ammonium acetate 77.1 g/L; Titriplex IV 4 g/L; ammonium chloride 26.75 g/L; acetic acid 96% 59 mL/L) and filled up to 40 mL. The fluoride concentration was measured by using an ISE (fluoride-ion sensitive electrode Orion type No. 9-09-00 and a reference electrode Orion type No. 90-02-00) by adding stepwise a 10 mg/L fluoride standard. The values obtained in mV were used for fluoride quantification (Table S4).

### 14. Steady-state kinetics experiments

Steady-state kinetic parameters for P3 were measured at a protein concentration of 0.5 mg/mL. The substrate fluoroacetate was added to final concentration of 0.4 mM; 0.8 mM; 1.6 mM; and 3.0 mM in 100mM Tris-SO_4_ (pH 8.5) and 20 mM; 50 mM; 100 mM in 300mM Tris-SO_4_ (pH 8.5). The reaction mixtures were incubated at 37 °C for the duration of the reaction period. Samples were quenched with acetonitrile at timepoints: 0 min, 10 min, 20 min, 40 min, 50 min, 60 min, 75 min, 90 min, 105 min, 120 min, 6h for the sample containing 0.4 mM-3 mM fluoroacetate and 0 min, 10 min, 30min, 60 min 90 min and 120 min for the samples containing 20-100 mM fluoroacetate. Following quenching, samples were centrifuged at 15000 RCF for 5 min and the supernatant was transferred into plastic vials for subsequent fluoride concentrations using identical methods as described in the above IC section. For each substrate concentration, the slope was calculated over the linear range of this curve, giving the velocity of product formation (mM fluoride/min). All the experiments were performed with three independent biological replicates and three technical replicates per biological replicate and per substrate concentration. Michaelis-Menten equations were used to calculate the kinetic parameters *K*_m_, *k*_cat_, and *k*_cat_*/K*_m_. A non-linear regression analysis was used to fit the data to the Michaelis–Menten equation using the R package “nlstools” (91) with 1000 non-parametric bootstrap resamplings to estimate the values and standard errors for the kinetic parameters (Table S6).

## Notes

https://github.com/MSM-group/gut_microbe_defluorination_paper/

### Competing Interest Statement

The authors have declared no competing interest.

### Summary of Updates

Refined significance: We have adjusted the Significance Statement and Introduction to better emphasize the advances in understanding enzymatic defluorination as a resource for enzyme discovery, while de-emphasizing the gut microbiome therapeutics aspect. Clarification of inaccuracies: We have added sentences on the conformational heterogeneity and limitations of the MD simulations. We have also corrected inaccuracies regarding previous work on fluorinated inhibitors and we have updated the terminology (e.g., mechanism-based inactivation rather than suicide inhibition).

