## Supplemental files for "Enzymatic carbon-fluorine bond cleavage by human gut microbes"

### Supplementary Figures

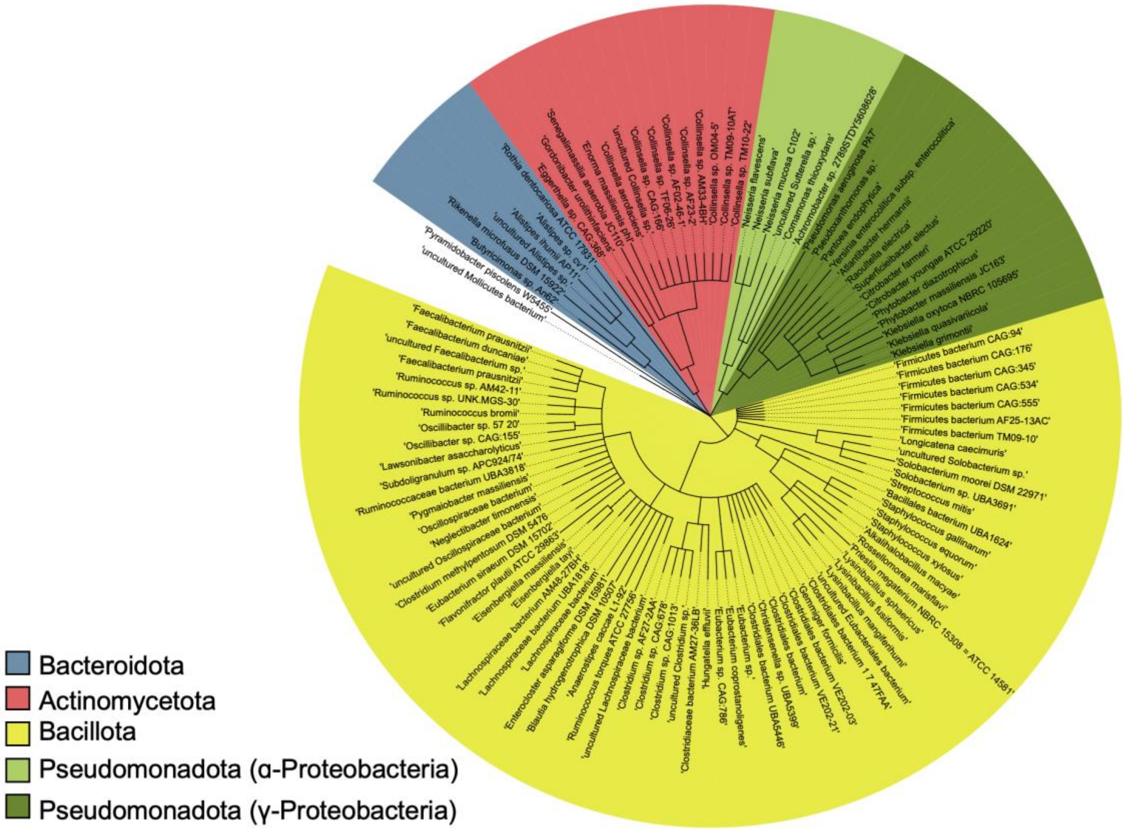

**Figure S1: Phylogenetic tree** of human gut microbial strains encoding homologs with a bitscore > 270 to the query haloacid dehalogenase from *Rhodococcus jostii* RHA1 (PDB ID: 3UMG).

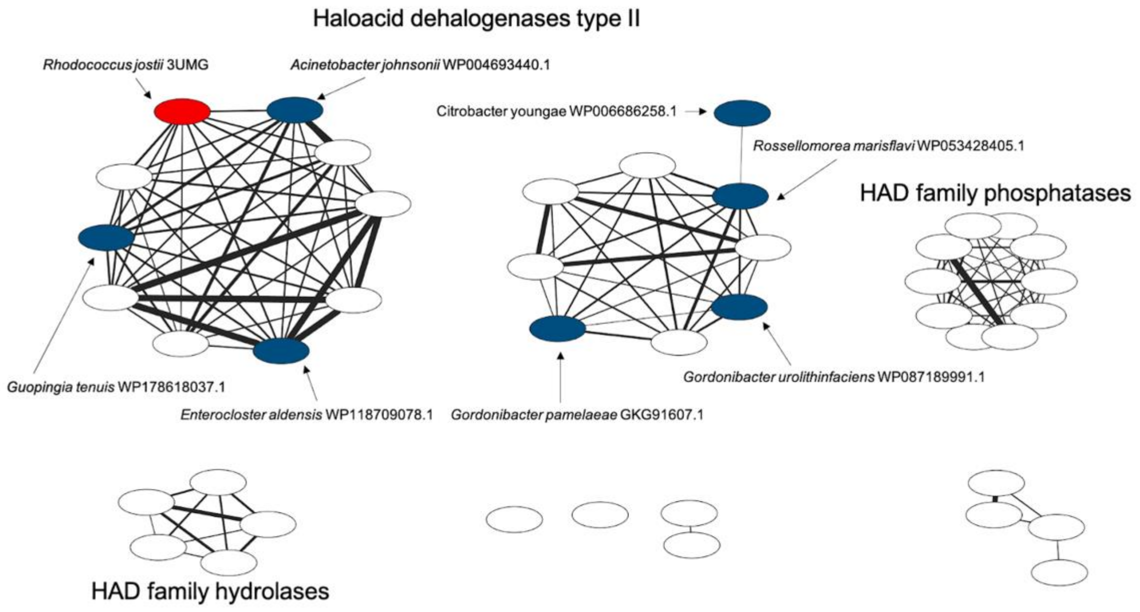

**Figure S2: Protein sequence similarity network** constructed from 60 protein sequences sharing homology to the 2-haloacid dehalogenases identified in the human gut microbiome. EFI-EST annotations of protein families are used. The width of interconnecting edges indicate the degree of relatedness between two nodes, with connections up to a raw distance of 0.95 retained. Red: Query protein sequence (known defluorinating haloacid dehalogenase) used for the initial homology search in the Humgut database. Blue: protein candidates picked for further in vitro experiments.

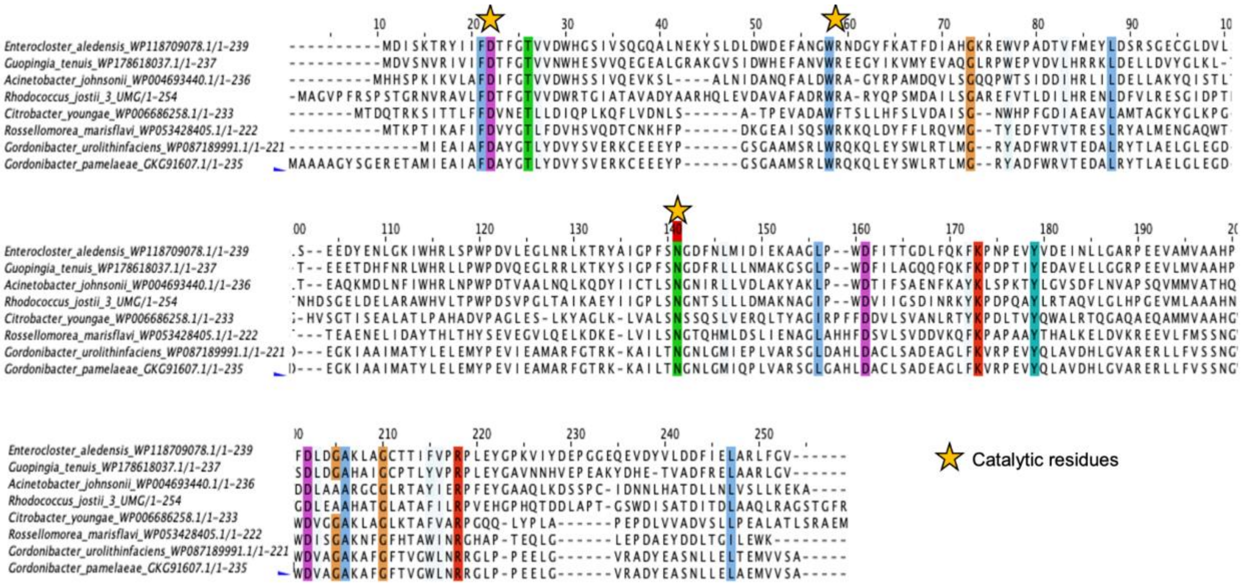

**Figure S3: Multiple sequence alignment** using Clustal Omega of eight proteins chosen for further in vitro experiments. Amino acid conservation of <95% are indicated by color. The three catalytic amino acids are marked with a star.

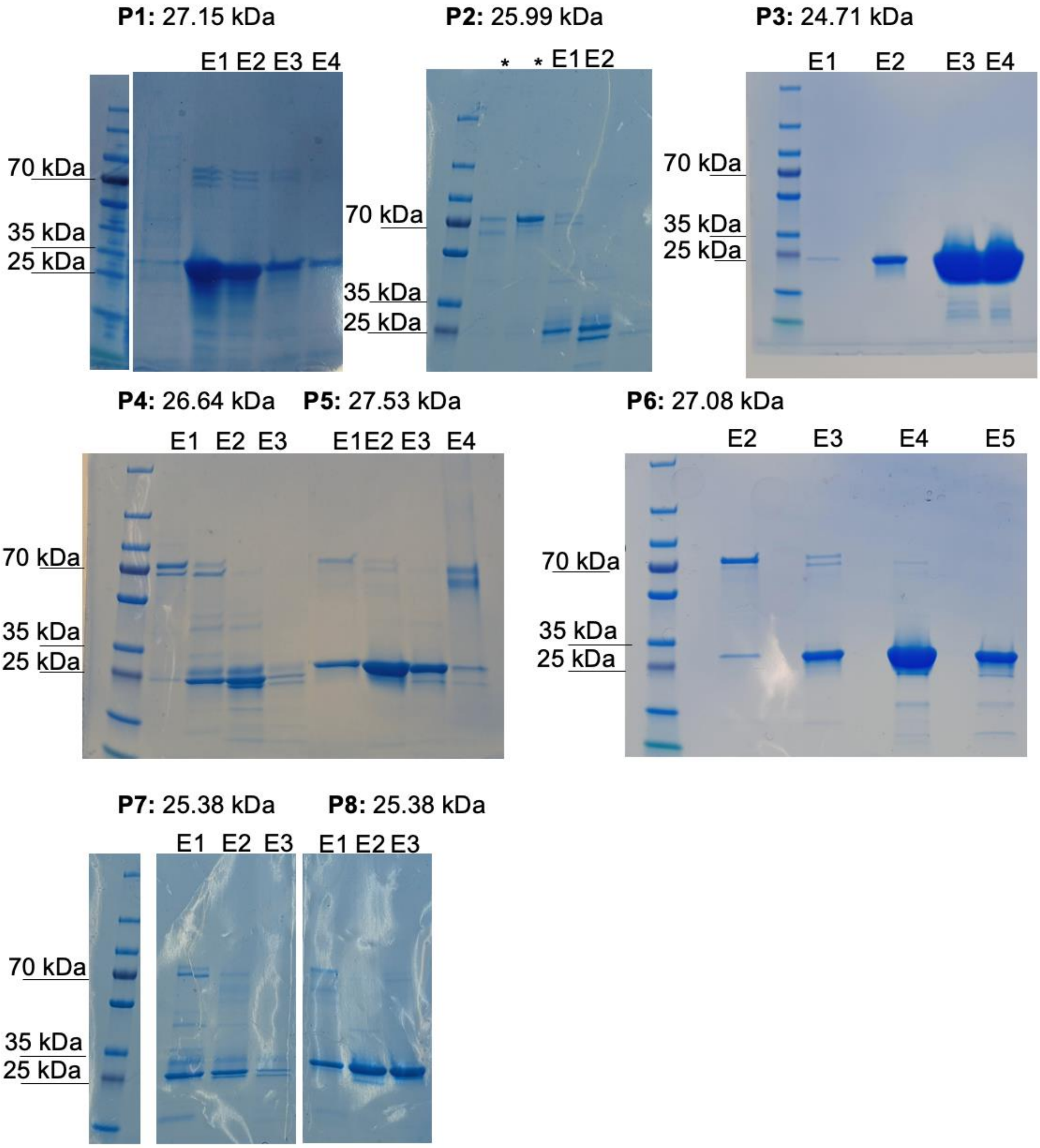

**Figure S4: SDS-PAGE gels** of the purified proteins P1-P8. *Asterisk indicates proteins not related to P1-P8. Expected sizes in kilodalton (kDa) for each protein are shown relative to the 180 kDa ladder (Thermo Scientific™ PageRuler™).

Protein 1: WP_118709078.1, *Enterocloster aldenensis*

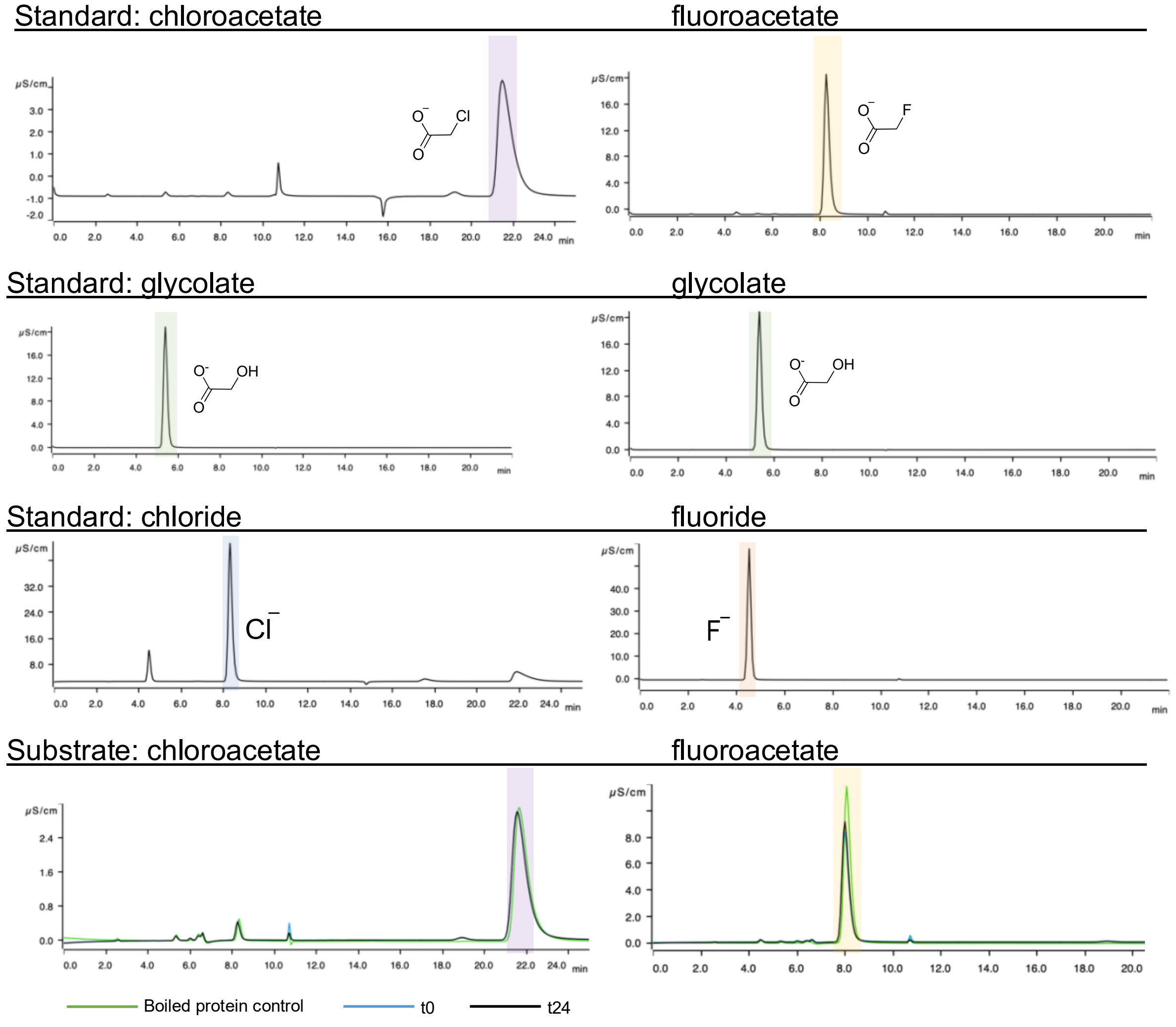

| Protein | substrate | % conversion | concentration glycolate [mM] |
|---|---|---|---|
| P1 | chloroacetate 2mM | <LOD | <LOD |
| P1 | fluoroacetate 2mM | <LOD | <LOD |

**Figure S5: Ion chromatography** traces of protein P1 (*Enterocloster aldenensis)* with two different substrates.
Green line: Boiled protein control; Blue line: t0h; Black line: t24h. The first three rows show standards. Chloroacetate shaded in purple, fluoroacetate shaded in yellow, glycolate shaded in green, chloride shaded in blue, and fluoride shaded in red. The last row represents the samples incubated with the substrates indicated. The table shows the conversion and measured glycolate concentrations. Glycolate is used for this calculation due to possible interfering chloride contaminations.

Protein 2: GKG91607.1, *Gordonibacter pamelaeae*

Standard: chloroacetate

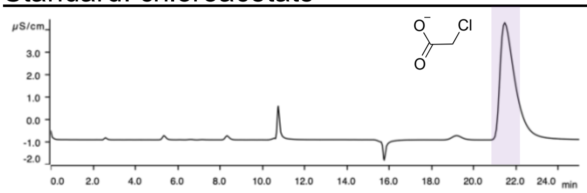

fluoroacetate

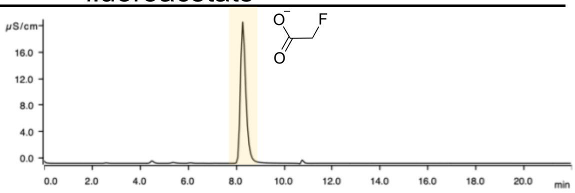

Standard: glycolate

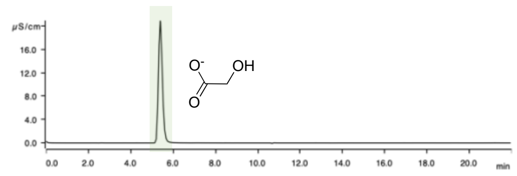

glycolate

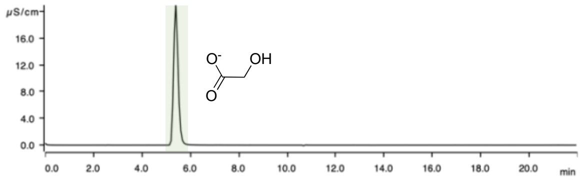

Standard: chloride

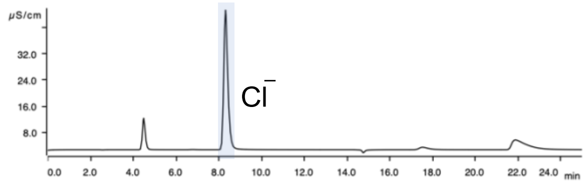

fluoride

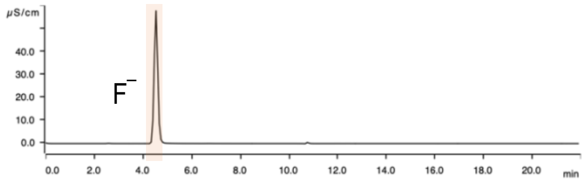

Substrate: chloroacetate

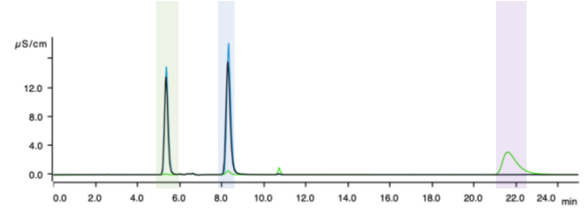

fluoroacetate

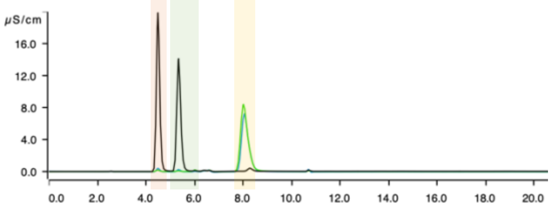

Boiled protein control  t0  t24

| Protein | substrate | % conversion | concentration glycolate [mM] |
|---|---|---|---|
| P2 | chloroacetate 2 mM | 117 | 2.34 |
| P2 | fluoroacetate 2.18 mM | 112 | 2.46 |

Substrate: 2-fluoropropionate

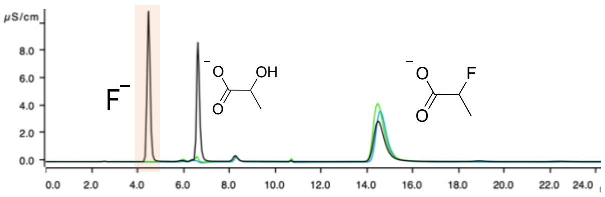

2,2-difluoroacetate

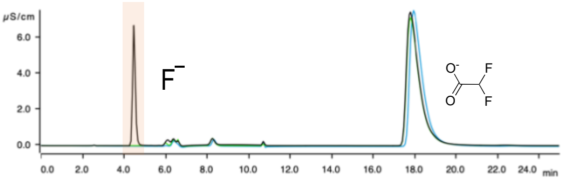

Boiled protein control  t0  t24

| Protein | substrate | % conversion | concentration fluoride [mM] |
|---|---|---|---|
| P2 | fluoropropionate 1.7 mM | 68.9 | 1.17 |
| P2 | difluoroacetate 3.7 mM | 10.85 | 0.7 |

**Figure S6: Ion chromatography** traces of protein P2 (*Gordonibacter pamelaeae*) with four different substrates.

Green line: Boiled protein control; Blue line: t0h; Black line: t24h. The first three rowsshow standards. Chloroacetate shaded in purple, fluoroacetate shaded in yellow, glycolate shaded in green, chloride shaded in blue, and fluoride shaded in red. The last rows represent the samples incubated with the substrates indicated. The table shows the conversion and measured glycolate concentrations. Glycolate is used for this calculation due to possible interfering chloride contaminations. Conversion values >100% are due to instrument variability and slight evaporation which could not be fully avoided.

Protein 3: WP_087189991.1, *Gordonibacter urolithinfaciens*

Standard: chloroacetate | fluoroacetate

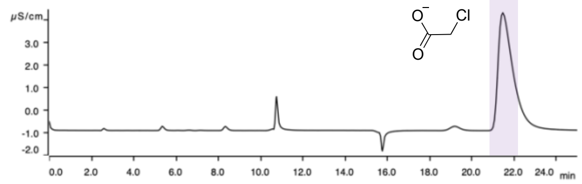

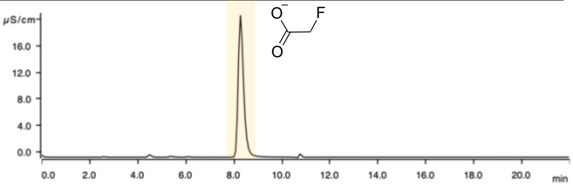

Standard: glycolate | glycolate

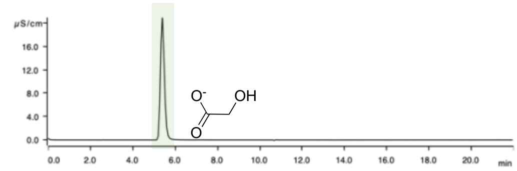

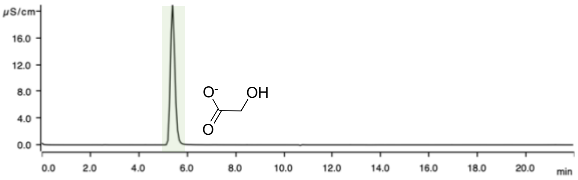

Standard: chloride | fluoride

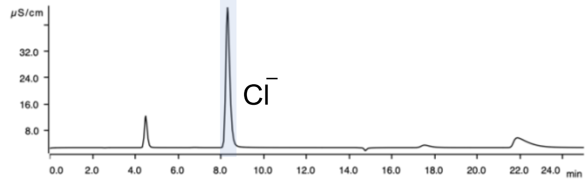

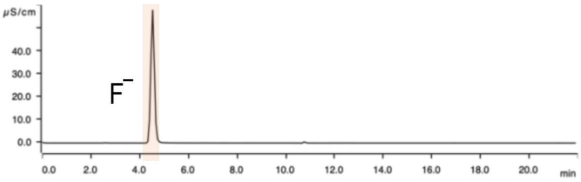

Substrate: chloroacetate | fluoroacetate

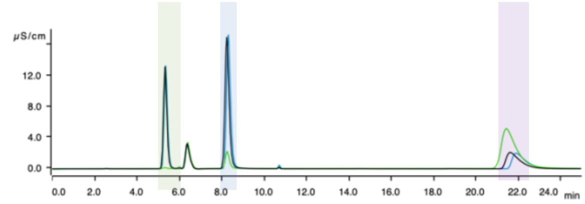

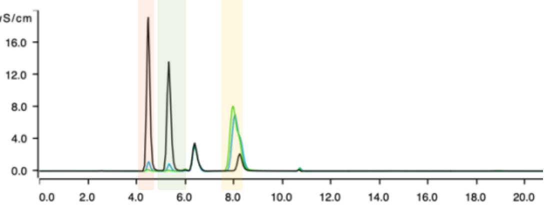

Boiled protein control — t0 — t24

| Protein | substrate | % conversion | concentration glycolate [mM] |
|---|---|---|---|
| P3 | chloroacetate 3.4 mM | 69.6 | 2.36 |
| P3 | fluoroacetate 2.4 mM | 103 | 2.45 |

Substrate: 2-fluoropropionate | 2,2-difluoroacetate

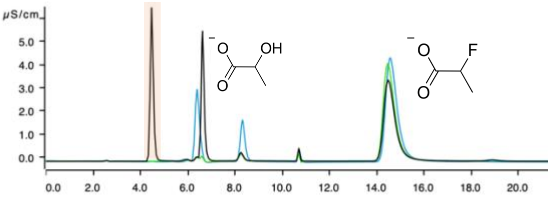

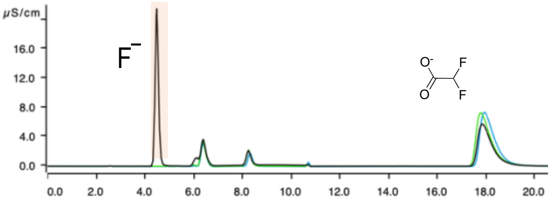

Boiled protein control — t0 — t24

| Protein | substrate | % conversion | concentration fluoride [mM] |
|---|---|---|---|

| | | | |
|---|---|---|---|
| P3 | fluoropropionate 1.7 mM | 45.5 | 0.77 |
| P3 | difluoroacetate 3.8 mM | 35 | 2.63 |

**Figure S7: Ion chromatography** traces of protein P3 (*Gordonibacter urolithinfaciens*) with four different substrates. Green line: Boiled protein control; Blue line: t0h; Black line: t24h. The first three rows show standards. Chloroacetate shaded in purple, fluoroacetate shaded in yellow, glycolate shaded in green, chloride shaded in blue, and fluoride shaded in red. The last rows represent the samples incubated with the substrates indicated. The tables show the conversion and measured glycolate concentrations. Glycolate is used for this calculation due to possible interfering chloride contaminations. Conversion values >100% are due to instrument variability and slight evaporation which could not be fully avoided.

Protein 4: WP_004693440.1, *Acinetobacter johnsonii*

Standard: chloroacetate fluoroacetate

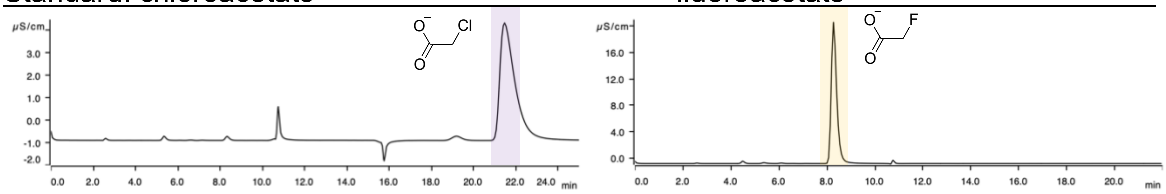

Standard: glycolate glycolate

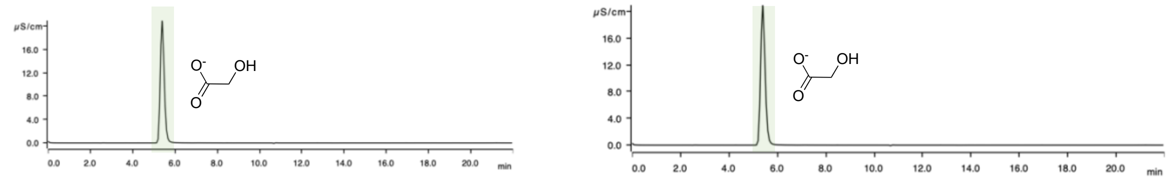

Standard: chloride fluoride

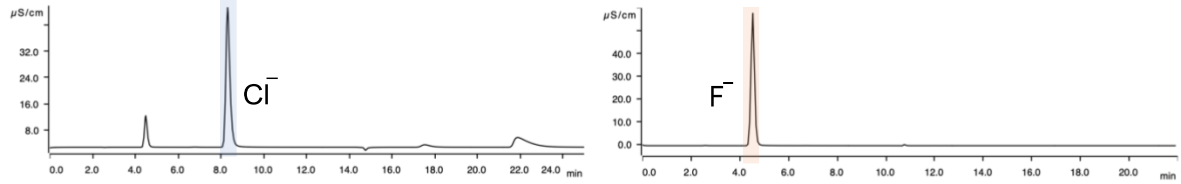

Substrate: chloroacetate fluoroacetate

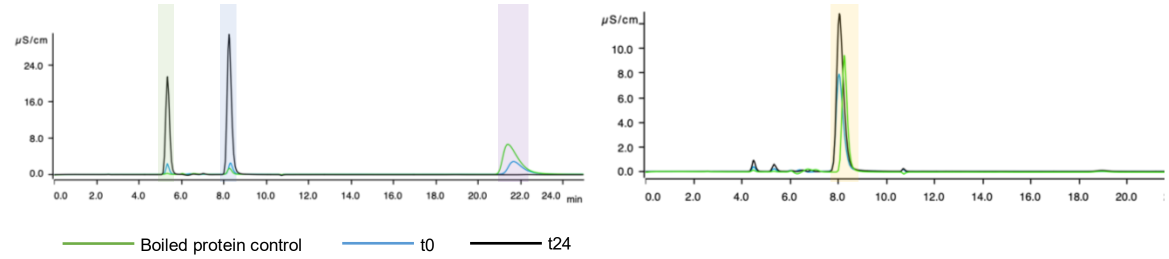

| Protein | substrate | % conversion | concentration glycolate [mM] |
|---|---|---|---|
| P4 | chloroacetate 4.4 mM | 105 | 4.62 |
| P4 | fluoroacetate 2.57 mM | 26.2 | 0.674 |

**Figure S8: Ion chromatography** traces of protein P4 (*Acinetobacter johnsonii*) with two substrates. Green line: Boiled protein control; Blue line: t0h; Black line: t24h. The first three rows show standards. Chloroacetate shaded in purple, fluoroacetate shaded in yellow, glycolate shaded in green, chloride shaded in blue, and fluoride shaded in red. The last row represents the samples incubated with the substrates indicate. The table shows the conversion and measured glycolate concentrations. Glycolate is used for this calculation due to possible interfering chloride contaminations. Conversion values >100% are due to instrument variability and slight evaporation which could not be fully avoided.

### Protein 5: 3UMG_1, *Rhodococcus jostii*

Standard: chloroacetate

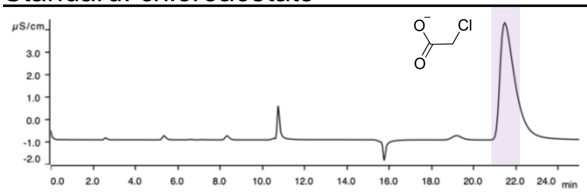

fluoroacetate

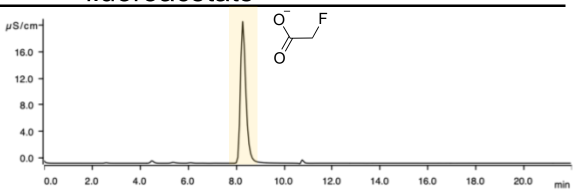

Standard: glycolate

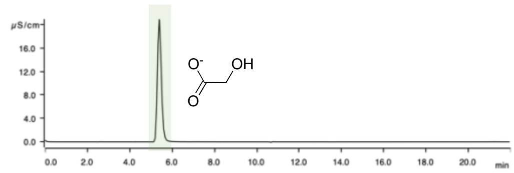

glycolate

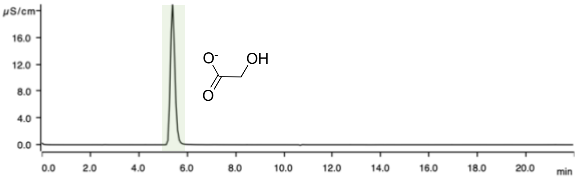

Standard: chloride

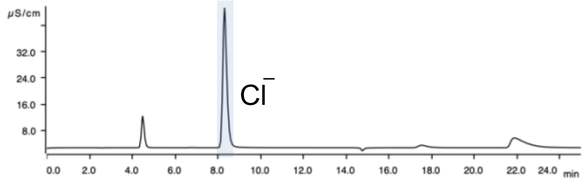

fluoride

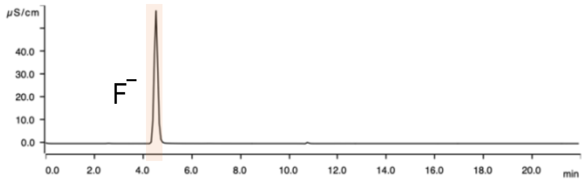

Substrate: chloroacetate

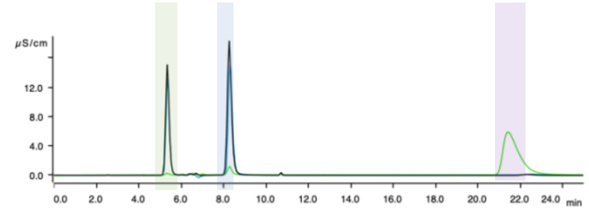

fluoroacetate

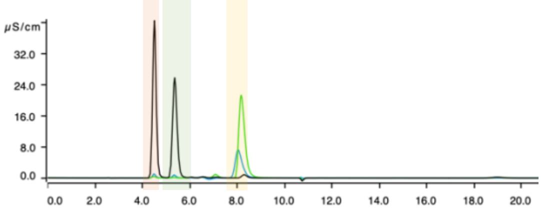

Boiled protein control — t0 — t24

| Protein | substrate | % conversion | concentration glycolate [mM] |
|---|---|---|---|
| P5 | chloroacetate 3.9 mM | 82.7 | 3.22 |
| P5 | fluoroacetate 4.7 mM | 79.8 | 3.78 |

Substrate: 2-fluoropropionate

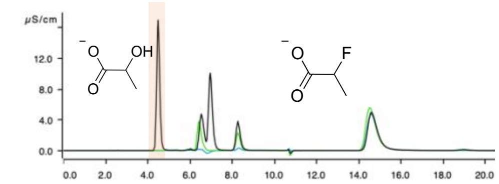

2,2-difluoroacetate

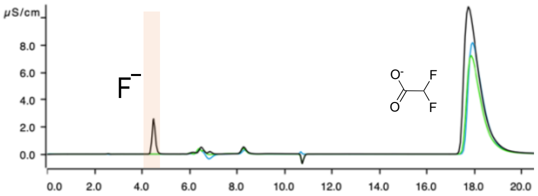

Boiled protein control — t0 — t24

| Protein | substrate | % conversion | concentration fluoride [mM] |
|---|---|---|---|
| P5 | fluoropropionate 2.35 mM | 68.3 | 1.6 |
| P5 | difluoroacetate 3.75 mM | 5.22 | 0.357 |

**Figure S9: Ion chromatography** traces of protein P5 (*Rhodococcus jostii* RHA1) with four different substrates. Green line: Boiled protein control; Blue line: t0h; Black line: t24h. The first three row show standards. Chloroacetate shaded in purple, fluoroacetate shaded in yellow, glycolate shaded in green, chloride shaded in blue, and fluoride shaded in red. The last rows represent the samples incubated with the substrates indicated. The table shows the conversion and measured glycolate concentrations. Glycolate is used for this calculation due to possible interfering chloride contaminations.

Protein 6: WP_178618037.1, *Guopingia tenuis*

Standard: chloroacetate

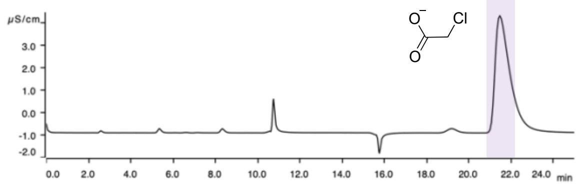

fluoroacetate

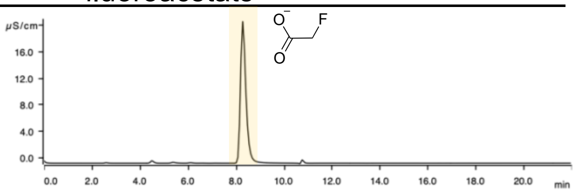

Standard: glycolate

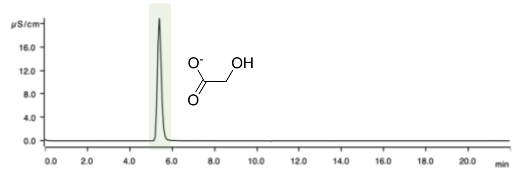

glycolate

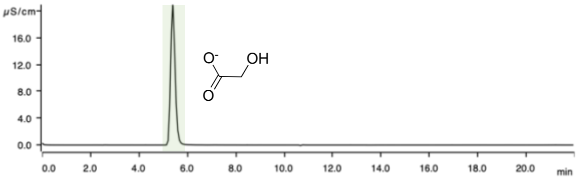

Standard: chloride

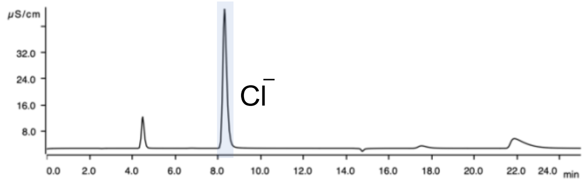

fluoride

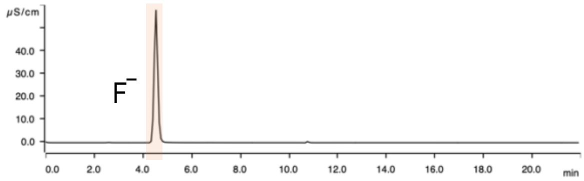

Substrate: chloroacetate

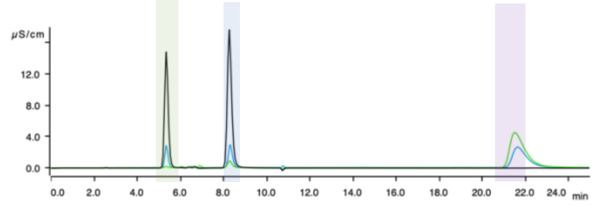

fluoroacetate

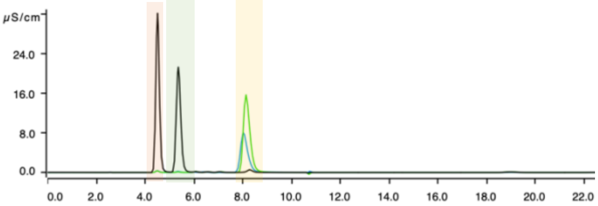

Boiled protein control t0 t24

| Protein | substrate | % conversion | concentration glycolate [mM] |
|---|---|---|---|
| P6 | chloroacetate 2.9 mM | 121 | 3.35 |
| P6 | fluoroacetate 3.4 mM | 101 | 3.70 |

Substrate: fluoro-beta-alanine

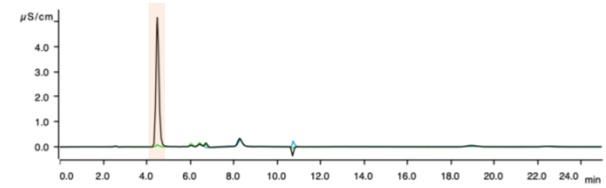

Boiled protein control t0 t24

| Protein | substrate | % conversion | concentration fluoride [mM] |
|---|---|---|---|
| P6 | Fluoro-beta-alanine 2 mM | 36.06 | 0.72 |

**Figure S10: Ion chromatography** traces of protein P6 (*Guopingia tenuis)* with different three substrates. Green line: Boiled protein control; Blue line: t0h; Black line: t24h. The first three rows show standards. Chloroacetate shaded in purple, fluoroacetate shaded in yellow, glycolate shaded in green, chloride shaded in blue and fluoride shaded in red. The last rows represent the samples incubated with the substrates indicated. The tables show the conversion and measured glycolate concentrations. Glycolate is used for this calculation due to possible interfering chloride contaminations. Conversion values >100% are due to instrument variability and slight evaporation which could not be fully avoided.

Protein 7: WP_006686258.1, *Citrobacter youngae*

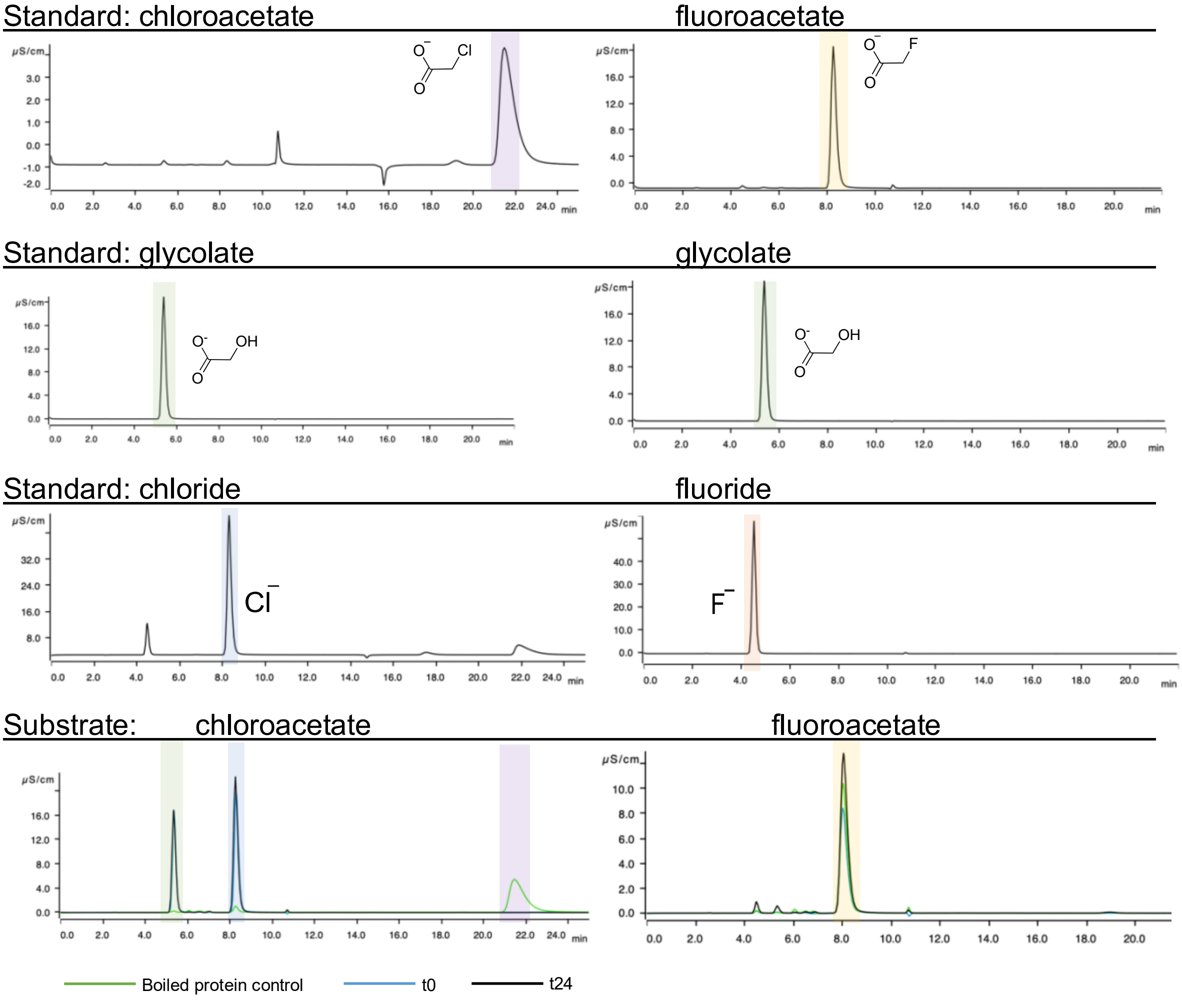

| Protein | substrate | % conversion | concentration glycolate [mM] |
|---|---|---|---|
| P7 | chloroacetate 3.5 mM | 92.6 | 3.24 |
| P7 | fluoroacetate 2.7 mM | 4.77 | 0.126 |

**Figure S11: Ion chromatography** traces of protein WP_006686258.1 (*Citrobacter youngae*) with two different substrates. Green line: Boiled protein control; Blue line: t0h; Black line: t24h. The first three rows show standards. Chloroacetate shaded in purple, fluoroacetate shaded in yellow, glycolate shaded in green, chloride shaded in blue, and fluoride shaded in red. The last row represent the samples incubated with the substrates indicated. The table shows the conversion and measured glycolate concentrations. Glycolate is used for this calculation due to possible interfering chloride contaminations.

Protein 8: WP_053428405.1, *Rossellomorea marisflavi*

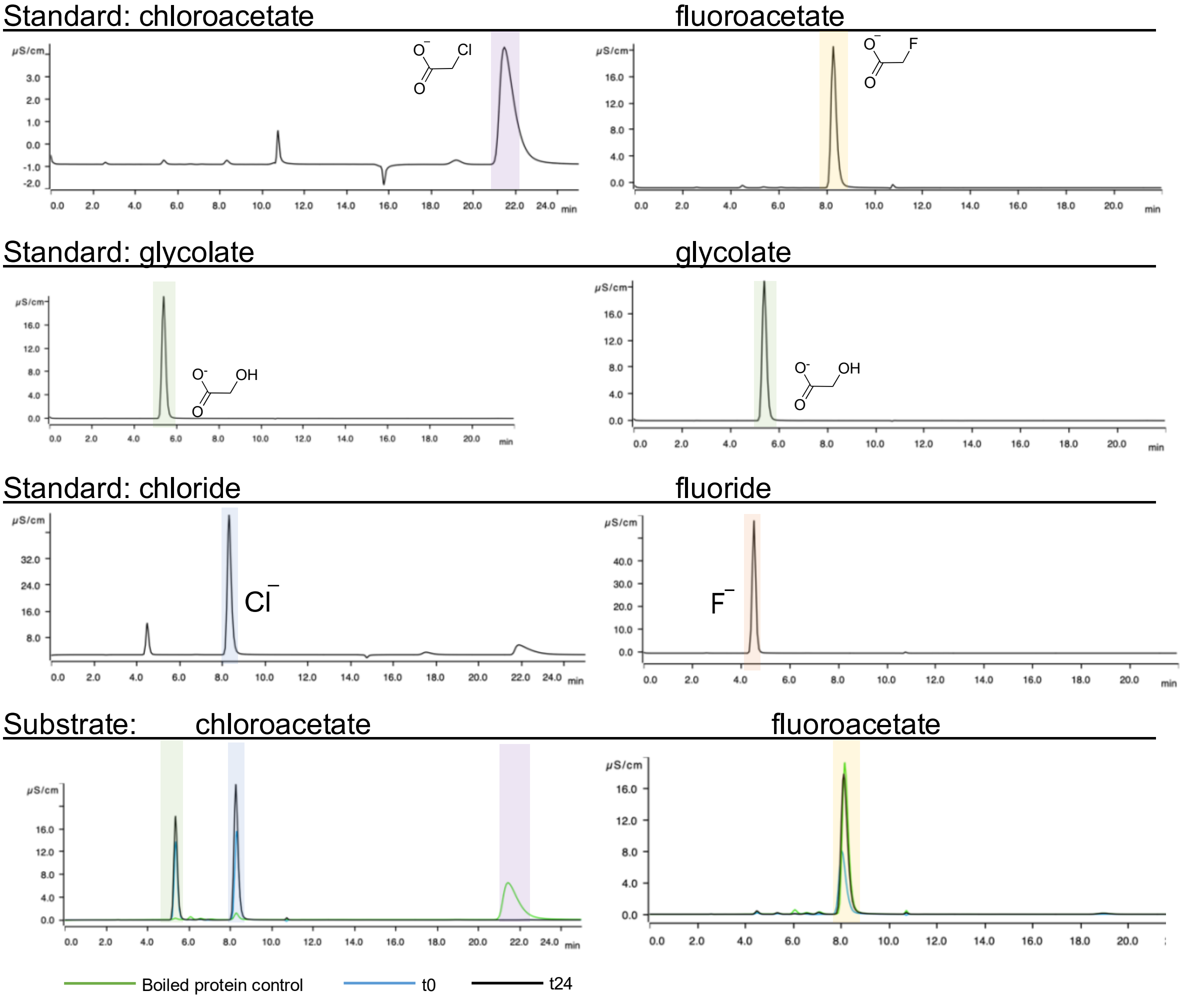

| Protein | substrate | % conversion | concentration glycolate [mM] |
|---|---|---|---|
| P8 | chloroacetate 4.3 mM | 89.5 | 3.84 |
| P8 | fluoroacetate 4.3 mM | 0.8 | 0.03 |

**Figure S12: Ion chromatography traces** of protein P8 (*Rossellomorea marisflavi)* with two different substrates. Green line: Boiled protein control; Blue line: t0h; Black line: t24h. The first three rows show standards. Chloroacetate shaded in purple, fluoroacetate shaded in yellow, glycolate shaded in green, chloride shaded in blue, and fluoride shaded in red. The last row represent the samples incubated with the substrates indicated. The table shows the conversion and measured glycolate concentrations. Glycolate is used for this calculation due to possible interfering chloride contaminations.

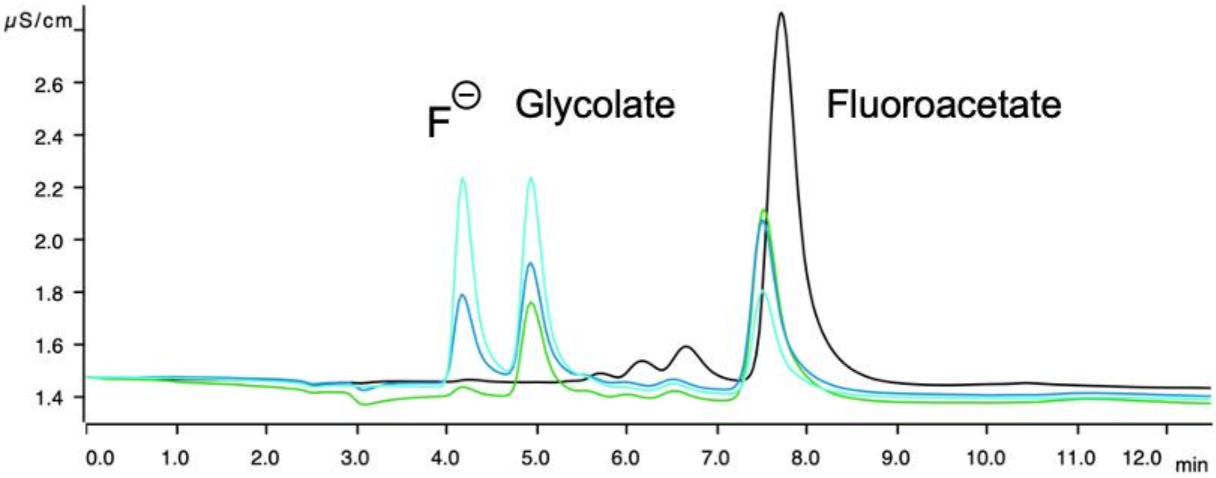

**Figure S13: Ion chromatography** traces of protein P3 (*Gordonibacter urolithinfaciens*) incubated anaerobically with fluoroacetate. Black: control; Green t1h; Blue: t3h; ligth blue: t20h.

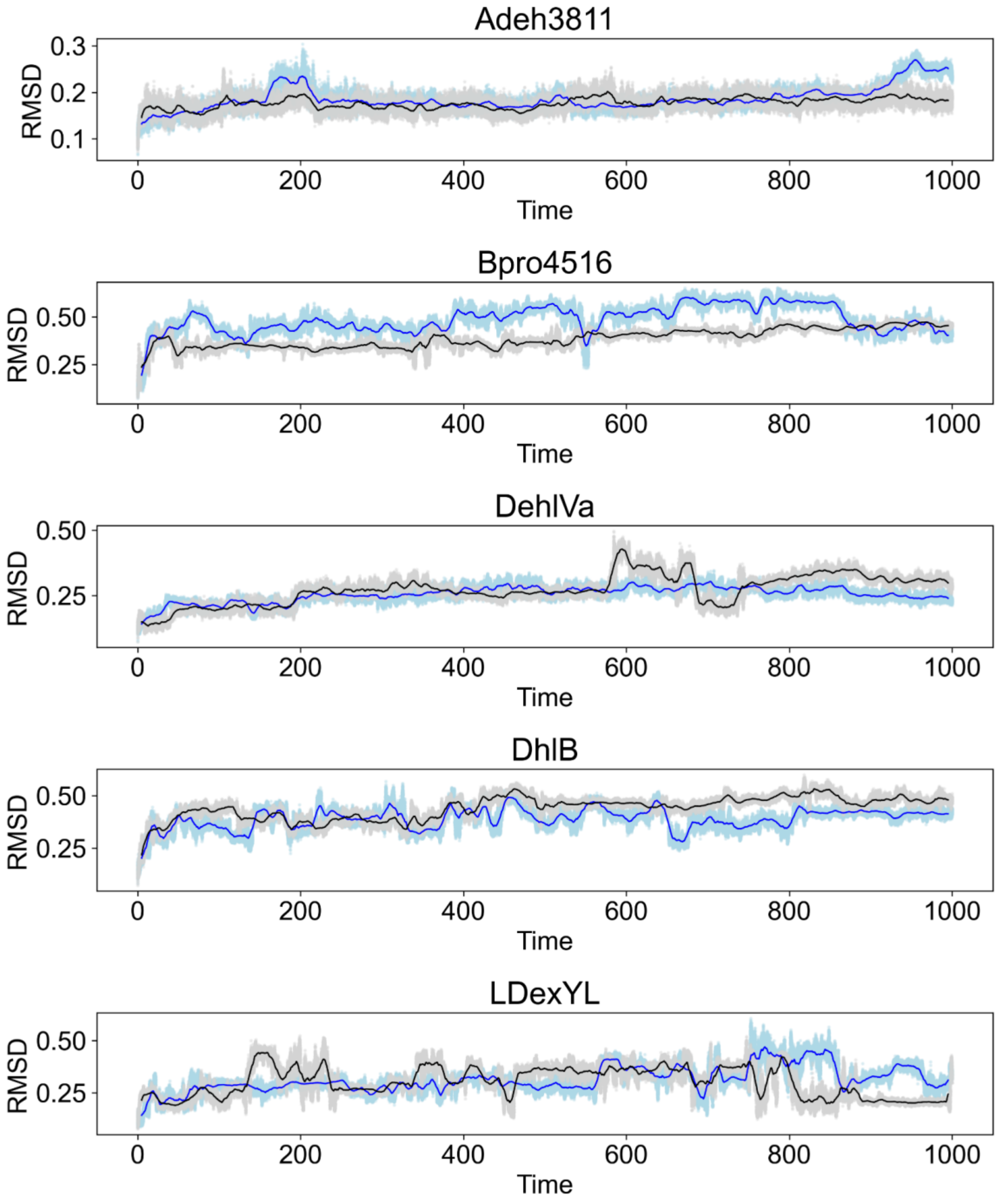
Adeh3811
RMSD
0.1
0.2
0.3
0
200
400
600
800
1000
Time
Bpro4516
RMSD
0.25
0.50
Time
DehIVa
RMSD
0.25
0.50
Time
DhlB
RMSD
0.25
0.50
Time
LDexYL
RMSD
0.25
0.50
Time

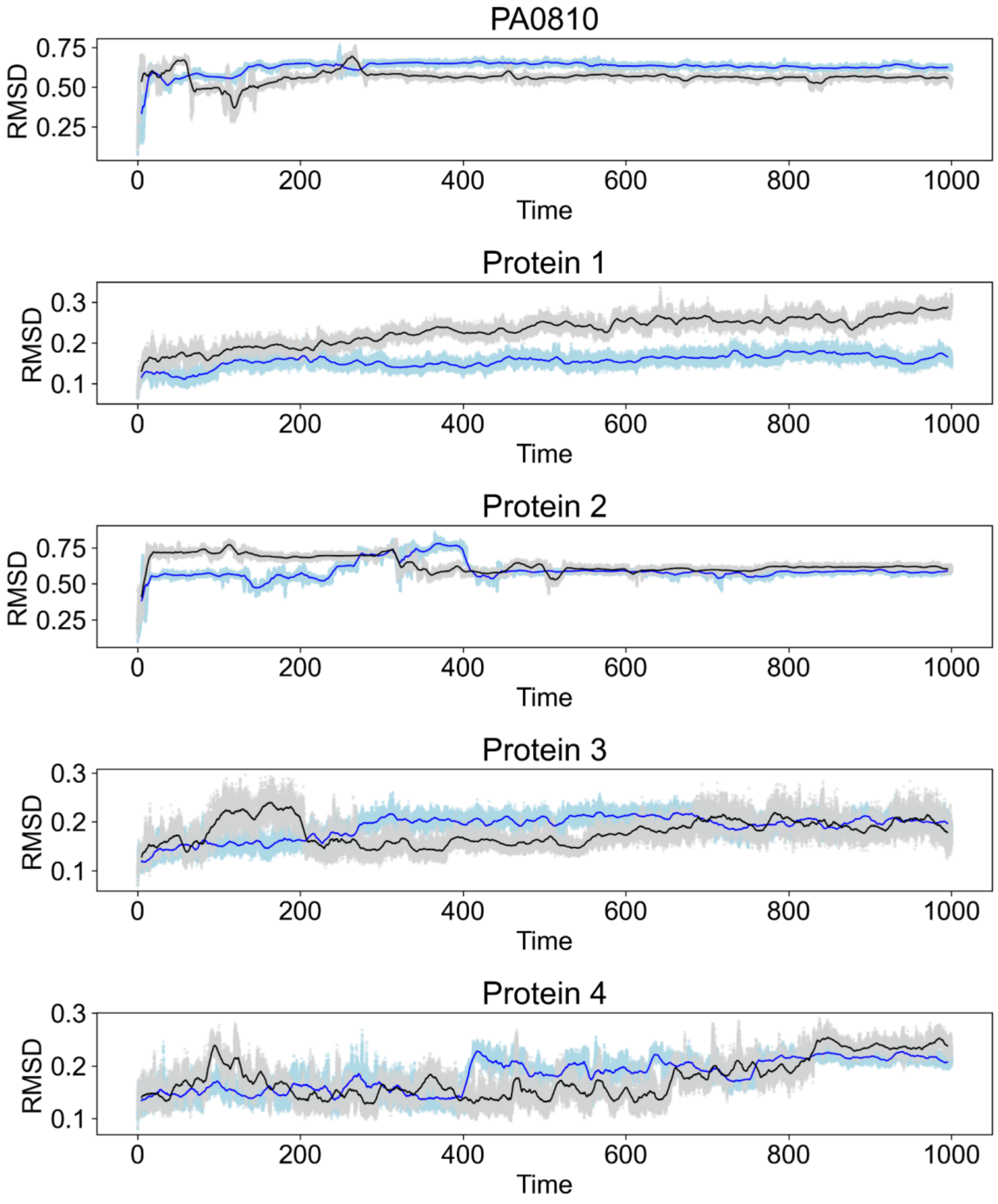
PA0810
RMSD
0.75
0.50
0.25
0
200
400
600
800
1000
Time
Protein 1
RMSD
0.3
0.2
0.1
Time
Protein 2
RMSD
0.75
0.50
0.25
Time
Protein 3
RMSD
0.3
0.2
0.1
Time
Protein 4
RMSD
0.3
0.2
0.1
Time

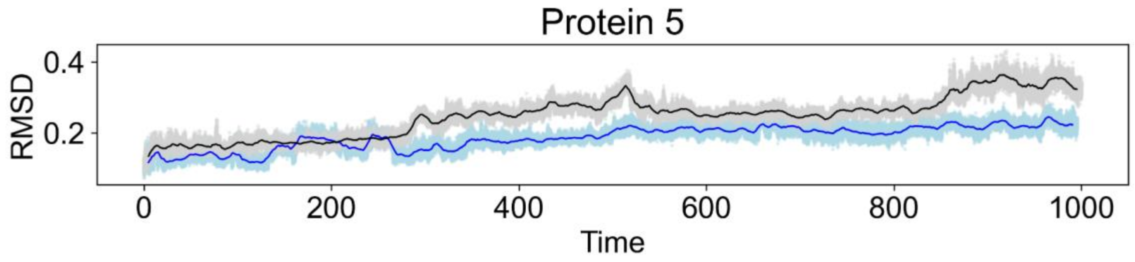
Protein 5
RMSD
0.4
0.2
0
200
400
600
800
1000
Time

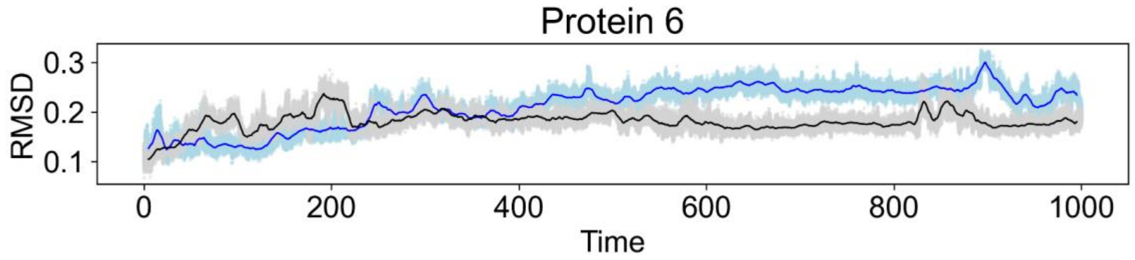
Protein 6
RMSD
0.3
0.2
0.1
0
200
400
600
800
1000
Time

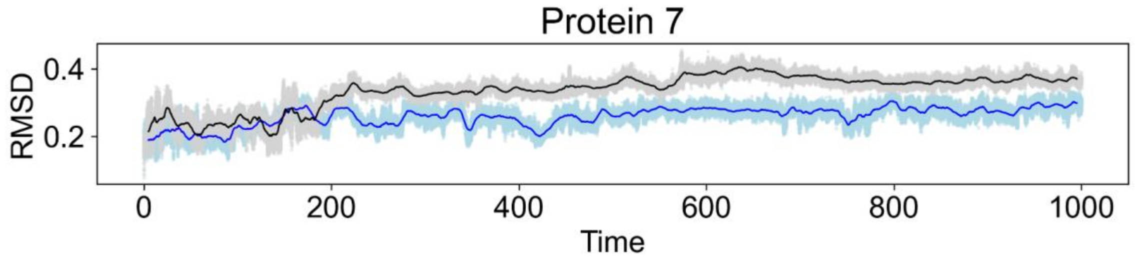
Protein 7
RMSD
0.4
0.2
0
200
400
600
800
1000
Time

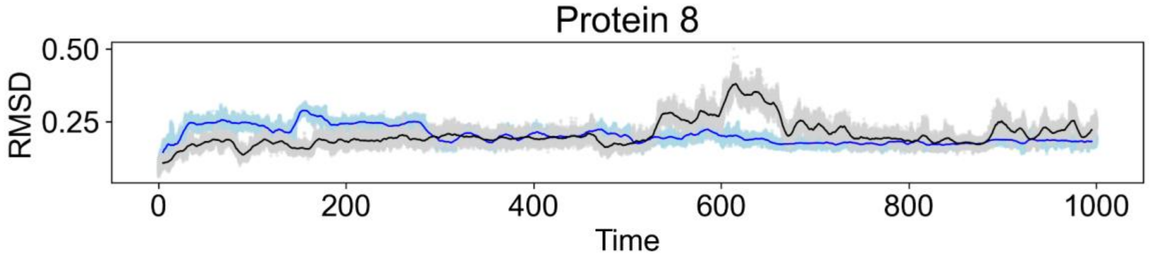
Protein 8
RMSD
0.50
0.25
0
200
400
600
800
1000
Time

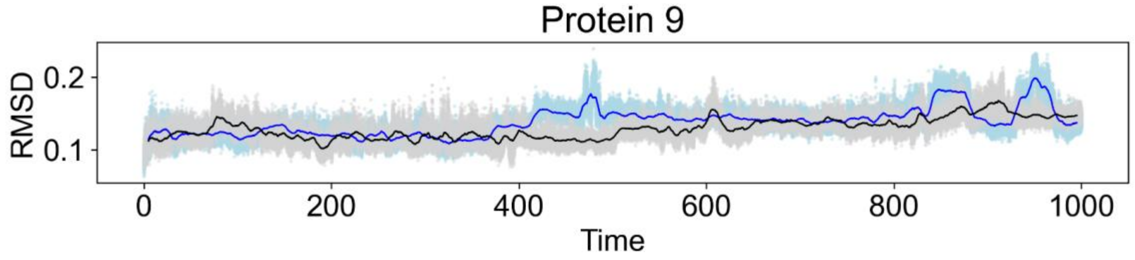
Protein 9
RMSD
0.2
0.1
0
200
400
600
800
1000
Time

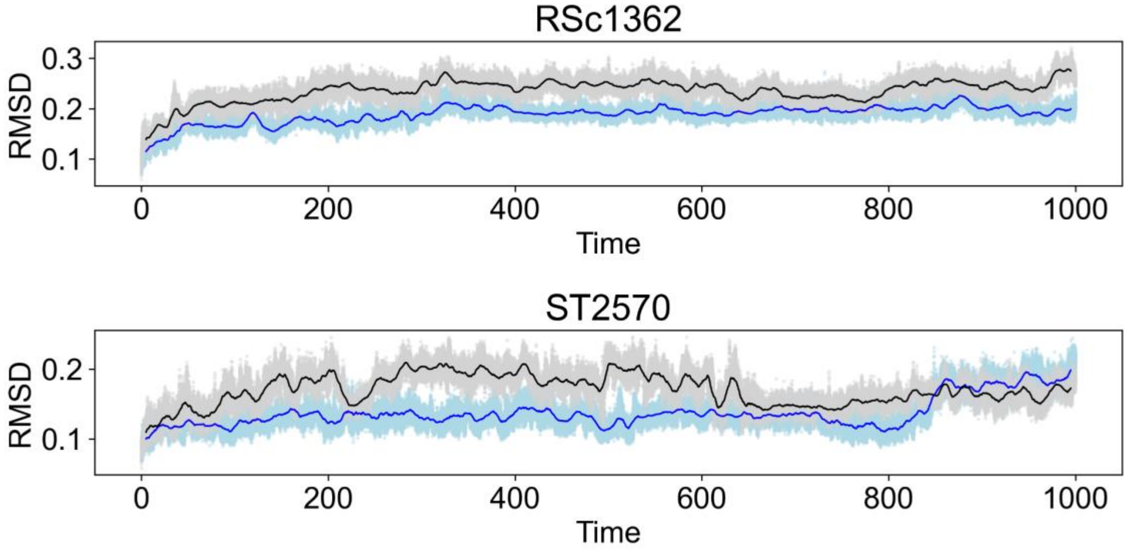

**Figure S14: Root Mean Square Deviation (RMSD, in nm) files from molecular dynamic simulations.** Two independent replicates with rolling averages of 1 ns are indicated as black and blue lines and standard deviations of the 1 ns window are represented by the gray/light blue shading. X-axis in ns, full simulation time 1μs. Black: replicate 1; Blue: replicate 2;

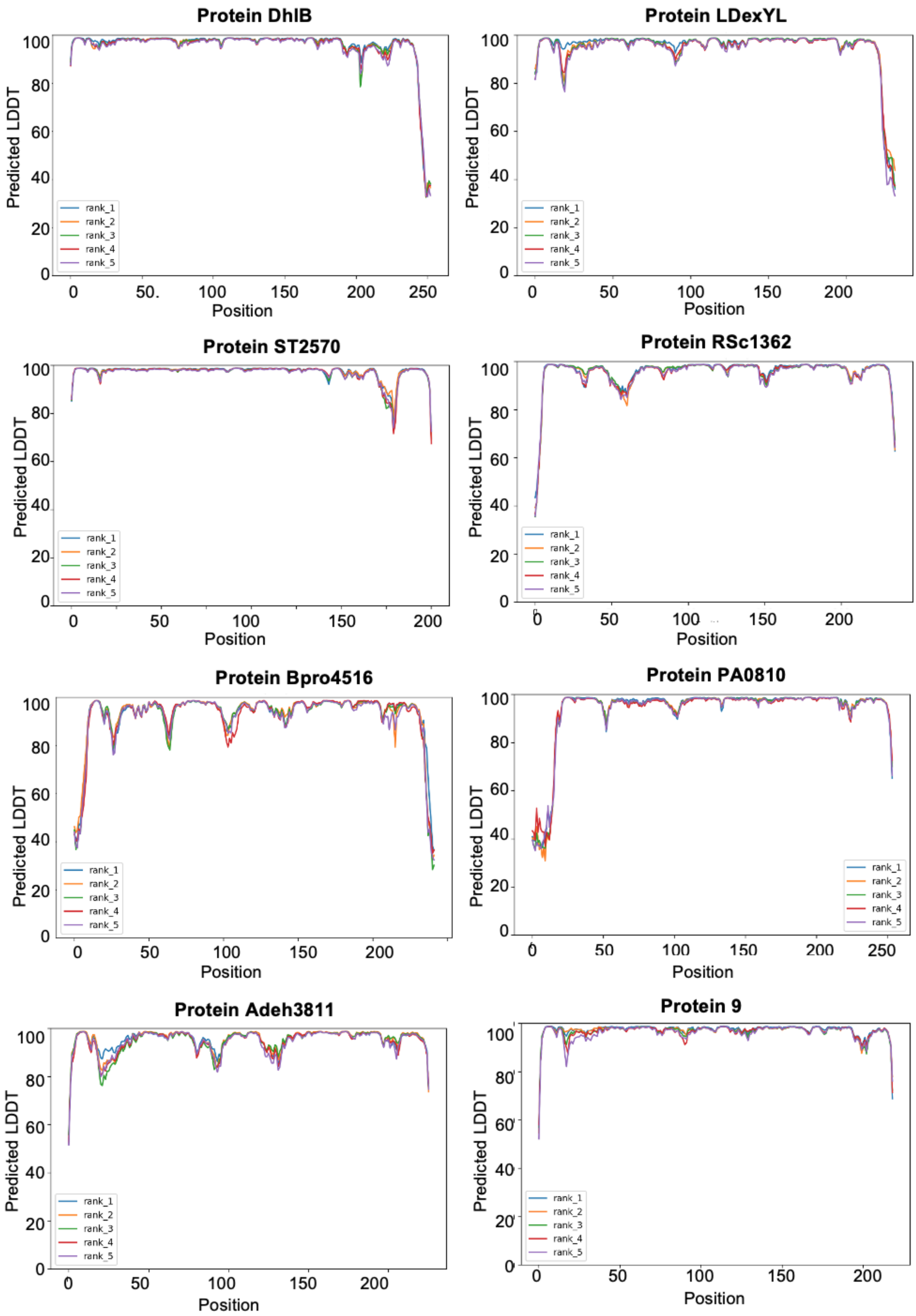
Protein DhlB
Predicted LDDT
Position
rank_1
rank_2
rank_3
rank_4
rank_5
Protein LDexYL
Protein ST2570
Protein RSc1362
Protein Bpro4516
Protein PA0810
Protein Adeh3811
Protein 9

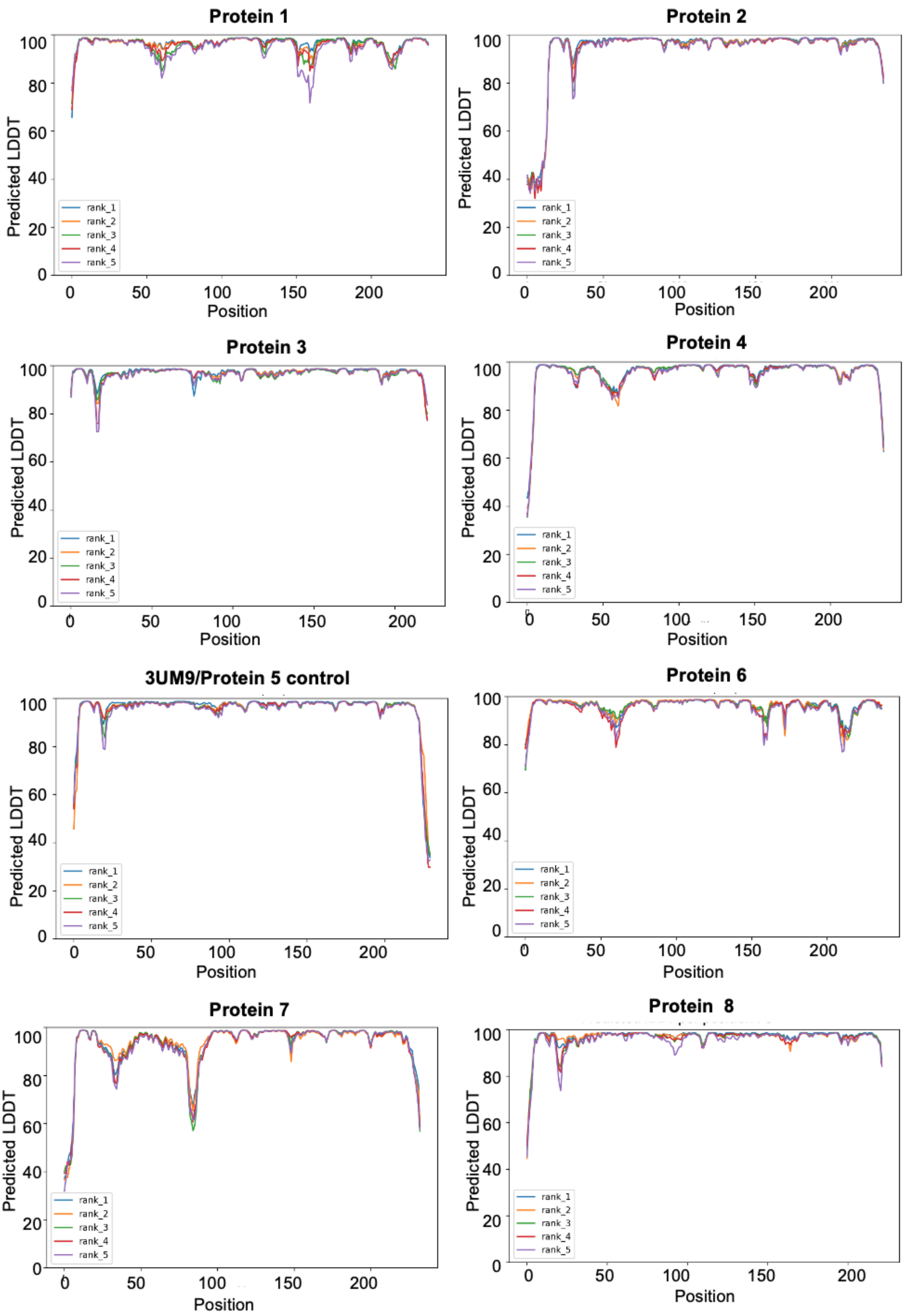

**Figure S15: Predicted Local Difference Distance Test (pLDDT)** of AlphaFold models used for MD simulations. A high pLDDT value corresponds to the confidence of the model at the specific amino acid position. All models are available in the Github repository: https://github.com/MSM-group/gut_microbe_defluorination_paper/

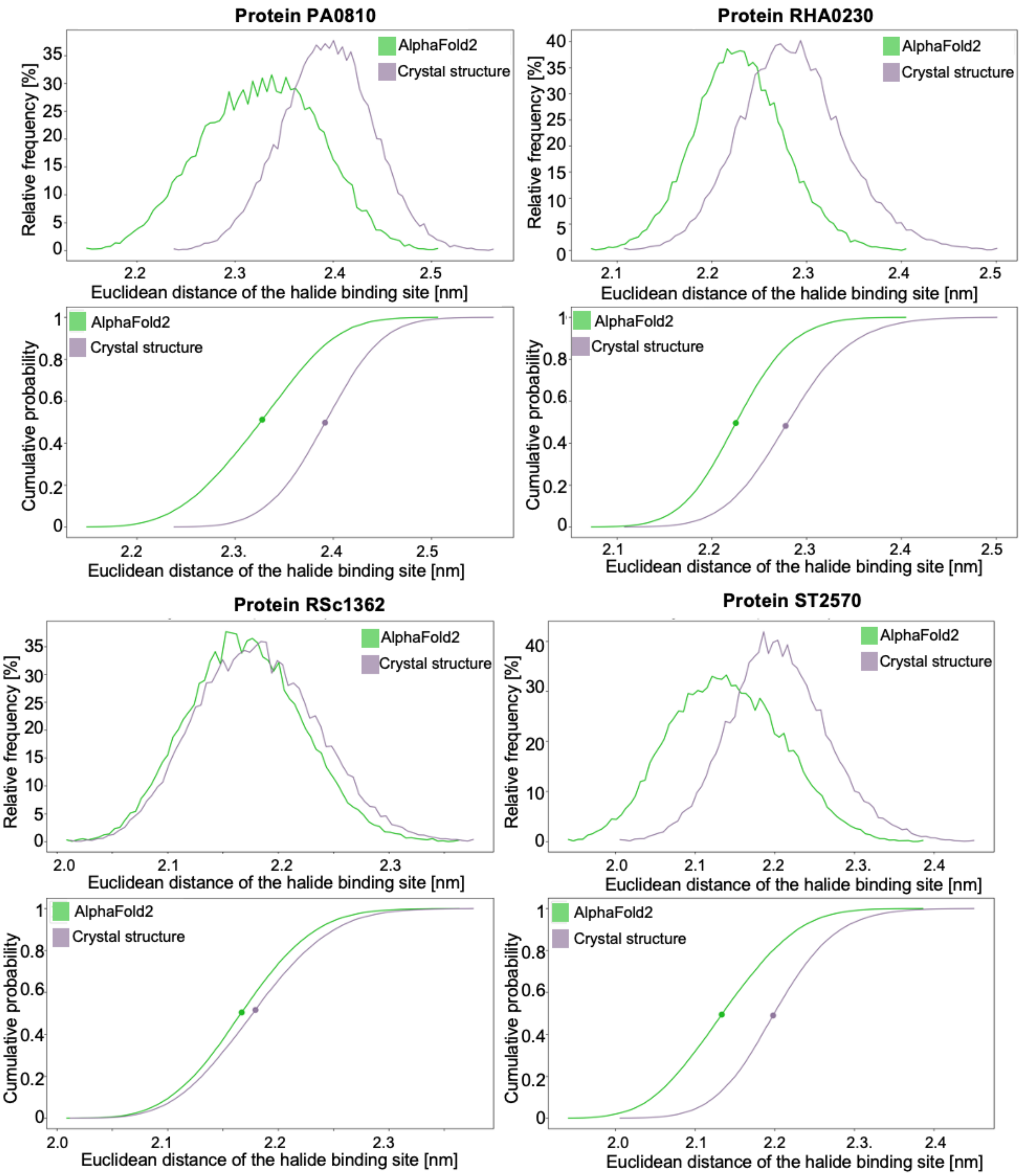

**Figure S16: Molecular dynamics (MD) simulations** of X-ray crystal structure compared to AlphaFold protein structures of identical proteins used in this study. The top graph of each protein represents the relative frequency of Euclidean distances in the halide binding site. The lower graph is representing the cumulative probability with the point showing the middle. Green: AlphaFold2 model; Purple: crystal structure

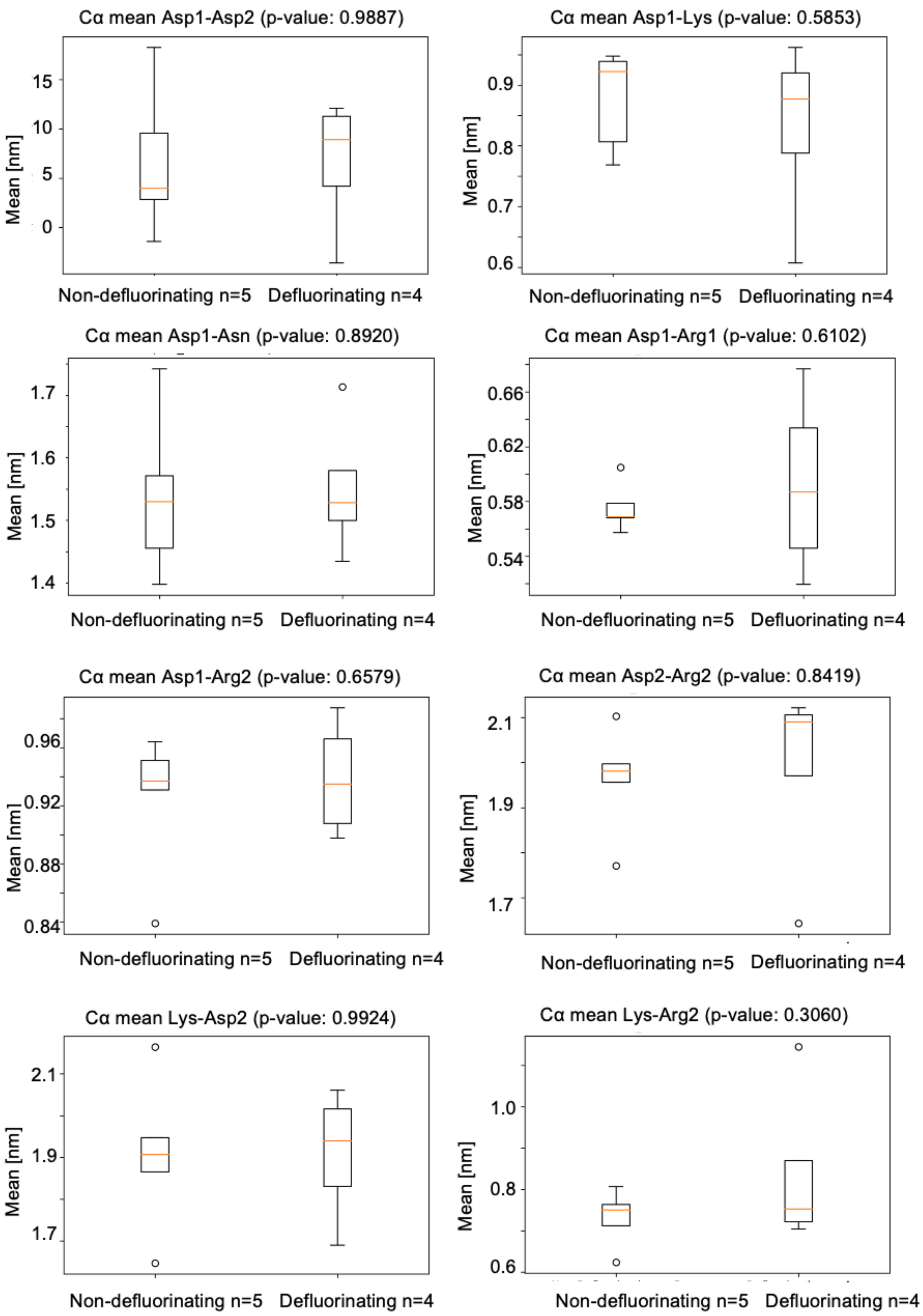
Cα mean Asp1-Asp2 (p-value: 0.9887)
Mean [nm]
15
10
5
0
Non-defluorinating n=5
Defluorinating n=4
Cα mean Asp1-Lys (p-value: 0.5853)
Mean [nm]
0.9
0.8
0.7
0.6
Non-defluorinating n=5
Defluorinating n=4
Cα mean Asp1-Asn (p-value: 0.8920)
Mean [nm]
1.7
1.6
1.5
1.4
Non-defluorinating n=5
Defluorinating n=4
Cα mean Asp1-Arg1 (p-value: 0.6102)
Mean [nm]
0.66
0.62
0.58
0.54
Non-defluorinating n=5
Defluorinating n=4
Cα mean Asp1-Arg2 (p-value: 0.6579)
Mean [nm]
0.96
0.92
0.88
0.84
Non-defluorinating n=5
Defluorinating n=4
Cα mean Asp2-Arg2 (p-value: 0.8419)
Mean [nm]
2.1
1.9
1.7
Non-defluorinating n=5
Defluorinating n=4
Cα mean Lys-Asp2 (p-value: 0.9924)
Mean [nm]
2.1
1.9
1.7
Non-defluorinating n=5
Defluorinating n=4
Cα mean Lys-Arg2 (p-value: 0.3060)
Mean [nm]
1.0
0.8
0.6
Non-defluorinating n=5
Defluorinating n=4

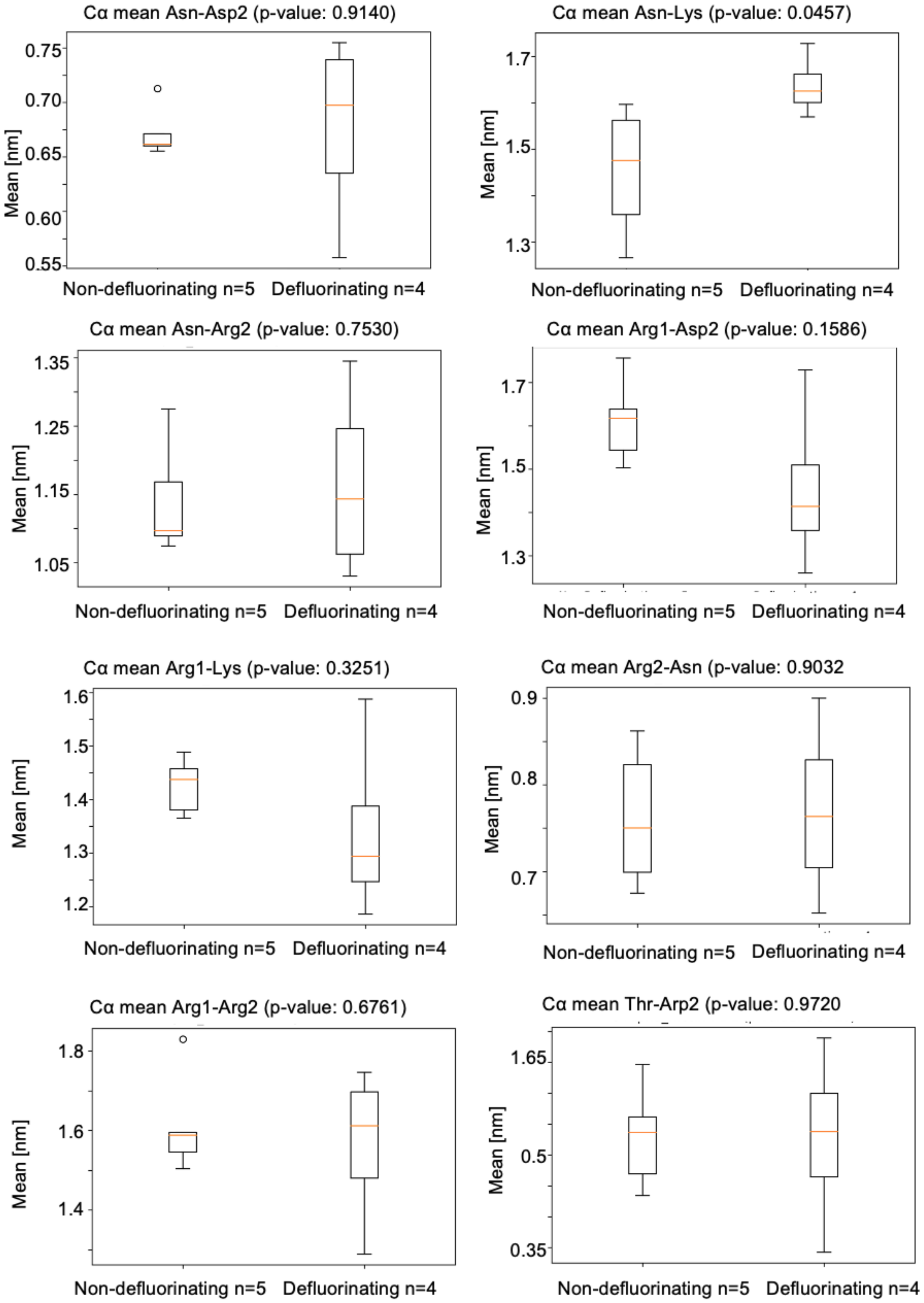
Cα mean Asn-Asp2 (p-value: 0.9140)
Mean [nm]
0.75
0.70
0.65
0.60
0.55
Non-defluorinating n=5
Defluorinating n=4
Cα mean Asn-Lys (p-value: 0.0457)
Mean [nm]
1.7
1.5
1.3
Non-defluorinating n=5
Defluorinating n=4
Cα mean Asn-Arg2 (p-value: 0.7530)
Mean [nm]
1.35
1.25
1.15
1.05
Non-defluorinating n=5
Defluorinating n=4
Cα mean Arg1-Asp2 (p-value: 0.1586)
Mean [nm]
1.7
1.5
1.3
Non-defluorinating n=5
Defluorinating n=4
Cα mean Arg1-Lys (p-value: 0.3251)
Mean [nm]
1.6
1.5
1.4
1.3
1.2
Non-defluorinating n=5
Defluorinating n=4
Cα mean Arg2-Asn (p-value: 0.9032
Mean [nm]
0.9
0.8
0.7
Non-defluorinating n=5
Defluorinating n=4
Cα mean Arg1-Arg2 (p-value: 0.6761)
Mean [nm]
1.8
1.6
1.4
Non-defluorinating n=5
Defluorinating n=4
Cα mean Thr-Arp2 (p-value: 0.9720
Mean [nm]
1.65
0.5
0.35
Non-defluorinating n=5
Defluorinating n=4

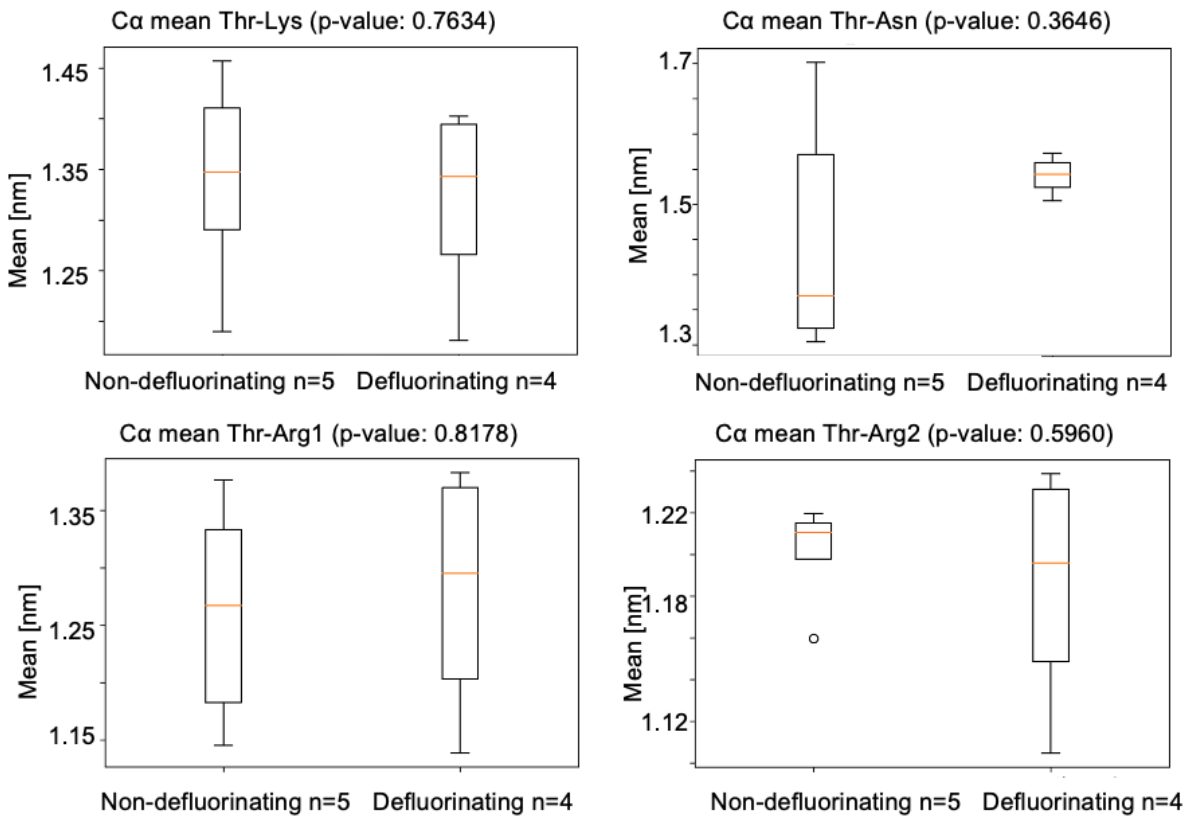

**Figure S17: Boxplots of the mean distances between conserved residues** during a 1 µs MD simulation and p-value of the two-sided pairwise t-tests of these distances between the defluorinating and non-defluorinating haloacid dehalogenases.

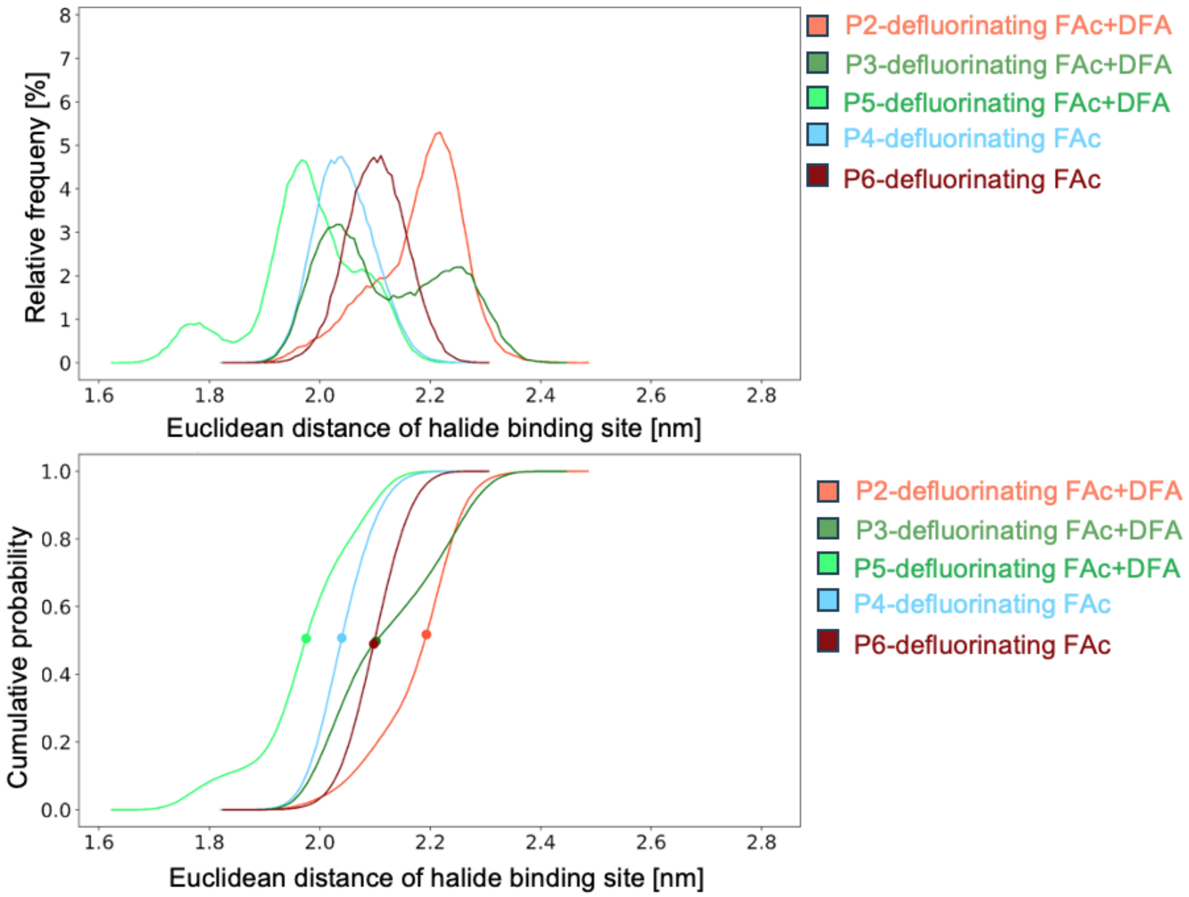

**Figure S18**: **Halide binding site (HBS) change during MD simulations** comparing HADs able to defluorinate fluoroacetate (FAc) and difluoroacetate (DFA) with HADs only defluorinating FAc. The top graph represents the relative frequency of Euclidean distances in the halide binding site. The lower graph is representing the cumulative probability with the point showing the midpoint.

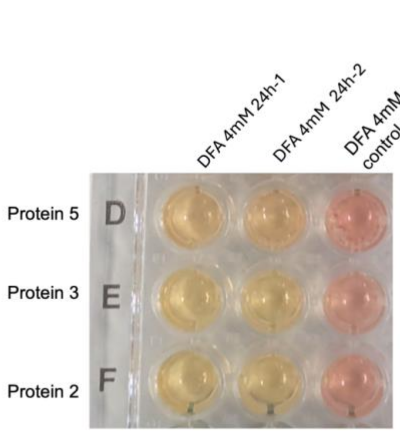

**Conversion of DFA (4 mM) colorimetric assay**

| | 24h-1 | | 24h-2 | | 24h-boiled protein control | |
|---|---|---|---|---|---|---|
| | $F^-$ conc. | conv. | $F^-$ conc. | conv. | $F^-$ conc. | conv. |
| Protein 5 | 602 μM | 7.7 % | 546 μM | 7 % | 98 μM | 1.19 % |
| Protein 3 | 861 μM | 10.5 % | 735 μM | 9.31 % | 147 μM | 1.82 % |
| Protein 2 | 777 μM | 9.45 % | 672 μM | 8.4 % | 126 μM | 1.61 % |

**Figure S19**: **Fluoride concentration and difluoroacetate conversion** for proteins 2,3 and 5 tested with the colorimetric assay. The difluoroacetate standard provided by the manufacturer has a purity of >98%.

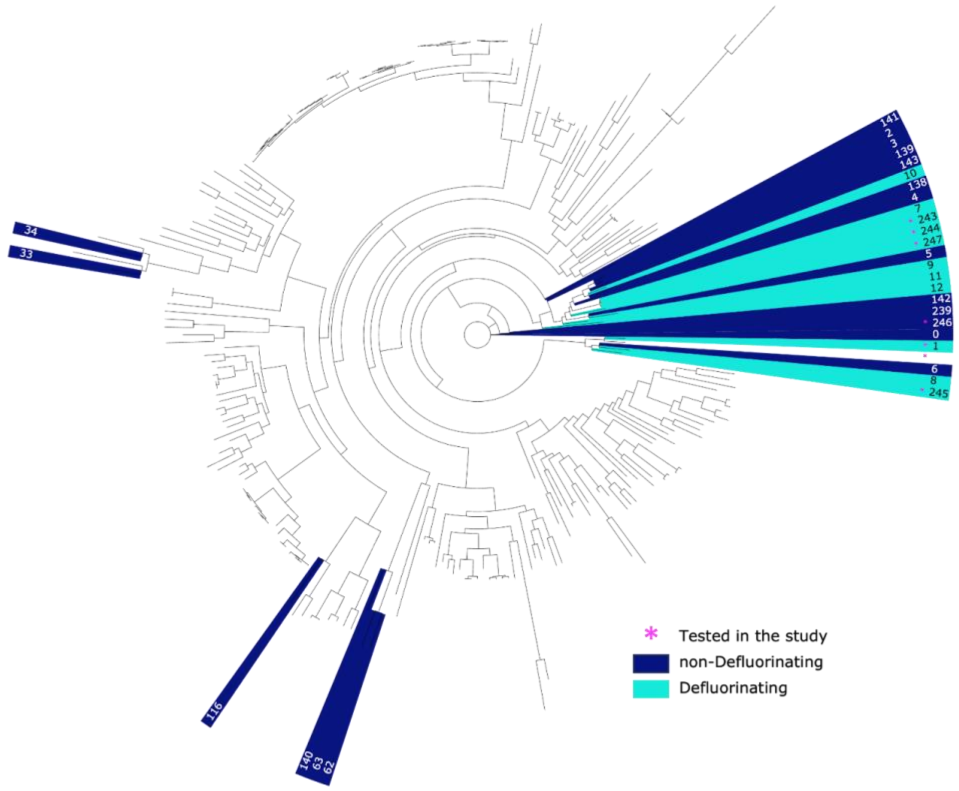

**Figure S20: Maximum-likelihood phylogenetic analysis of HAD-like superfamily enzymes**. Colored leaves are the subset of the experimentally confirmed dehalogenating enzymes relative to HAD-like superfamily enzymes performing other functions (e.g., phosphoryl transfer). The different colors correspond to the defluorination properties. Our data set contained 248 homologous HAD-like enzymes, 29 of which were experimentally confirmed dehalogenases. Among the 29 dehalogenases, ten were confirmed to defluorinate (four newly identified in this study, six from literature). The indices in the tree correspond to the indices in supplementary table S9 with the full metadata of the HAD-like dataset. Enzymes that have been tested in this study are marked with an asterisk.

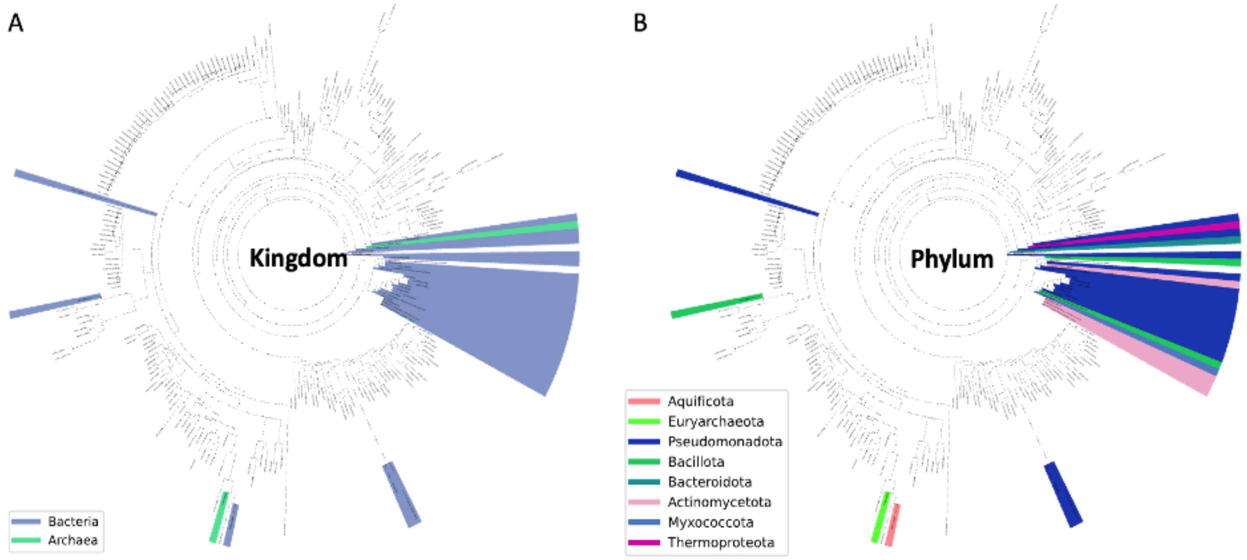

**Figure S21: Dehalogenase subset location in the HAD-like family phylogenetic tree and their taxonomy.** A) Colored leaves correspond to the dehalogenating enzymes subset and show that they are sourced solely from Bacteria and Archaea. B) The corresponding phyla of the dehalogenase subset is shown in A).

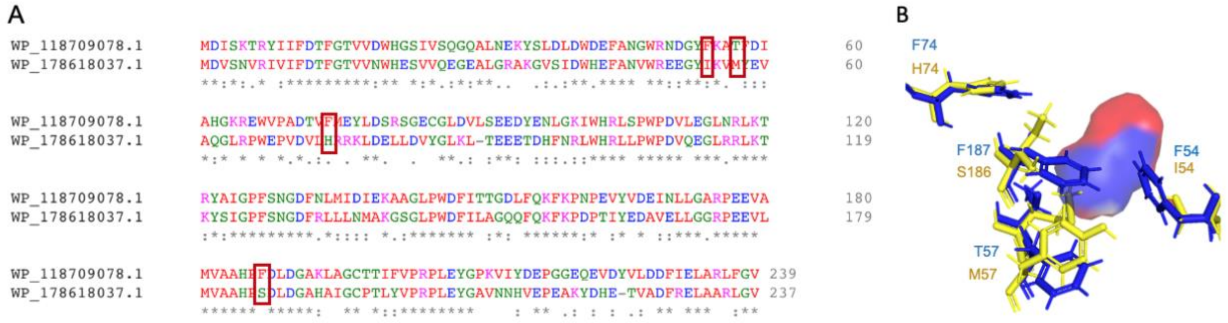

**Figure S21: A) Sequence alignment** of the defluorinating P6 and the non defluorinating protein P1. Red boxes: the only amino acids different and located in the active center. B) location of the marked amino acids with the active pocket. Blue (WP_118709078.1) = P1, Yellow (WP_178618037.1) = P6.

Substrate: chloroacetate fluoroacetate

**P1 - F54I**

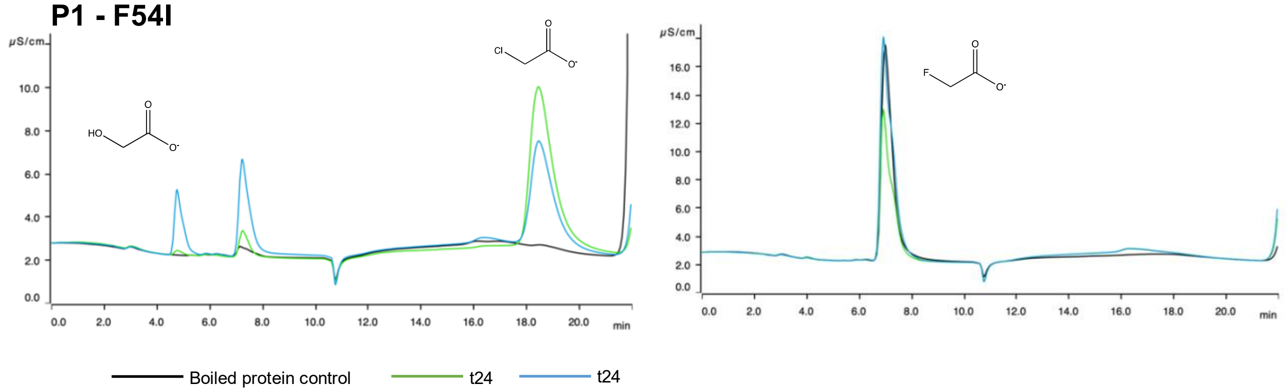

| Protein variant | substrate | % conversion | concentration glycolate [mM] |
|---|---|---|---|
| P1-F54I | chloroacetate 4mM | 25% | 1.01 |
| P1-F54I | fluoroacetate 4mM | 0% | <LOD |

**P1 - T57M**

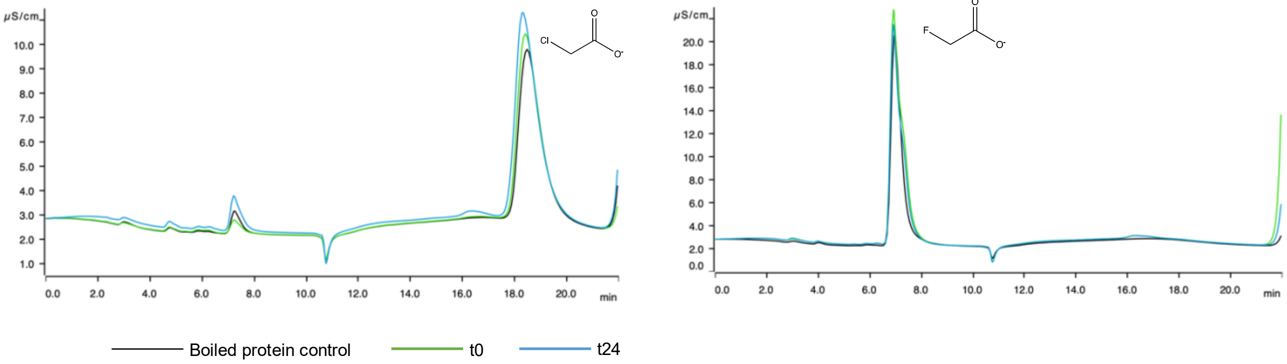

| Protein variant | substrate | % conversion | concentration glycolate [mM] |
|---|---|---|---|
| P1-T57M | chloroacetate 4mM | 0% | <LOD |
| P1-T57M | fluoroacetate 4mM | 0% | <LOD |

**P1 - H74F**

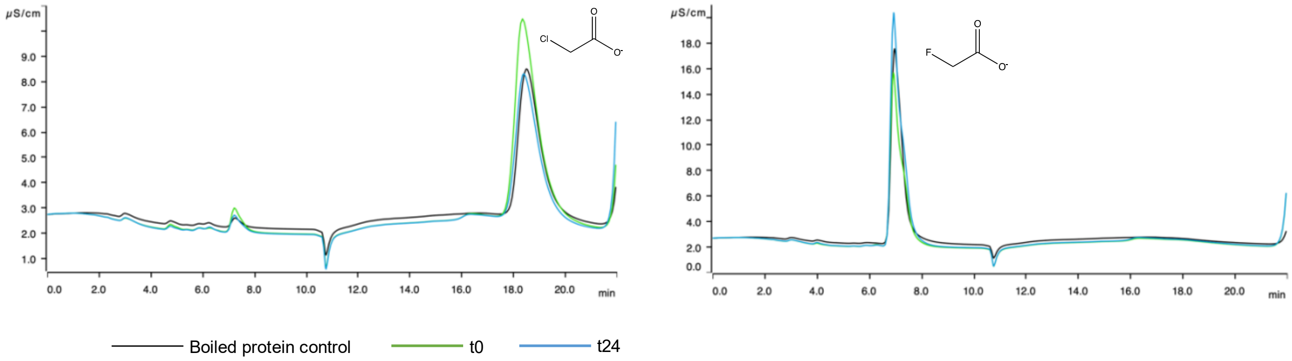

| Protein variant | substrate | % conversion | concentration glycolate [mM] |
|---|---|---|---|
| P1-H74F | chloroacetate 4mM | 0% | <LOD |
| P1-H74F | fluoroacetate 4mM | 0% | <LOD |

**Figure S22: Ion chromatography** traces of the single amino acid variants for protein P1 (*Enterocloster aldenensis*) with two different substrates. Left column: chloroacetate; Right column: fluoroacetate at different timepoints. Black: boiled protein control; Green: t0h; Blue: t24h.

The tables show the conversion and measured glycolate concentration. Glycolate is used for this calculation due to possible interfering chloride contaminations at min 7.0.

Substrate: chloroacetate fluoroacetate

**P1 - F54I + T57M**

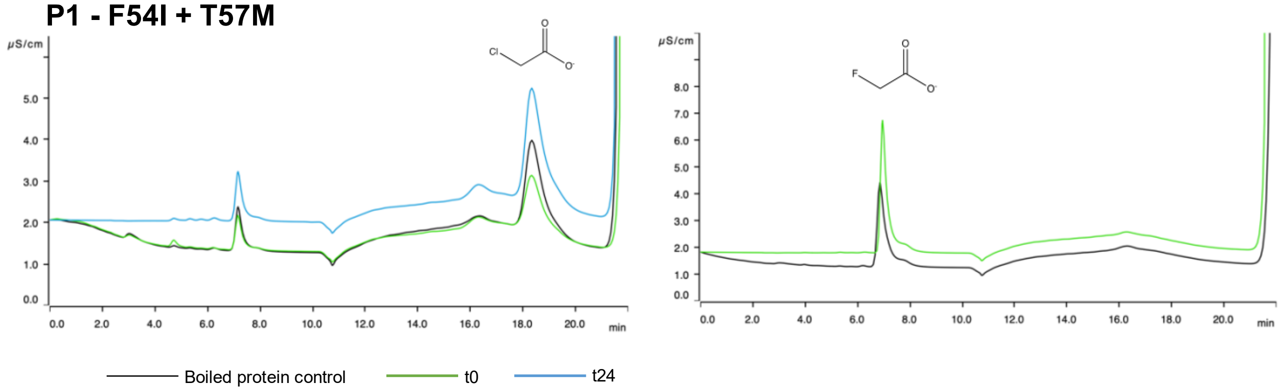

| Protein variant | substrate | % conversion | concentration glycolate [mM] |
|---|---|---|---|
| P1-F54I+T57M | chloroacetate 1 mM | 0% | <LOD |
| P1-F54I+T57M | fluoroacetate 1 mM | 0% | <LOD |

**P1 - F54I+ H74F**

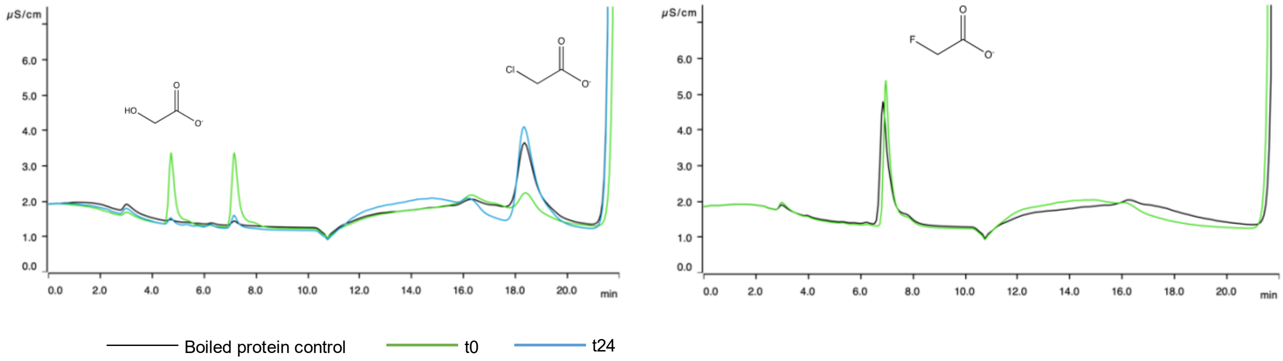

| Protein variant | substrate | % conversion | concentration glycolate [mM] |
|---|---|---|---|
| P1-F54I+T57M | chloroacetate 1 mM | 23% | 0.326 |
| P1-F54I+T57M | fluoroacetate 1 mM | 0% | <LOD |

**P1 - F54I + F187S**

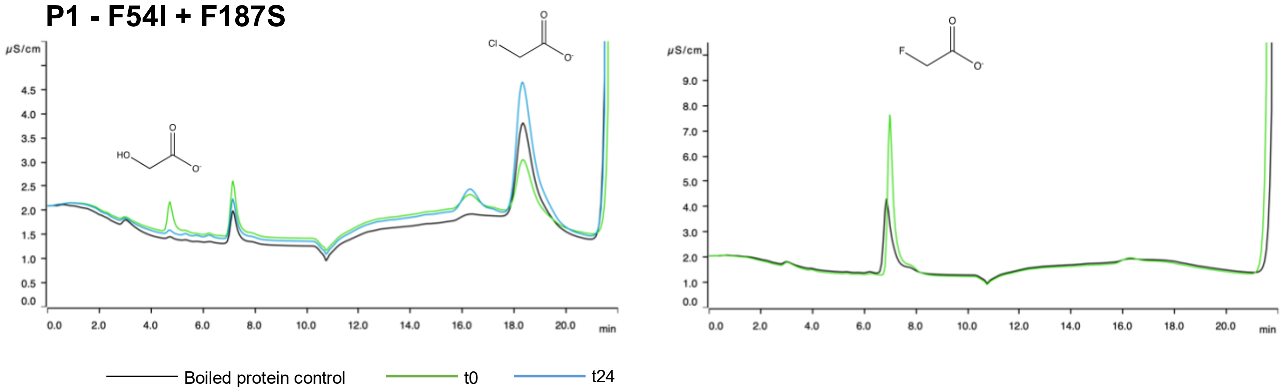

| Protein variant | substrate | % conversion | concentration glycolate [mM] |
|---|---|---|---|
| P1-F54I+T57M | chloroacetate 1 mM | 1.6% | 0.07 |
| P1-F54I+T57M | fluoroacetate 1 mM | 0% | <LOD |

**Figure S23: Ion chromatography traces** of the double mutants for protein P1 (*Enterocloster aldenensis*) with two different substrates. Left column: chloroacetate; Black: boiled protein control; Blue: t0h; Green: t24h. The tables show the conversion and measured glycolate concentration. Glycolate is used for this calculation due to possible interfering chloride contaminations at min 7.0.

Substrate: chloroacetate fluoroacetate

**P1 - F54I + T57M + H74F**

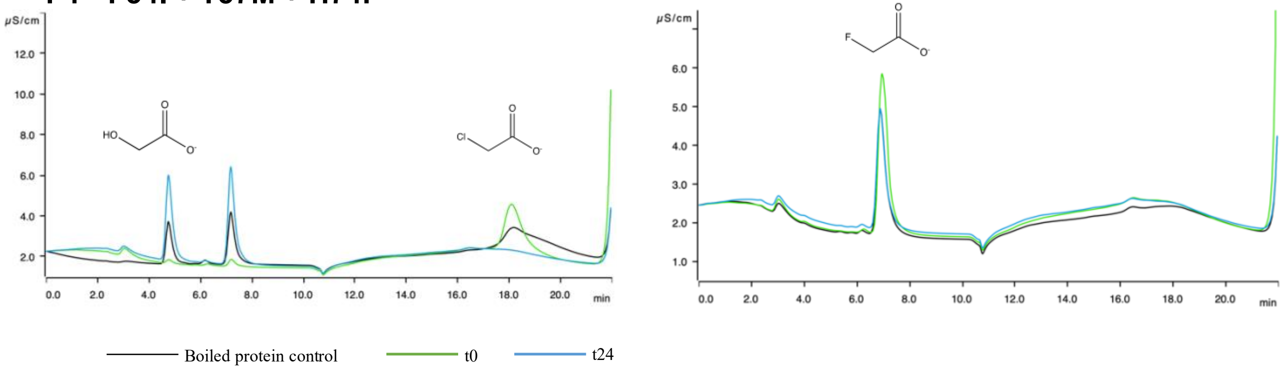

| Protein variant | substrate | % conversion | concentration glycolate [mM] |
|---|---|---|---|
| P1-F54I+T57M+H74F | chloroacetate 1 mM | 118% | 1.184 |
| P1-F54I+T57M+H74F | fluoroacetate 1 mM | 0% | <LOD |

**P1 - F54I + H74F + F187S**

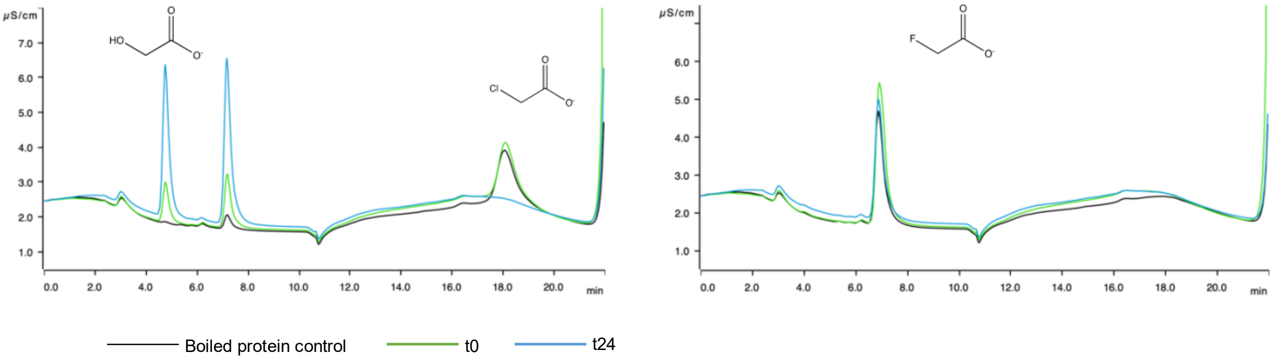

| Protein variant | substrate | % conversion | concentration glycolate [mM] |
|---|---|---|---|
| P1-F54I+H74F+F187S | chloroacetate 1 mM | 112% | 1.128 |
| P1-F54I+H74F+F187S | fluoroacetate 1 mM | 0% | <LOD |

**P1 - F54I + F187S + T57M**

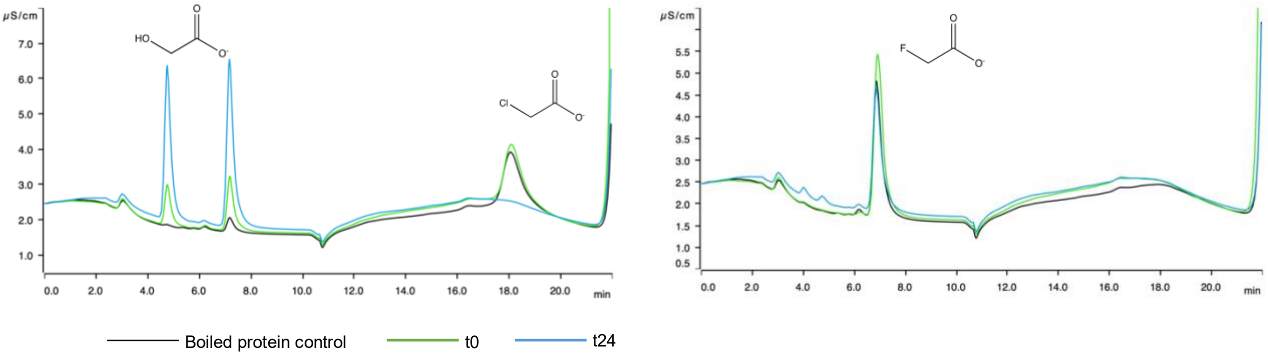

| Protein variant | substrate | % conversion | concentration glycolate [mM] |
|---|---|---|---|
| P1-F54I+T57M+F187S | chloroacetate 1 mM | 117% | 1.172 |
| P1-F54I+T57M+F187S | fluoroacetate 1 mM | 0% | <LOD |

**Figure S24: Ion chromatography** traces of the triple mutants for protein WP_118709078.1 (*Enterocloster aldenensis*) with two different substrates. Left column: chloroacetate Black: boiled protein control; Green: t0h; Blue: t24h. The tables show the conversion and measured glycolate concentration. Glycolate is used for this calculation due to possible interfering chloride contaminations at min 7.0. Conversion values >100% are due to instrument variability and slight evaporation which could not be fully avoided.

Substrate: chloroacetate fluoroacetate

**P1 - F54I + F187S + T57M + H74F**

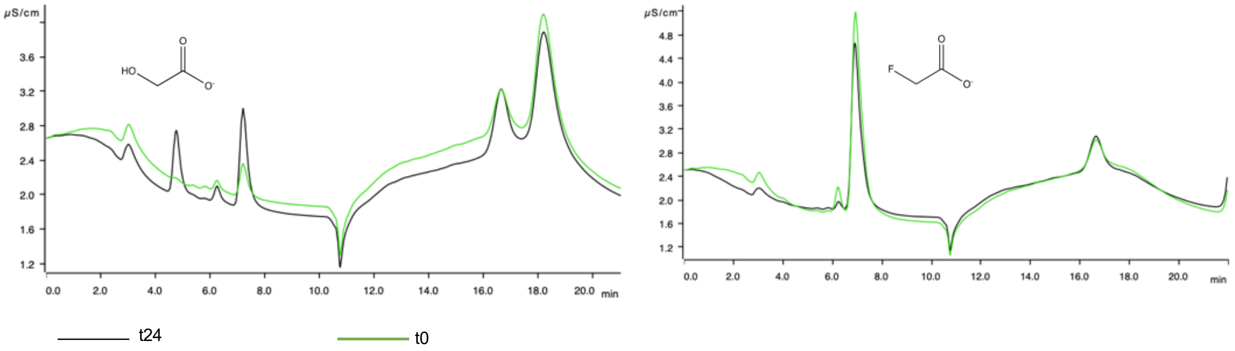

| Protein variant | substrate | % conversion | concentration glycolate [mM] |
|---|---|---|---|
| P1-F54I+T57M+F187S | chloroacetate 1 mM | 8.6% | 0.086 |
| P1-F54I+T57M+F187S | fluoroacetate 1 mM | 0% | <LOD |

**Figure S25: Ion chromatography** traces of the quadruple mutants for protein P1 (*Enterocloster aldenensis*) with two different substrates. Left column: chloroacetate Black: t24; Green: t0h. The table shows the conversion and measured glycolate concentration. Glycolate is used for this calculation due to possible interfering chloride contaminations at min 7.0.

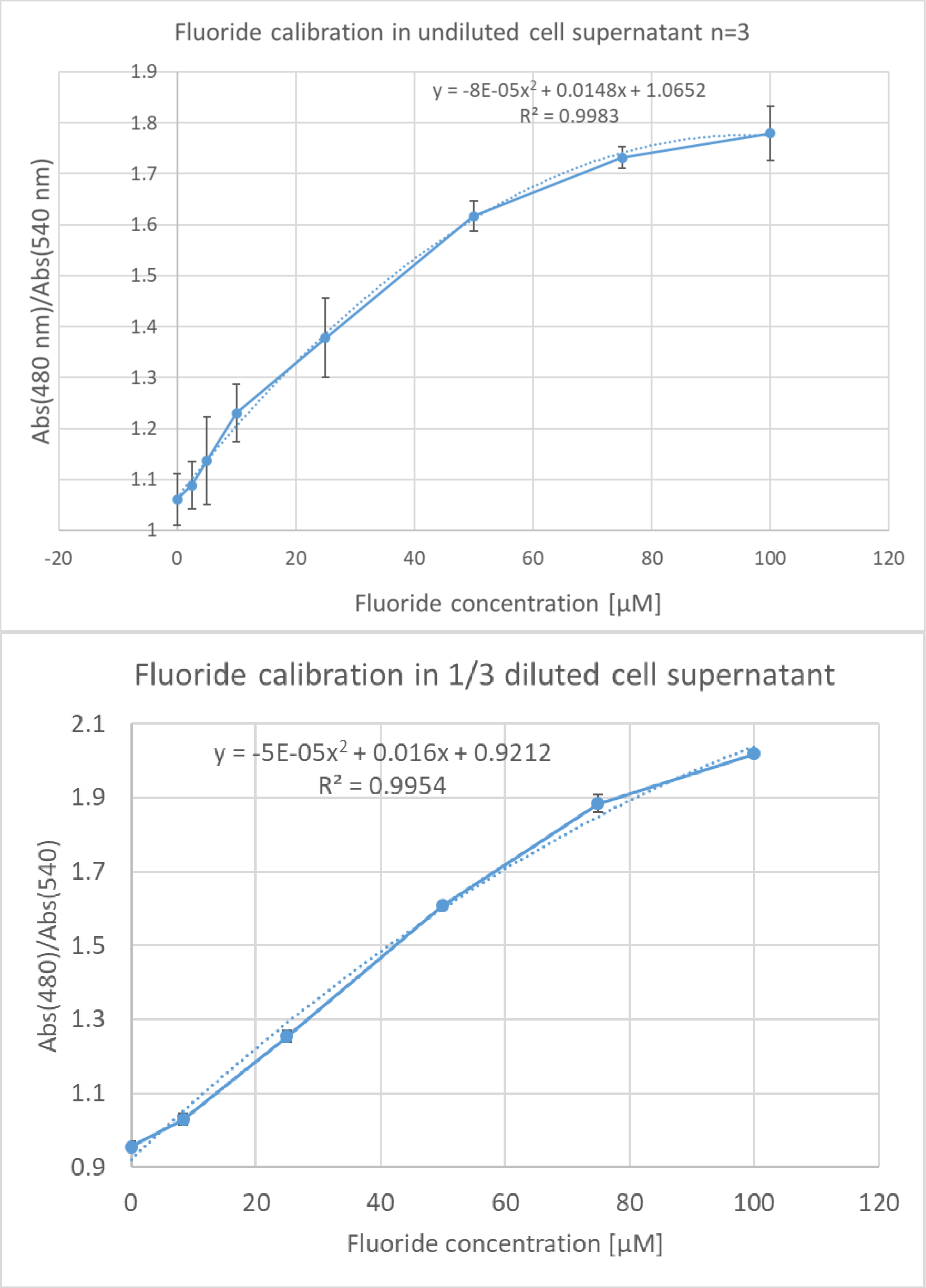

**Figure S26: Miniaturized new xylenol orange colorimetric assay:** Standards in undiluted and ⅓ diluted cell supernatant of *E. coli* producing inactive enzyme, which were used to calibrate the measurements of the alanine scanning library in the respective dilution.

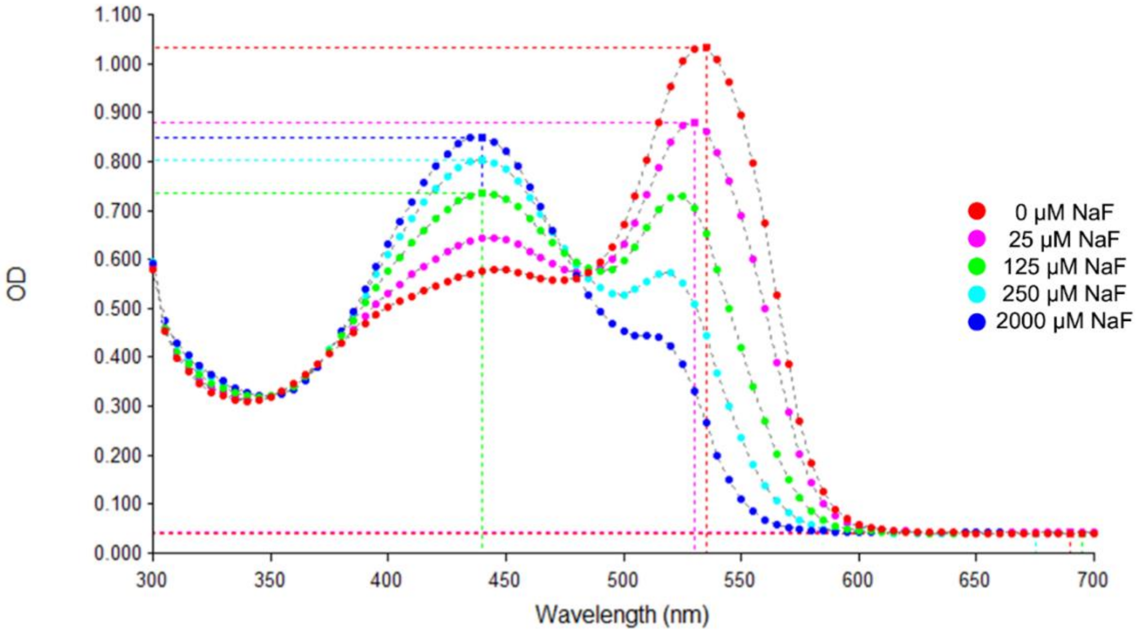

**Figure S27: Isosbestic points measured using the new miniaturized xylenol orange assay.** Absorbance spectra are shown for a standard curve of fluoride ion concentrations.

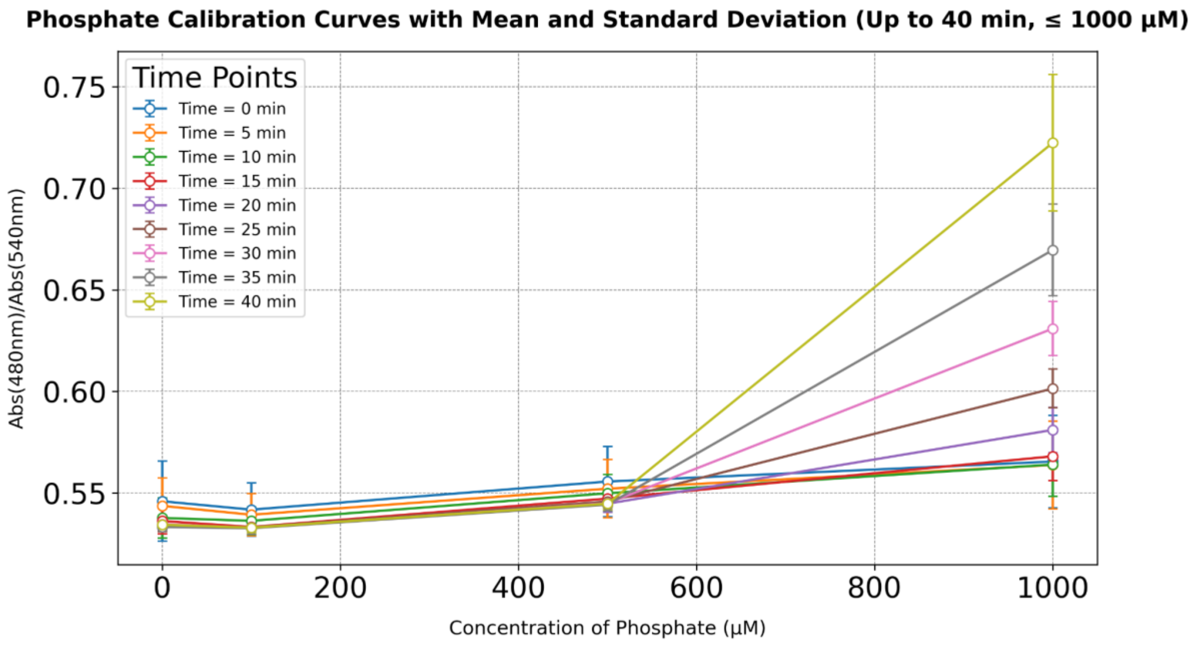

**Figure S28: Interference of phosphate on the fluoride assay relative to concentration and time.** 4 replicates of each phosphate concentration were measured during 40 min. No change in absorption was measured up to a concentration of phosphate of 0.5 mM. Phosphate was shown to disrupt the red complex similarly to fluoride, but at higher concentrations and more slowly.

P9 *Priestia megaterium*

Substrate: chloroacetate fluoroacetate

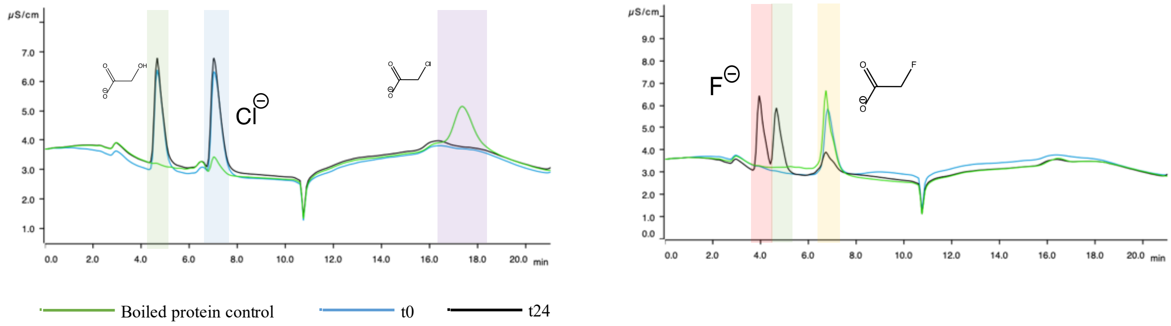

| Protein | substrate | % conversion | concentration glycolate [mM] |
|---|---|---|---|
| P9 | chloroacetate 1 mM | 62% | 0.62 |
| P9 | fluoroacetate 1 mM | 50% | 0.5 |

**Figure S29: Ion chromatography** traces of protein P9 (*Priestia megaterium)* with two different substrates. Green: control; Blue: t0h; Black: t24h. Chloroacetate shaded in purple, fluoroacetate shaded in yellow, glycolate shaded in green, chloride shaded in blue, and fluoride shaded in red. The table shows the conversion and measured glycolate concentration. Glycolate is used for this calculation due to possible interfering chloride contaminations.

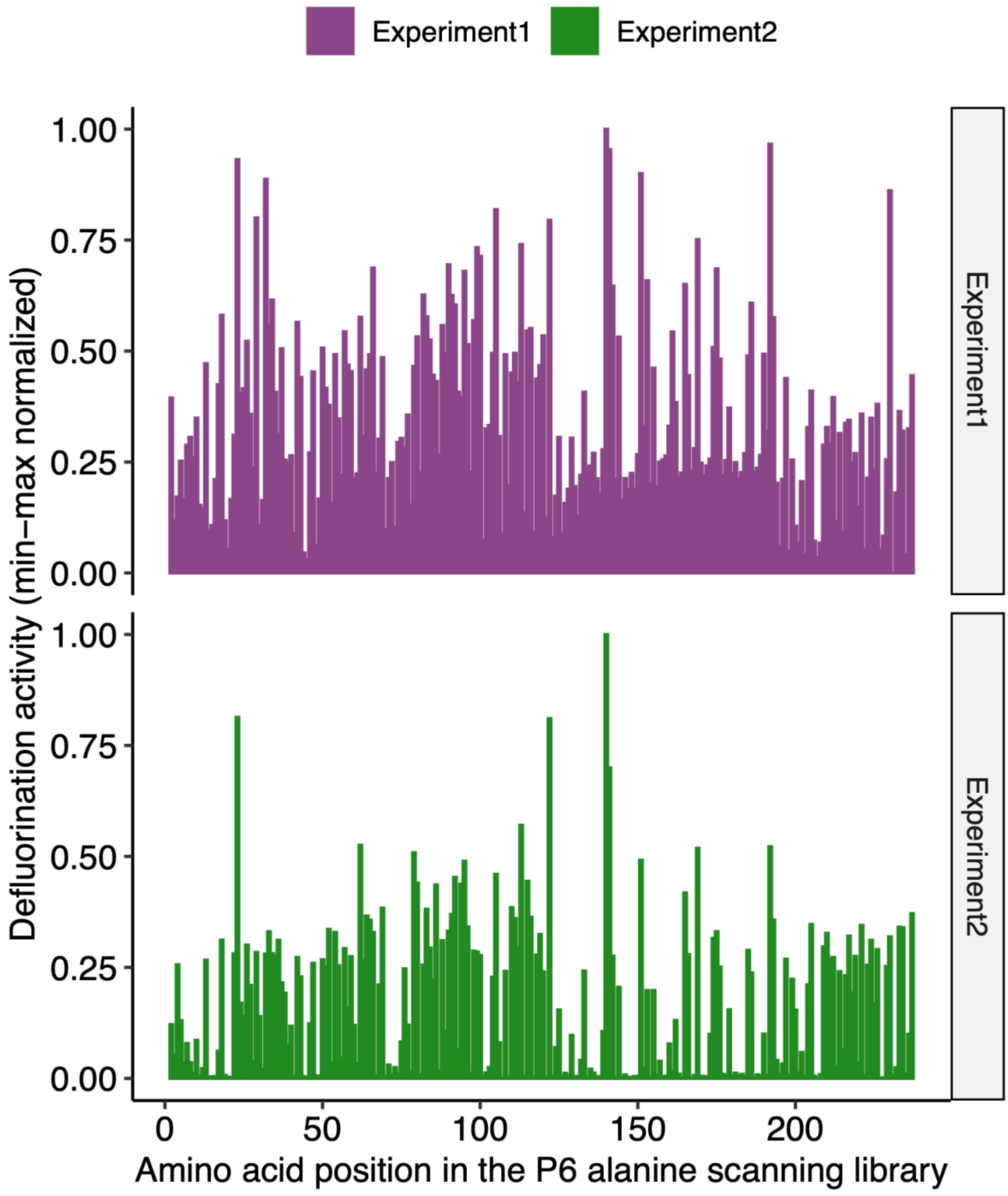

**Figure S30: Comparison of Experiment 1 and Experiment 2** results from defluorination activity screening of the alanine scanning library. Overall background levels are higher in Experiment 1 due to the introduction of additional washing and dilution steps in Experiment 2 which increased the signal-to-noise ratio.

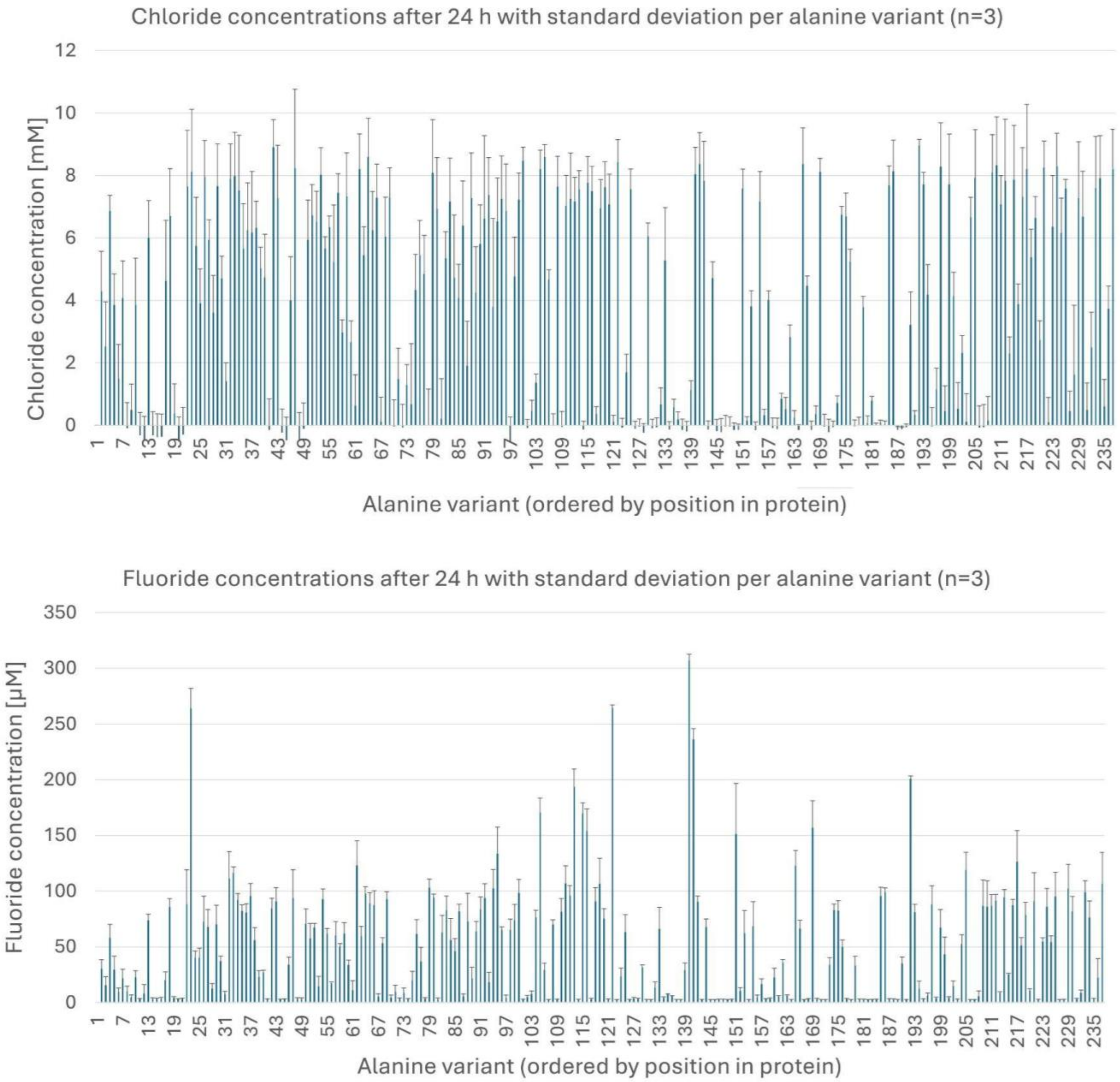

**Figure S31: Mean fluoride and chloride concentrations and standard deviations of 3 independent replicates for each mutant.** Fluoride and chloride concentrations were measured with colorimetric assays. The first measurement corresponds to the second amino acid mutated to alanine.

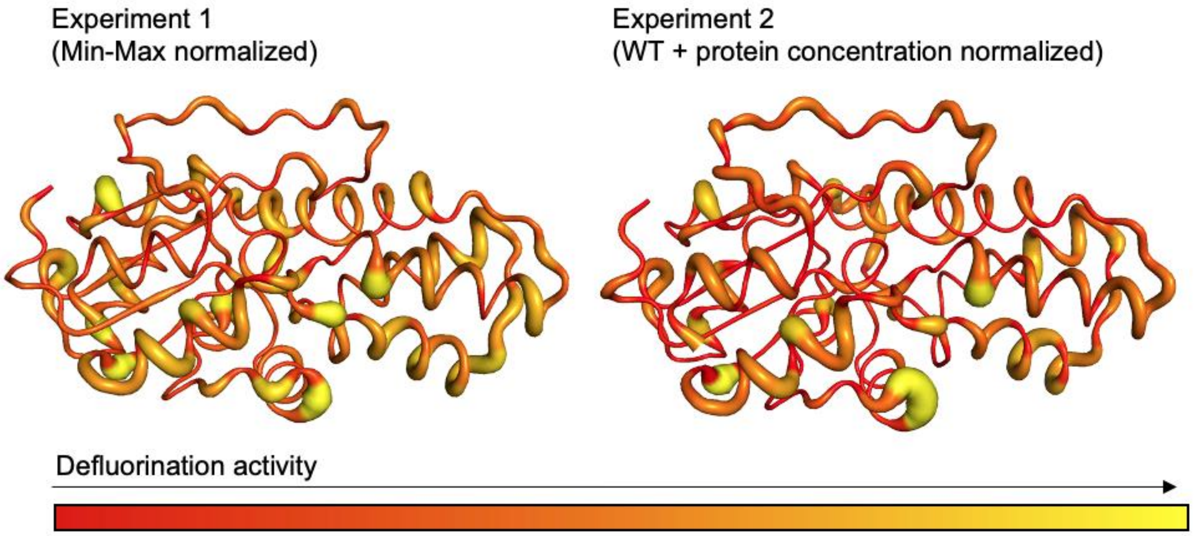

**Figure S32: Defluorination activity** of the alanine scanning for protein P6 for experiment 1 and 2. The defluorination activity is based on the measured fluoride concentration of our alanine scanning experiment. The representation shows the merged protein variants. The higher the measured fluoride concentration for the individual variant the thicker and brighter is the line in the representation at the location of the mutated amino acid.

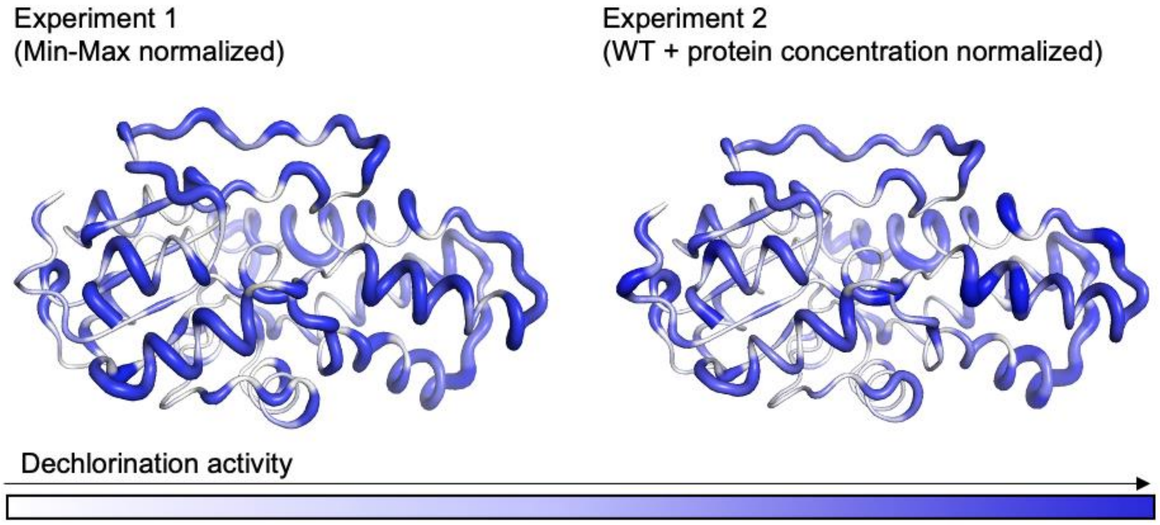

**Figure S33: Dechlorination activity** of the alanine scanning for protein P6 for experiment 1 and 2. The dechlorination activity is based on the measured chloride concentration of our alanine scanning experiment. The representation shows the merged protein variants. The higher the measured chloride concentration for the individual variant the thicker and bluer is the line in the representation at the location of the mutated amino acid.

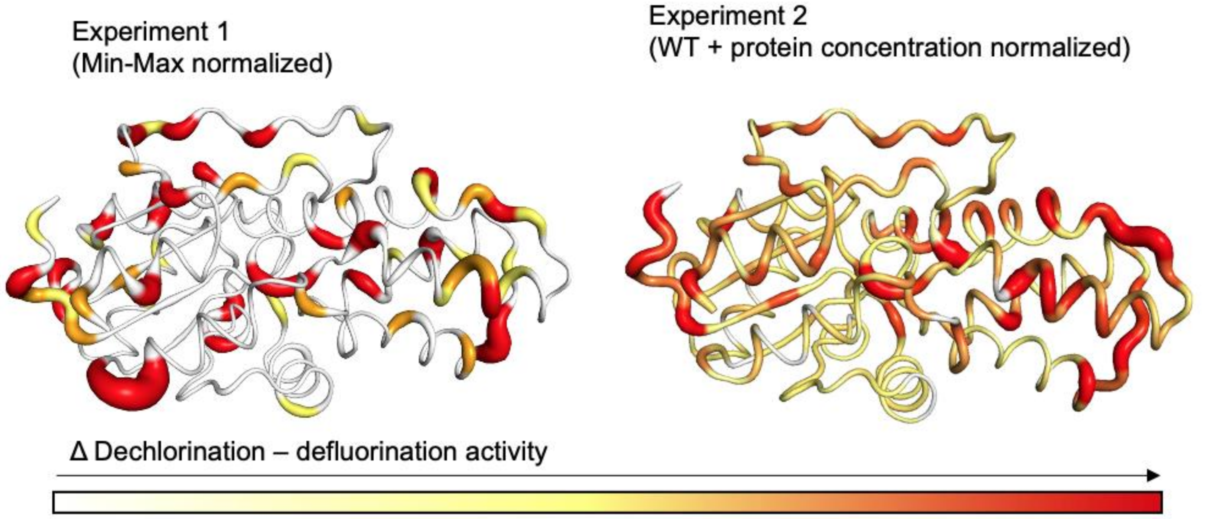

**Figure S34: Defluorination influence. Defluorination activity subtracted from the dechlorination activity** results of the alanine scanning for protein P6 from experiment 1 and 2. The defluorination activity and the dechlorination activity is based on the measured fluoride and chloride concentration of our alanine scanning experiments. The representation shows the merged protein variants. P6 alanine variants which lost their defluorination activity but kept their dechlorination activity influencing the defluorination. The thicker and redder the line in the representation, the greater the difference between measured chloride and fluoride concentrations for the P6 variant with the alanine mutation at that location.

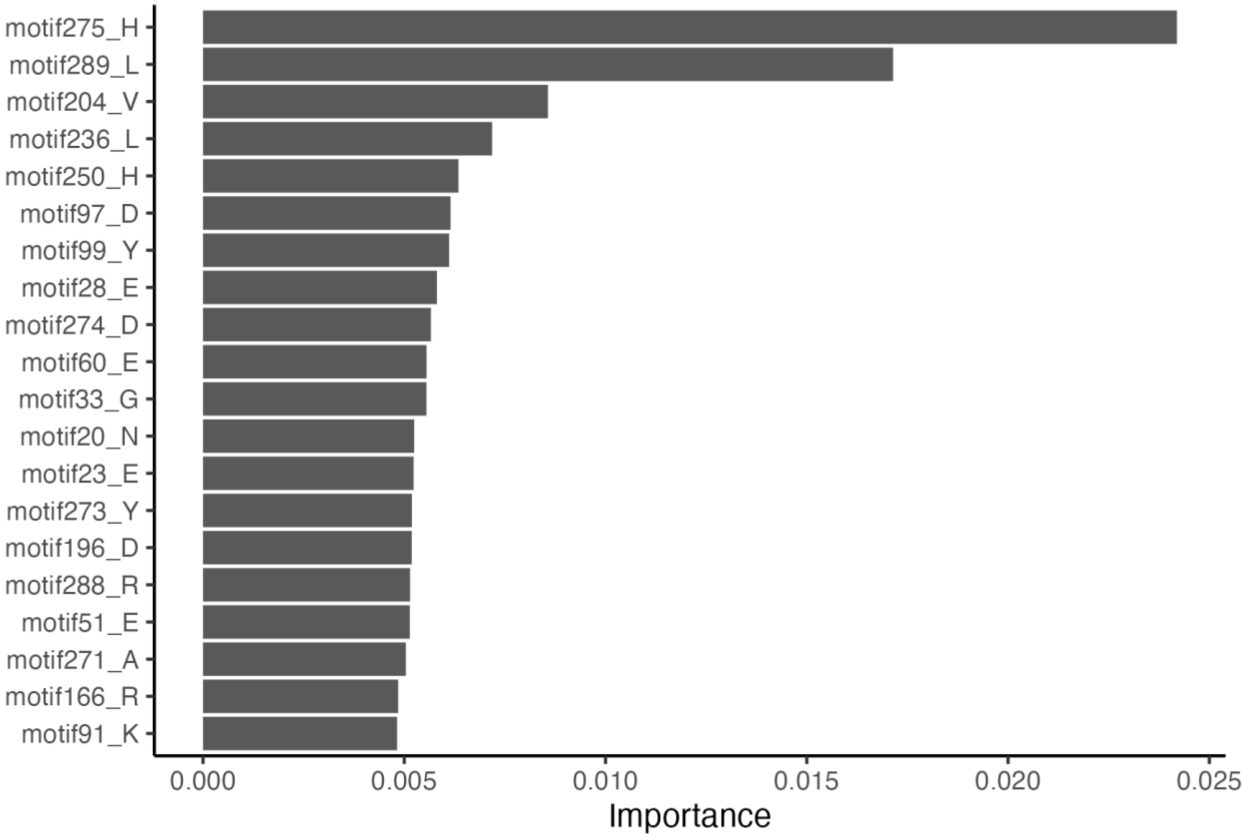

**Figure S35: Experiment 1 trained random forest classification model** identified the top twenty most important features to separate defluorinating from non-defluorinating proteins. Feature importance was determined by permutation.

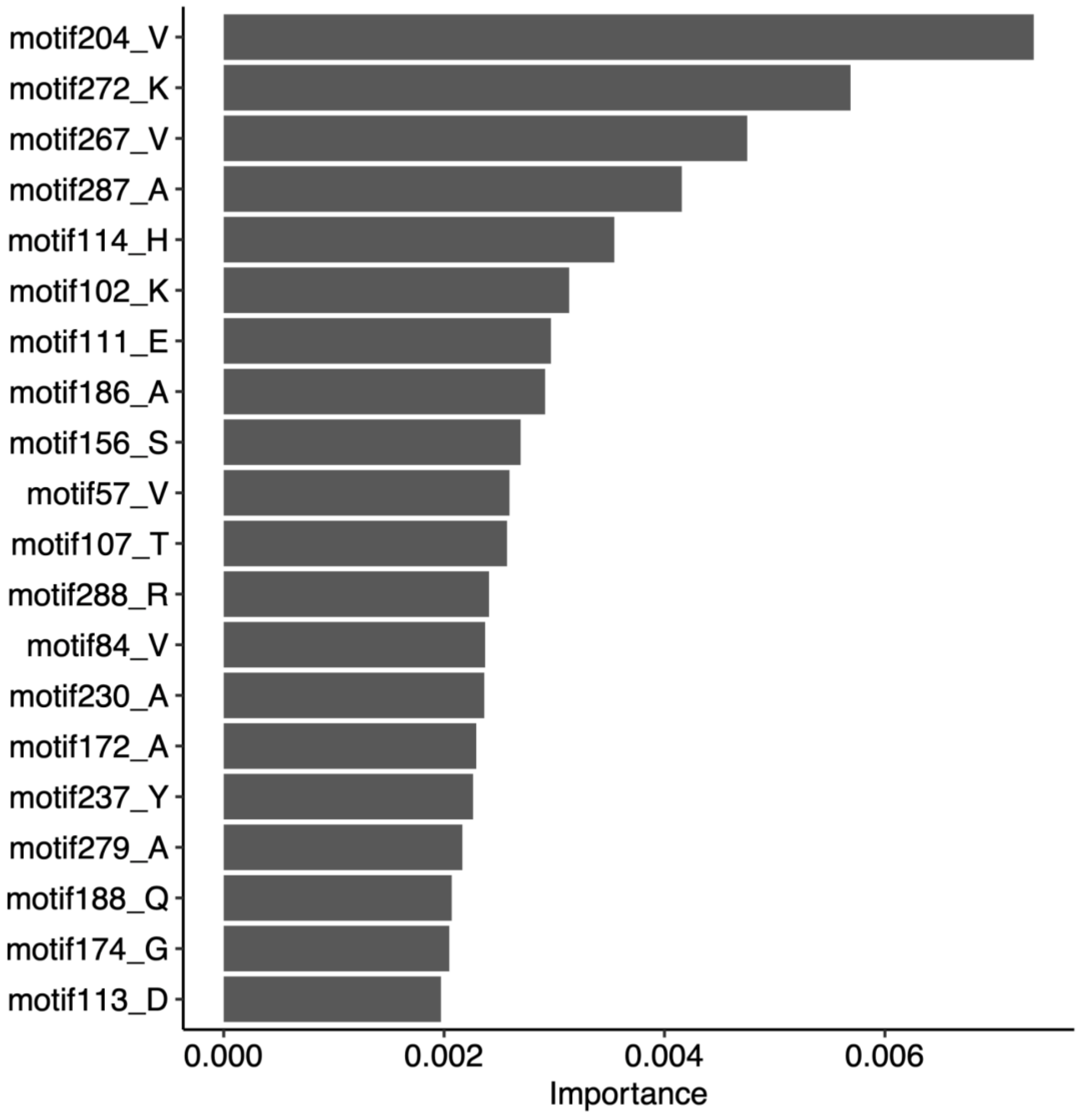

**Figure S36**: **Experiment 2 trained random forest classification model** identified the top twenty most important features to separate defluorinating from non-defluorinating proteins. Feature importance was determined by permutation.

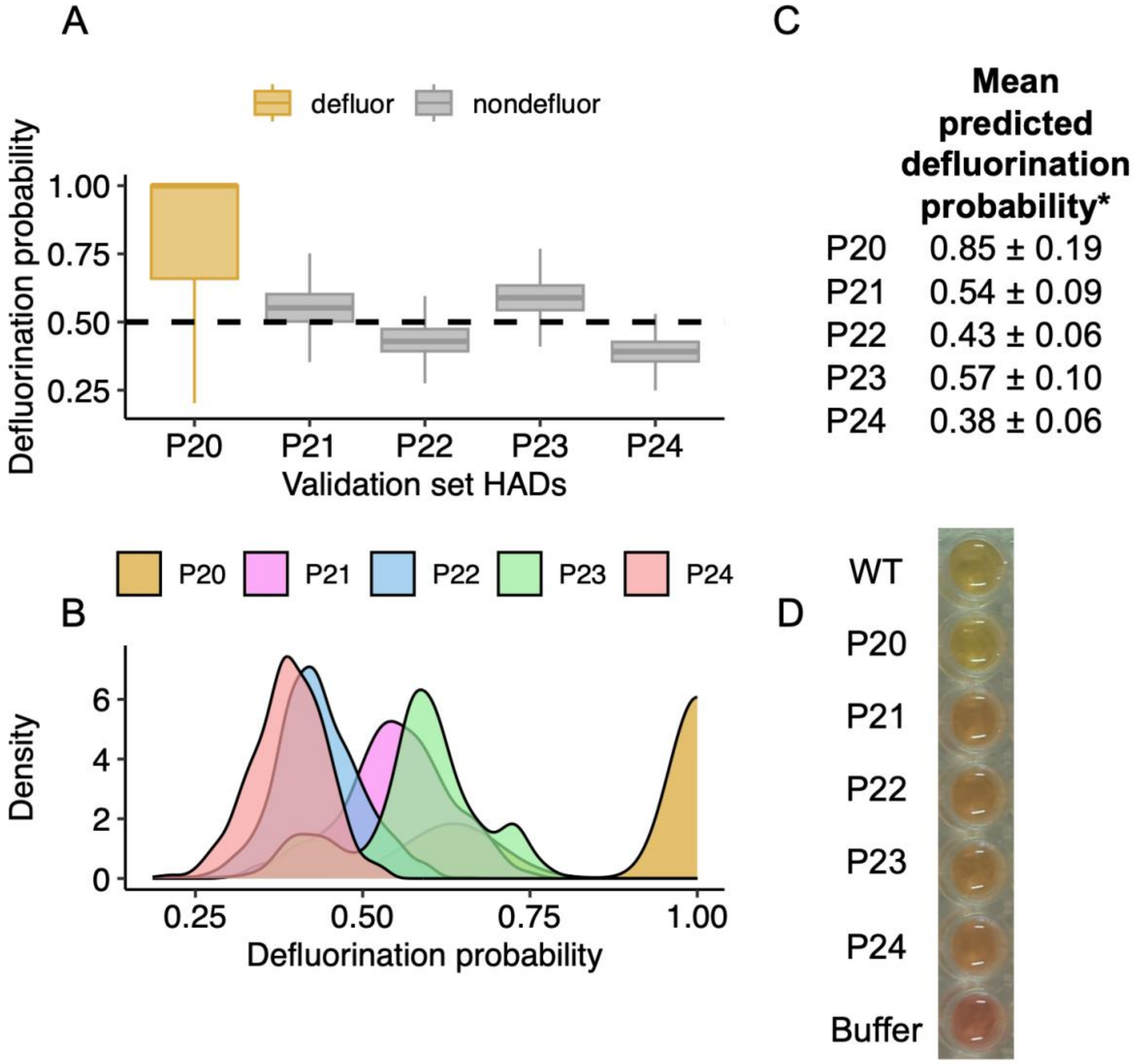

**Figure S37.** A) Comparison of defluorination probabilties predicted for five new validation set HADs. P20 = *Raoultibacter phocaeensis (*WP_139652913.1); P21 = *Stellaceae bacterium* (HTS94301.1); P22 = *Ca. Rokuibacteriota (*PYN88313.1)*;* P23 = *Gemmatimonadota bacterium (*MDH3205452.1); P24 = *Eubacterium sp. (*MBS6900484.1). B) Density plots of the probabilities demonstrate the higher probability distribution of P20 relative to other validation set HADs. C) Mean predicted defluorination probability scores for the validation set sequences predicted by random forest classification. D) Results from the colorimetric fluoride ion detection assay demonstrating defluorination activity of P20 comparable to wild-type (WT) while P21-P24 have no activity (slight assay discoloration is due to protein, comparable to a buffer control).

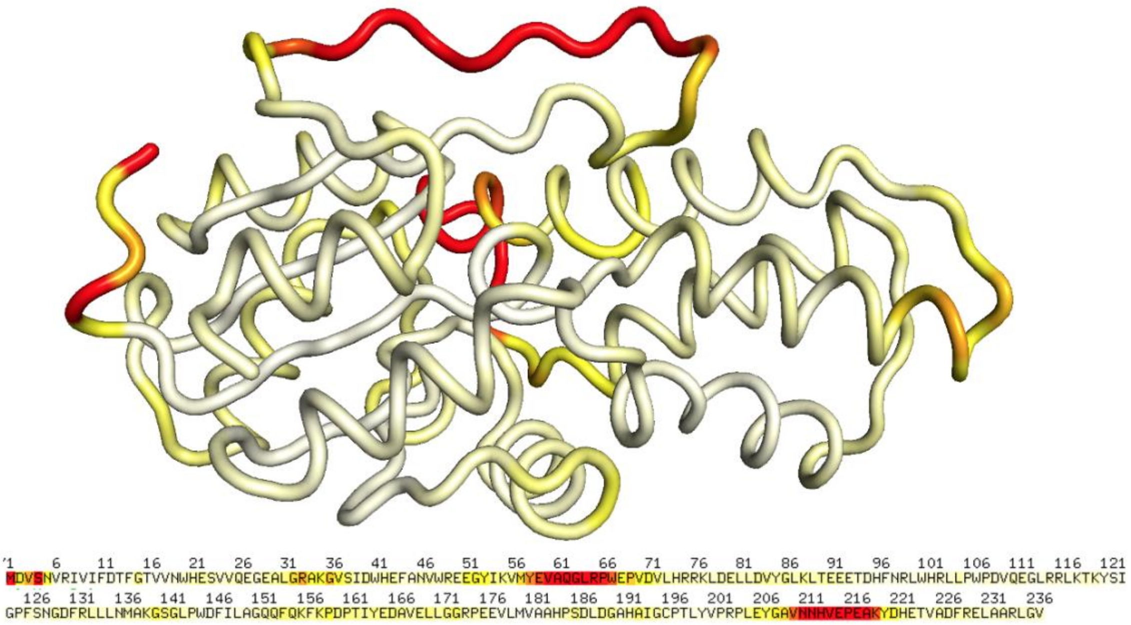

**Figure S38: Visualization of the Root Mean Square Fluctuation (RMSF)** of the MD simulation of P6. The RMSF represents the flexibility of the protein. The color represents the flexibility from small to high with the corresponding colors from white to yellow to red.

Chimera: Substrate: chloroacetate fluoroacetate

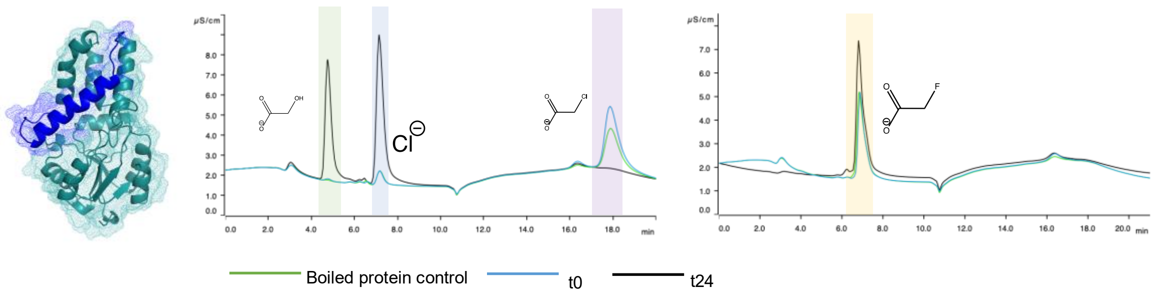

| Protein | substrate | % conversion | concentration glycolate [mM] |
|---|---|---|---|
| P1-P6-blue | chloroacetate 1.3 mM | 75 | 0.98 |
| P1-P6-blue | fluoroacetate 1 mM | 0 | <LOD |

Chimera: Substrate: chloroacetate fluoroacetate

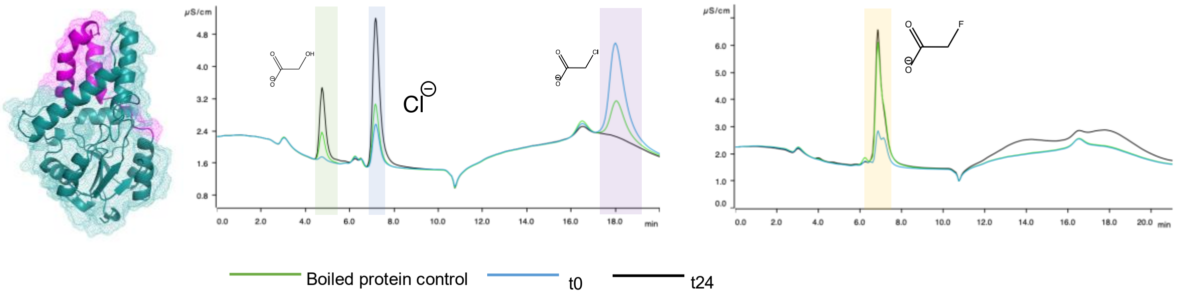

| Protein | substrate | % conversion | concentration glycolate [mM] |
|---|---|---|---|
| P1-P6-pink | chloroacetate 0.6 mM | 68.66 | 0.43 |
| P1-P6-pink | fluoroacetate 1 mM | 0 | <LOD |

Chimera: Substrate: chloroacetate fluoroacetate

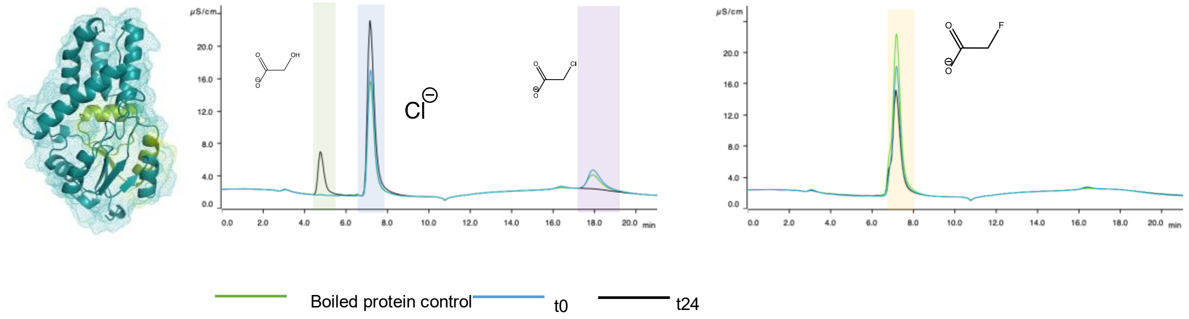

| Protein | substrate | % conversion | concentration glycolate [mM] |
|---|---|---|---|
| P1-P6-green | chloroacetate 1.2 mM | 72.5 | 0.87 |
| P1-P6-green | fluoroacetate 1 mM | 0 | <LOD |

Chimera: Substrate: chloroacetate fluoroacetate

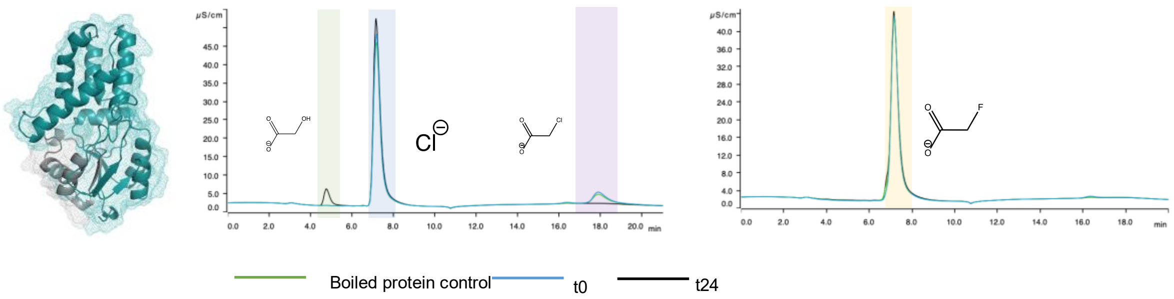

| Protein | substrate | % conversion | concentration glycolate [mM] |
|---|---|---|---|
| P1-P6-green | chloroacetate 1.8 mM | 75.7 | 1.4 |
| P1-P6-green | fluoroacetate 1 mM | 0 | <LOD |

Chimera: Substrate: chloroacetate fluoroacetate

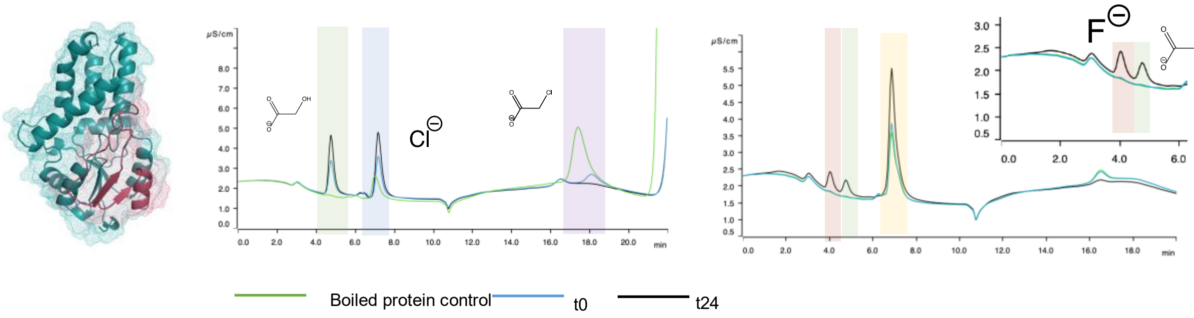

| Protein | substrate | % conversion | concentration glycolate [mM] |
|---|---|---|---|
| P1-P6-red | chloroacetate 1 mM | 84 | 0.84 |
| P1-P6-red | fluoroacetate 1.14 mM | 2.2 | 0.022 |

**Figure S39: Ion chromatograms** of protein in vitro assays with the different chimeric proteins. Tested substrates are fluoroacetate and chloroacetate. Blue: 0h; green: boiled protein control; black: 24h. The turquoise part of the protein structure corresponds to the non-deflourinating protein P1. The colored part of the protein structure corresponds to the homologous region in the defluorinating protein P6.
Chloroacetate shaded in purple, fluoroacetate shaded in yellow, glycolate shaded in green, chloride shaded in blue, and fluoride shaded in red. The tables show the conversion and measured glycolate concentration. Glycolate is used for this calculation due to possible interfering chloride contaminations.

**Supplementary tables:**

**Table S1**: Proteins from the human gut microbiota database (HumGut DB with similarity to the query sequence P5 (PDB ID: 3UMG).

| Query | Subject | % Identity | Alignment length | e-value | Bit score |
|---|---|---|---|---|---|
| 3UMG_1\|Chains | WP_121129595.1 | 46.186 | 236 | 5.50e-74 | 217 |
| 3UMG_1\|Chains | KUG38984.1 | 43.162 | 234 | 2.11e-63 | 191 |
| 3UMG_1\|Chains | NWK50657.1 | 42.308 | 234 | 1.01e-61 | 186 |
| 3UMG_1\|Chains | WP_227153894.1 | 41.818 | 55 | 1.59e-09 | 47.8 |
| 3UMG_1\|Chains | WP_157004912.1 | 41.818 | 55 | 1.61e-09 | 47.8 |
| 3UMG_1\|Chains | WP_087191411.1 | 41.818 | 55 | 1.69e-09 | 47.8 |
| 3UMG_1\|Chains | WP_004693440.1 | 41.453 | 234 | 1.76e-60 | 183 |
| 3UMG_1\|Chains | WP_151836607.1 | 40.773 | 233 | 3.70e-61 | 184 |
| 3UMG_1\|Chains | WP_178618037.1 | 40 | 235 | 5.51e-56 | 171 |
| 3UMG_1\|Chains | WP_118709330.1 | 39.344 | 61 | 1.18e-06 | 39.3 |
| 3UMG_1\|Chains | EEQ56360.1 | 39.224 | 232 | 4.13e-55 | 169 |
| 3UMG_1\|Chains | WP_118709078.1 | 38.462 | 234 | 1.12e-53 | 166 |
| 3UMG_1\|Chains | WP_006686258.1 | 33.6 | 125 | 4.87e-09 | 46.2 |
| 3UMG_1\|Chains | HCH98167.1 | 33.333 | 87 | 2.96e-09 | 47 |
| 3UMG_1\|Chains | PWL57511.1 | 33.333 | 87 | 2.98e-09 | 47 |
| 3UMG_1\|Chains | RGD90338.1 | 32.645 | 242 | 1.25e-42 | 137 |
| 3UMG_1\|Chains | WP_008979552.1 | 30.682 | 88 | 1.67e-09 | 47.8 |
| 3UMG_1\|Chains | WP_008689090.1 | 30.682 | 88 | 1.78e-09 | 47.4 |
| 3UMG_1\|Chains | WP_132223766.1 | 30.682 | 88 | 1.81e-09 | 47.4 |
| 3UMG_1\|Chains | WP_227150888.1 | 30.682 | 88 | 1.90e-09 | 47.4 |
| 3UMG_1\|Chains | MBS5585750.1 | 30.57 | 193 | 7.32e-10 | 48.9 |
| 3UMG_1\|Chains | HIV28856.1 | 30.57 | 193 | 7.53e-10 | 48.9 |
| 3UMG_1\|Chains | ERI71420.1 | 30.208 | 96 | 1.37e-09 | 48.1 |
| 3UMG_1\|Chains | WP_117543986.1 | 30.208 | 96 | 1.44e-09 | 48.1 |
| 3UMG_1\|Chains | GKH53471.1 | 29.487 | 78 | 4.91e-08 | 43.5 |
| 3UMG_1\|Chains | WP_156074201.1 | 29 | 200 | 6.46e-11 | 51.6 |
| 3UMG_1\|Chains | WP_270479031.1 | 29 | 200 | 1.04e-10 | 51.2 |
| 3UMG_1\|Chains | WP_249259624.1 | 27.542 | 236 | 1.53e-11 | 53.5 |
| 3UMG_1\|Chains | WP_036120264.1 | 27.273 | 110 | 1.45e-07 | 42 |
| 3UMG_1\|Chains | WP_027291866.1 | 27.193 | 228 | 3.13e-11 | 52.8 |
| 3UMG_1\|Chains | WP_112144081.1 | 27.136 | 199 | 2.74e-10 | 50.1 |
| 3UMG_1\|Chains | WP_053428405.1 | 26.432 | 227 | 6.59e-17 | 68.9 |
| 3UMG_1\|Chains | WP_027290230.1 | 25.833 | 240 | 2.97e-08 | 43.9 |
| 3UMG_1\|Chains | WP_026089469.1 | 25.652 | 230 | 3.75e-12 | 55.5 |
| 3UMG_1\|Chains | WP_081745164.1 | 25.243 | 206 | 3.40e-14 | 61.2 |
| 3UMG_1\|Chains | WP_069514564.1 | 25 | 208 | 3.81e-09 | 46.6 |
| 3UMG_1\|Chains | WP_102290114.1 | 24.757 | 206 | 1.06e-13 | 59.7 |
| 3UMG_1\|Chains | WP_066861885.1 | 24.309 | 181 | 4.71e-09 | 46.2 |
| 3UMG_1\|Chains | HIW22760.1 | 24.257 | 202 | 5.08e-13 | 57.8 |

| | | | | | |
|---|---|---|---|---|---|
| 3UMG_1\|Chains | WP_263041397.1 | 24.257 | 202 | 7.05e-13 | 57.4 |
| 3UMG_1\|Chains | WP_031546286.1 | 24 | 200 | 5.11e-09 | 46.2 |
| 3UMG_1\|Chains | WP_005951934.1 | 23.81 | 210 | 1.66e-08 | 44.7 |
| 3UMG_1\|Chains | CDD30753.1 | 23.669 | 169 | 7.97e-10 | 48.9 |
| 3UMG_1\|Chains | WP_048310185.1 | 23.478 | 230 | 2.06e-09 | 47.4 |
| 3UMG_1\|Chains | WP_098579072.1 | 23.394 | 218 | 3.01e-10 | 49.7 |
| 3UMG_1\|Chains | WP_050688410.1 | 23.394 | 218 | 5.37e-10 | 48.9 |
| 3UMG_1\|Chains | WP_064471359.1 | 23.394 | 218 | 5.58e-10 | 48.9 |
| 3UMG_1\|Chains | WP_107514404.1 | 22.984 | 248 | 2.37e-13 | 58.9 |
| 3UMG_1\|Chains | CDD29996.1 | 22.772 | 202 | 3.46e-12 | 55.5 |
| 3UMG_1\|Chains | WP_065338764.1 | 22.5 | 240 | 6.96e-14 | 60.5 |
| 3UMG_1\|Chains | WP_034355080.1 | 22.326 | 215 | 5.19e-09 | 46.2 |
| 3UMG_1\|Chains | WP_034377360.1 | 22.326 | 215 | 5.39e-09 | 46.2 |
| 3UMG_1\|Chains | WP_053359908.1 | 22.273 | 220 | 1.38e-10 | 50.8 |
| 3UMG_1\|Chains | WP_036125194.1 | 22.222 | 198 | 9.13e-10 | 48.5 |
| 3UMG_1\|Chains | WP_187031722.1 | 22.176 | 239 | 5.65e-12 | 55.1 |
| 3UMG_1\|Chains | WP_070051353.1 | 21.951 | 205 | 2.67e-13 | 58.5 |
| 3UMG_1\|Chains | WP_029378231.1 | 21.635 | 208 | 4.55e-13 | 58.2 |
| 3UMG_1\|Chains | WP_233640236.1 | 21.635 | 208 | 5.07e-13 | 57.8 |
| 3UMG_1\|Chains | WP_009253081.1 | 21.296 | 216 | 2.99e-09 | 47 |
| 3UMG_1\|Chains | WP_107597559.1 | 19.917 | 241 | 4.94e-10 | 49.3 |
| 3UMG_1\|Chains | GKG91607.1 | 17.961 | 206 | 3.14e-04 | 32 |
| 3UMG_1\|Chains | WP_087189991.1 | 17.961 | 206 | 3.27e-04 | 32 |

**Table S2:** Proteins selected for in vitro work. The previously characterized L-2 haloacid dehalogenase P5 (PDB ID: 3UMG) was included as a positive control.

| **NCBI identifier** | | **NCBI taxonomy** | **HumGut DB taxonomy** |
|---|---|---|---|
| WP_118709078.1 | P1 | *Enterocloster aldenensis* | *Faecalibacterium prausnitzii* |
| GKG91607.1 | P2 | MULTISPECIES: Gordonibacter | *Gordonibacter pamelaeae* |
| WP_087189991.1 | P3 | *Gordonibacter urolithinfaciens* | *Gordonibacter pamelaeae* |
| WP_004693440.1 | P4 | *Acinetobacter johnsonii* | *Acinetobacter johnsonii* |
| WP_011593529.1 | P5 | *Rhodococcus jostii* RHA1 | Not in HumGut DB (soil bacterium) |
| WP_178618037.1 | P6 | *Guopingia tenuis* | *Catabacter* sp. UBA5399 |
| WP_006686258.1 | P7 | *Citrobacter youngae* | *Citrobacter youngae* |
| WP_053428405.1 | P8 | *Rossellomorea marisflavi* | *Rossellomorea marisflavi* |

**Table S3**: Genome neighborhood context of 55 bioinformatically-identified HADs in this study. The second most abundant PFAM is PF13419, referrs to the PFAM of the HAD queries. *Number of hits summed within 55 HAD genome neighborhoods **Frequency of the PFAM (per neighborhood) within ± 6 genes of the query HAD

| ID | PFAM | Number | and descriptions of hits (for 55 HAD neighbourhoods) | Frequency* |
|---|---|---|---|---|
| 1 | - | 169 | NA/hypothetical proteins | 1.352 |
| 2 | PF13419 | 49 | Haloacid dehalogenase-like hydrolase | 0.392 |
| 3 | PF00005 | 14 | ABC transporter | 0.112 |
| 4 | PF12833 | 11 | Helix-turn-helix domain | 0.088 |
| 5 | PF01380 | 9 | SIS domain | 0.072 |
| 6 | PF00528 | 8 | Binding-protein-dependent transport system inner membrane component | 0.064 |
| 7 | PF01019 | 8 | Gamma-glutamyltranspeptidase | 0.064 |
| 8 | PF01261 | 8 | Xylose isomerase-like TIM barrel | 0.064 |
| 9 | PF01381 | 8 | Helix-turn-helix | 0.064 |
| 10 | PF00474 | 7 | Sodium:solute symporter family | 0.056 |
| 11 | PF00480 | 6 | ROK family | 0.048 |
| 12 | PF00583 | 6 | Acetyltransferase (GNAT) family | 0.048 |
| 13 | PF00701 | 6 | Dihydrodipicolinate synthetase family | 0.048 |
| 14 | PF03180 | 6 | NlpA lipoprotein | 0.048 |
| 15 | PF04203 | 6 | Sortase domain | 0.048 |
| 16 | PF00248 | 5 | Aldo/keto reductase family | 0.04 |
| 17 | PF04131 | 5 | Putative N-acetylmannosamine-6-phosphate epimerase | 0.04 |
| 18 | PF04464 | 5 | CDP-Glycerol:Poly(glycerophosphate) glycerophosphotransferase | 0.04 |
| 19 | PF09587 | 5 | Bacterial capsule synthesis protein PGA_cap | 0.04 |
| 20 | PF00072 | 4 | Response regulator receiver domain | 0.032 |
| 21 | PF00117 | 4 | Glutamine amidotransferase class-I | 0.032 |
| 22 | PF00535 | 4 | Glycosyl transferase family 2 | 0.032 |
| 23 | PF00664 | 4 | ABC transporter transmembrane region | 0.032 |
| 24 | PF00860 | 4 | Permease family | 0.032 |
| 25 | PF01022 | 4 | Bacterial regulatory protein, arsR family | 0.032 |

| | | | | |
|---|---|---|---|---|
| 26 | PF01451 | 4 | Low molecular weight phosphotyrosine protein phosphatase | 0.032 |
| 27 | PF01743 | 4 | Poly A polymerase head domain | 0.032 |
| 28 | PF02378 | 4 | Phosphotransferase system, EIIC | 0.032 |
| 29 | PF02417 | 4 | Chromate transporter | 0.032 |
| 30 | PF02436 | 4 | Conserved carboxylase domain | 0.032 |
| 31 | PF02518 | 4 | Histidine kinase-, DNA gyrase B-, and HSP90-like ATPase | 0.032 |
| 32 | PF02687 | 4 | FtsX-like permease family | 0.032 |
| 33 | PF03358 | 4 | NADPH-dependent FMN reductase | 0.032 |
| 34 | PF03466 | 4 | LysR substrate binding domain | 0.032 |
| 35 | PF06962 | 4 | Putative rRNA methylase | 0.032 |
| 36 | PF08028 | 4 | Acyl-CoA dehydrogenase, C-terminal domain | 0.032 |
| 37 | PF08240 | 4 | Alcohol dehydrogenase GroES-like domain | 0.032 |
| 38 | PF08282 | 4 | haloacid dehalogenase-like hydrolase | 0.032 |
| 39 | PF08840 | 4 | BAAT / Acyl-CoA thioester hydrolase C terminal | 0.032 |
| 40 | PF09359 | 4 | VTC domain | 0.032 |
| 41 | PF13192 | 4 | Thioredoxin domain | 0.032 |
| 42 | PF13302 | 4 | Acetyltransferase (GNAT) domain | 0.032 |
| 43 | PF13338 | 4 | Transcriptional regulator, AbiEi antitoxin | 0.032 |
| 44 | PF14262 | 4 | Carbohydrate-binding domain-containing protein Cthe_2159 | 0.032 |
| 45 | PF16316 | 4 | Domain of unknown function (DUF4956) | 0.032 |

**Table S5:** Chemicals used in this study

| Chemical | Molecular structure | Abbr. | Supplier | CAS | Purity |
|---|---|---|---|---|---|
| chloroacetic acid | 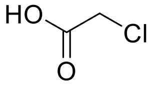 | ClAc | Sigma Aldrich | 3926-62-3 | 98 % |
| fluoroacetic acid | 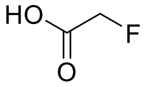 | FAc | Sigma Aldrich, abcr | 144-49-0 | 98 % |
| 2,2-difluoroacetic acid | 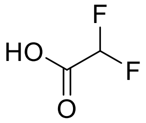 | DFA | Sigma Aldrich | 381-73-7 | 98 % |
| trifluoroacetic acid | 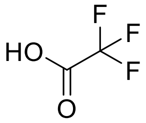 | TFA | Sigma Aldrich | 2923-18-4 | 98 % |
| 2-fluoropropionic acid | 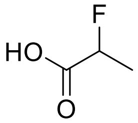 | FP | Sigma Aldrich | 6087-13-4 | 97 % |
| 3-amino-2-fluoropropionic acid | 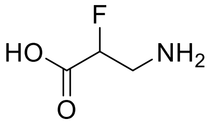 | FBAL | Sigma Aldrich | 3821-81-6 | 95 % |
| 2-2-difluoropropionic acid | 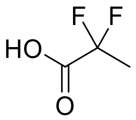 | DFA | Sigma Aldrich | 373-96-6 | 97 % |
| 3,3,3-trifluoropropionic acid | 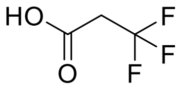 | TFP | Sigma Aldrich | 2516-99-6 | 98 % |
| 5,5,5-trifluoropentanoic acid | 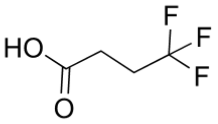 | TFB | Sigma Aldrich | 406-93-9 | 97% |
| heptafluorobutyric acid | 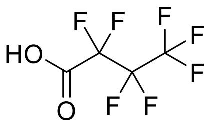 | HFB | Sigma Aldrich | 375-22-4 | 98% |

**Table S4:** Defluorination data from *Gordonibacter pamelaeae* DSM 110924 in GMM medium. Incubated with 10 mM fluoroacetate for 24h. Std: standard, Gran linearization with slope index value -59.16. Measured with a Metrohm fluoride ion selective electrode (ISE).

| Sample name | Standard added [mL] | [mV] | Gran linearization | Statistics | $F^-$ conc [mg |
|---|---|---|---|---|---|
| *G. pamelaeae* in GMM-1 | 0 | 218.5 | 0.008103751 | a: 0.121 | 0.24 |
| | 0.2 | 187.1 | 0.027644974 | b: 0.006 | |
| | 0.2 | 170.4 | 0.053217788 | r2: 0.998 | |
| | 0.2 | 160.6 | 0.078316624 | | |
| | 0.2 | 153.6 | 0.10335019 | | |
| *G. pamelaeae* in GMM-2 | 0 | 217.1 | 0.008557575 | a: 0.118 | 0.28 |
| | 0.2 | 186.3 | 0.028519298 | b: 0.007 | |
| | 0.2 | 170.8 | 0.052395682 | r2: 0.998 | |
| | 0.2 | 160.8 | 0.077709353 | | |
| | 0.2 | 153.9 | 0.10215045 | | |
| *G. pamelaeae* in GMM-3 | 0 | 220.8 | 0.007409834 | a: 0.124 | 0.18 |
| | 0.2 | 187.2 | 0.027537585 | b: 0.004 | |
| | 0.4 | 161.2 | 0.076508902 | r2: 0.999 | |
| | 0.6 | 143.6 | 0.15402154 | | |
| | 0.8 | 131.3 | 0.253421984 | | |
| *G. pamelaeae* in GMM + 10mM FAc-1 | 0 | 171.4 | 0.050679478 | a: 0.124 | 2.03 |
| | 0.2 | 161.6 | 0.074584857 | b: 0.050 | |
| | 0.2 | 154.1 | 0.100364662 | r2: 0.999 | |
| | 0.2 | 148.5 | 0.125424982 | | |
| | 0.2 | 144.1 | 0.149586621 | | |
| *G. pamelaeae* in GMM + 10mM FAc-2 | 0 | 170.7 | 0.052079218 | a: 0.128 | 2.03 |
| | 0.2 | 160.7 | 0.077243803 | b: 0.052 | |
| | 0.2 | 153.5 | 0.102736038 | r2 : 0.999 | |
| | 0.2 | 148 | 0.127889739 | | |
| | 0.2 | 143.3 | 0.15431758 | | |
| *G. pamelaeae* in GMM + 10mM FAc-3 | 0 | 171.6 | 0.050286507 | a: 0.122 | 2.00 |
| | 0.2 | 161.8 | 0.074006523 | b: 0.049 | |
| | 0.4 | 149.9 | 0.11877346 | r2: 0.999 | |
| | 0.6 | 137.4 | 0.196056842 | | |
| | 0.8 | 127.5 | 0.293816971 | | |
| 10mM FAc in GMM-1 | 0 | 169.7 | 0.054146173 | a: 0.131 | 2.05 |
| | 0.2 | 160 | 0.079377235 | b: 0.054 | |
| | 0.2 | 152.7 | 0.105985258 | r2: 0.999 | |
| | 0.2 | 147.1 | 0.132449 | | |
| | 0.2 | 142.6 | 0.158579748 | | |
| 10mM FAc in GMM-2 | 0 | 171 | 0.05250415 | a: 0.111 | 2.13 |
| | 0.2 | 165.8 | 0.064597328 | b: 0.047 | |
| | 0.2 | 157.7 | 0.088970237 | r2: 0.987 | |
| | 0.2 | 151.5 | 0.113801612 | | |
| | 0.2 | 146.5 | 0.138917823 | | |

**Table S6:** Kinetic parameters for the P3 haloacid dehalogenase (this study) compared to known haloacid dehalogenases DeHa2 and fluoroacetate dehalogenases DeHa4 reported by Farajollahi et al., 2024 (1) and RPA1163 reported by Mehrabi et al., 2019 (2). RPA1163 and P3 were assayed at the same pH (8.5) while DeHa2 and DeHa4 were measured at pH 7.0. The $K_M$ and the catalytic efficiency for DeHa2 and DeHa4 is lower compared to P3 and RPA1163. FAc = fluoroacetate.

| **Enzyme** | **Substrate** | **pH** | **$K_m$ (mM)** | **$k_{cat}$ ($s^{-1}$)** | **$k_{cat}/K_m$ ($s^{-1}$ $mM^{-1}$)** |
|---|---|---|---|---|---|
| P3 (this study) | FAc | 8.5 | 59.8 ± 7.7 | 0.25 ± 0.02 | 4.24 ± 0.08 |
| RPA1163 | FAc | 8.5 | 86.8 ± 13.7 | 1.07 ± 0.11 | 12.3 ± 2.3 |
| DeHa2 | FAc | 7 | 5.47 ± 1.54 | $(8.5±1.2)×10^{-4}$ | $(1.8±0.2)×10^{-4}$ |
| DeHa4 | FAc | 7 | 2.19 ± 0.37 | $(2.1±0.22)×10^{-4}$ | $(1.056±0.011)×10^{-4}$ |

**Table S7**: Proteins with available crystal structures used in this study for crystal structure AlphaFold comparison, comparison of MD simulations and Cα calculations.

| **Protein** | **PDB** | **organism** | **defluorination** |
|---|---|---|---|
| LDexYL | 1JUD | *Pseudomonas* sp. YL | No |
| DhlB | 1AQ6 | *Xanthobacter autotrophicus* | No |
| Rsc1362 | 3UMB | *Ralstonia solanacearum* | No |
| PA0810 | 3UMC | *Pseudomonas aeruginosa* | No |
| Rha0230 | 3UMG | *Rhodococcus jostii* RHA1 | yes |
| DehIVA | 2NO4 | *Burkholderia cepacia* | No |

**Table S8**: Conserved residue numbers and $C_\alpha$ atom indices used for the distance analysis between defluorinating and non-defluorinating HADs.

Residue number of conserved residues

| Protein ID | D1 | T | R1 | N | K | D2 | R2 |
|---|---|---|---|---|---|---|---|
| DehIVa | 20 | 24 | 51 | 129 | 161 | 190 | 206 |
| DhlB | 8 | 12 | 39 | 115 | 147 | 176 | 192 |
| GKG | 22 | 26 | 53 | 129 | 161 | 190 | 206 |
| PA0810 | 28 | 32 | 65 | 143 | 173 | 202 | 218 |
| Rha0230 | 21 | 25 | 58 | 139 | 169 | 198 | 214 |
| RSc1362 | 10 | 14 | 41 | 123 | 155 | 184 | 200 |
| WP087 | 8 | 12 | 39 | 115 | 147 | 176 | 192 |
| WP118 | 12 | 16 | 49 | 129 | 159 | 188 | 204 |
| WP178 | 12 | 16 | 49 | 128 | 158 | 187 | 203 |

$C_\alpha$ Atom Index

| Protein ID | D1 Cα | T Cα | R1 Cα | N Cα | K Cα | D2 Cα | R2 Cα |
|---|---|---|---|---|---|---|---|
| DehIVa | 305 | 355 | 750 | 2043 | 2531 | 2983 | 3215 |
| DhlB | 125 | 175 | 594 | 1800 | 2260 | 2691 | 2913 |
| GKG | 296 | 346 | 768 | 2022 | 2465 | 2940 | 3168 |
| PA0810 | 428 | 483 | 1019 | 2282 | 2733 | 3184 | 3431 |
| Rha0230 | 313 | 366 | 871 | 2133 | 2595 | 3023 | 3248 |
| RSc1362 | 152 | 202 | 620 | 1929 | 2408 | 2831 | 3068 |
| WP087 | 115 | 165 | 587 | 1841 | 2287 | 2762 | 2990 |
| WP118 | 203 | 256 | 760 | 2036 | 2506 | 2951 | 3177 |
| WP178 | 189 | 242 | 752 | 2102 | 2584 | 3015 | 3237 |

**Table S9:** Phylogenetic tree tip indices and UniProt accessions for the HAD-like superfamily enzymes included in the maximum-likelihood phylogenetic tree (Figure S20).

| **Index** | **UniProt ID** | **Index** | **UniProt ID** | **Index** | **UniProt** |
|---|---|---|---|---|---|
| 1 | A0A099PBW3 | 84 | P42941 | 167 | A0A2R8Y6K4 |
| 2 | A0A926DHK7 (P6) | 85 | P9WGJ3 | 168 | A0A2R8YET7 |

| | | | | | |
|---|---|---|---|---|---|
| 3 | Q51645 | 86 | Q12A06 | 169 | A0A2R8YGM9 |
| 4 | Q53464 | 87 | Q21YU0 | 170 | A0A8M2B9H5 |
| 5 | Q60099 | 88 | Q5M3B3 | 171 | A0A8M2BHD8 |
| 6 | Q8XZN3 | 89 | Q5QXU4 | 172 | A0A8M2BHD9 |
| 7 | Q9I5C9 | 90 | Q7M7U5 | 173 | A0A8M2BHG2 |
| 8 | A0A423UMR7 | 91 | Q9CHW3 | 174 | A0A8M2BHU7 |
| 9 | Q0SK70 (P5) | 92 | Q9KPM2 | 175 | A0A8M3AP55 |
| 10 | Q122Z0 | 93 | Q9S281 | 176 | A0A8M6YVK2 |
| 11 | Q12G50 | 94 | P60487 | 177 | A0JME1 |
| 12 | Q2IG66 | 95 | Q3ZBF9 | 178 | A6NCB9 |
| 13 | Q96XE7 | 96 | Q8VD52 | 179 | B1APR7 |
| 14 | Q94K71 | 97 | Q96GD0 | 180 | E7EQM5 |
| 15 | P95649 | 98 | P21829 | 181 | E7ETN2 |
| 16 | Q88M11 | 99 | P27848 | 182 | F8WB53 |
| 17 | Q9HZ62 | 100 | Q8TCD6 | 183 | O94279 |
| 18 | A4QFW4 | 101 | Q9D9M5 | 184 | P40081 |
| 19 | Q15JF8 | 102 | P71447 | 185 | Q6P4T3 |
| 20 | F4KFT7 | 103 | P77366 | 186 | Q86V88 |
| 21 | Q9SU92 | 104 | O06995 | 187 | A0A1W2P721 |
| 22 | Q8A947 | 105 | P0A8Y3 | 188 | A0A804HHY3 |
| 23 | P53981 | 106 | P77625 | 189 | A0A804HL00 |
| 24 | Q9P6N2 | 107 | O58690 | 190 | A0A8C8KDW0 |
| 25 | F4JTE7 | 108 | P76329 | 191 | A2A953 |
| 26 | P40106 | 109 | O31156 | 192 | B1AKW3 |
| 27 | P41277 | 110 | Q9I433 | 193 | B7QN98 |
| 28 | Q8VZP1 | 111 | Q7ZAP3 | 194 | E5RHZ7 |
| 29 | O33194 | 112 | D0VWZ7 | 195 | E7ESD5 |

| | | | | | |
|---|---|---|---|---|---|
| 30 | O07565 | 113 | Q5XAQ1 | 196 | E9PLN6 |
| 31 | Q84VZ1 | 114 | Q8R2H9 | 197 | F2Z2Y1 |
| 32 | A0A2K3DU55 | 115 | Q8TCT1 | 198 | Q8BY78 |
| 33 | Q9LXR9 | 116 | Q9FN41 | 199 | Q8H684 |
| 34 | C0QRP9 | 117 | O31667 | 200 | P9WPI9 |
| 35 | Q12XX5 | 118 | P0DV34 | 201 | Q9HTR2 |
| 36 | Q86ZR7 | 119 | Q9I6F6 | 202 | P0A8Y5 |
| 37 | A0A0U5GNT1 | 120 | P32662 | 203 | P75792 |
| 38 | P46891 | 121 | Q9HLQ2 | 204 | Q8VZ10 |
| 39 | P0ADP0 | 122 | O67359 | 205 | Q9ZVJ5 |
| 40 | P75809 | 123 | P0DKC3 | 206 | P31467 |
| 41 | P42962 | 124 | P40852 | 207 | P77475 |
| 42 | Q9LDD5 | 125 | P40853 | 208 | J9VX38 |
| 43 | P70947 | 126 | Q0I1W8 | 209 | P94592 |
| 44 | P96741 | 127 | Q8GWU0 | 210 | Q8RYE9 |
| 45 | F1RIU1 | 128 | Q6EP66 | 211 | Q9X0Y1 |
| 46 | Q498V7 | 129 | Q84MD8 | 212 | A0A061ACH4 |
| 47 | A0A8V1ALE8 | 130 | Q8TBE9 | 213 | A0A061ACK7 |
| 48 | C9JBI3 | 131 | Q5M969 | 214 | A0A061ADS3 |
| 49 | E1C9F4 | 132 | Q9CPT3 | 215 | A0A061AKN1 |
| 50 | O25367 | 133 | P0A8Y1 | 216 | B3DI21 |
| 51 | Q8YI30 | 134 | O32125 | 217 | B7PHY5 |
| 52 | Q8Z598 | 135 | P0AF24 | 218 | F1QGU0 |
| 53 | V7LE91 | 136 | P64636 | 219 | O06480 |
| 54 | Q67YC0 | 137 | Q8DPQ3 | 220 | O59346 |
| 55 | Q9H008 | 138 | Q9I767 | 221 | P44447 |
| 56 | Q0VD18 | 139 | Q52087 | 222 | P9WGJ1 |

| | | | | | |
|---|---|---|---|---|---|
| 57 | Q9D7I5 | 140 | P60527 | 223 | P9WMS5 |
| 58 | Q9JMQ2 | 141 | O06652 | 224 | Q12486 |
| 59 | A0A8V1AB74 | 142 | B2Z3V8 | 225 | Q2QQ95 |
| 60 | P77247 | 143 | R4GRT2 | 226 | Q2R0W7 |
| 61 | P38773 | 144 | Q01399 | 227 | Q2UEK4 |
| 62 | P38774 | 145 | Q08623 | 228 | Q2UEL0 |
| 63 | Q52086 | 146 | Q9D5U5 | 229 | Q5XD45 |
| 64 | Q8KLS9 | 147 | A6NDG6 | 230 | Q836C7 |
| 65 | A0A1P8BAY5 | 148 | D3ZDK7 | 231 | Q8E044 |
| 66 | P32626 | 149 | Q8CHP8 | 232 | Q8GAX4 |
| 67 | Q9UHY7 | 150 | P97767 | 233 | Q95Q10 |
| 68 | A0A8I6ADH3 | 151 | Q99502 | 234 | Q9U1W6 |
| 69 | Q8BGB7 | 152 | O00167 | 235 | Q9U1W7 |
| 70 | A0A1L1RQM2 | 153 | O08575 | 236 | Q9UTA6 |
| 71 | A0A8V0XML6 | 154 | O17670 | 237 | Q9VYT0 |
| 72 | P0C8L5 | 155 | O82162 | 238 | B6YTD6 |
| 73 | O53289 | 156 | O95677 | 239 | B7Q9S3 |
| 74 | O82796 | 157 | P97480 | 240 | G0L7V6 |
| 75 | P0AGB0 | 158 | Q05201 | 241 | O58216 |
| 76 | P78330 | 159 | Q99504 | 242 | Q7A484 |
| 77 | A0QJI1 | 160 | Q9D967 | 243 | P1 |
| 78 | P94512 | 161 | A0A0G2JZ01 | 244 | P2 |
| 79 | P94526 | 162 | A0A8I6A077 | 245 | P3 |
| 80 | Q58989 | 163 | A0A8I6AMN9 | 246 | P4 |
| 81 | Q72H00 | 164 | A0A8M3AZ17 | 247 | P7 |
| 82 | Q99LS3 | 165 | Q6DBY3 | 248 | P8 |
| 83 | A7H590 | 166 | Q9W6E8 | | |

**Table S10:** Comparison betweens fluoride concentrations measured by ion chromatography and measured by colorimetric assay. The samples were taken from the alanine scanning experiment (1mM fluoroacetate, 24h incubation at 37°C).

| | P6_R99A | P6_L100A | P6_W101A | P6_H102A | P6_R103A | P6_L104A | P6_P107A | P6_P108A | P6_D109A | P6_V110A |
|---|---|---|---|---|---|---|---|---|---|---|
| **Colorimetric µM** | 62 | 108 | 116 | 3 | 5 | 10 | 34 | 3 | 108 | 3 |
| **Ion Chromat. µM** | 79 | 132 | 149 | 11 | 17 | 27 | 51 | 11 | 139 | 12 |
| **Standard deviation** | 10 | 24 | 33 | 8 | 12 | 17 | 17 | 8 | 31 | 9 |

| | P6_Q111A | P6_E112A | P6_G113A | P6_K118A | P6_T119A | P6_Y121A | P6_S122A |
|---|---|---|---|---|---|---|---|
| **Colorimetric µM** | 84 | 108 | 116 | 3 | 105 | 86 | 3 |
| **Ion Chromat. µM** | 108 | 179 | 120 | 14 | 122 | 128 | 14 |
| **Standard deviation** | 24 | 71 | 4 | 11 | 17 | 42 | 6 |

**Table S11:** Interfering substances and buffers for the colorimetric fluoride assay. To solutions of 0 and 100 µM fluoride the indicated concentrations of compound were added. Then it was measured if there was a change in absorption compared to the control without the compound.

| **Compound and concentration** | **Interference** |
|---|---|
| 13 g/liter nutrient broth | yes |
| 0.1% yeast extract | yes |
| 37 g/liter brain heart infusion | yes |
| > 500 µM sodium phosphate monobasic | yes |
| 25% acetone | no |
| 300 mM acetate buffer | no |
| 20 mM HEPES | no |
| 10% glycerol | no |
| 100 mM sodium chloride | no |
| 14 mM sodium bromide | no |

| | |
|---|---|
| 500 µM sodium sulfate | no |
| 10 mM sodium nitrate | no |
| 47 µM borate (borax) | no |
| 2 µM EDTA | no |
| 0.1% Casamino acids | no |
| 50 mM succinic acid | no |
| 100 mM magnesium chloride | no |
| 200 mM calcium chloride | no |
| 2 mM lithium chloride | no |
| 2% glucose | no |
| 2% arabinose | no |
| 2% sucrose | no |
| 1 mM thiamine hydrochloride | no |
| 1 mM biotin | no |
| 0.2 M Tris | no |
| 0.2 M MOPS | no |
| 20 mM ferric chloride | no |
| 20% ethanol | no |
| 100 mM MES | no |
| 20% DMSO | no |
| 100 mM imidazole | no |
| 100 mM zinc chloride | no |
| 1 mM cupric chloride | no |

| | |
|---|---|
| 10 µM sodium citrate dihydrate | no |
| 2.5 mM ammonium chloride | no |
| 500 mM potassium chloride | no |
| 2% DMF | no |
| 20% acetonitrile | no |
| 10 mM Tricine | no |
| <500 µM sodium phosphate monobasic | no |

**Table S12:** Additional HADs included in this study used for colorimetric assay development.

| Protein id | NCBI identifier | Organism | Defluorination |
|---|---|---|---|
| P9 | WP_050688410.1 | *Priestia megaterium* | Yes |
| P10 | WP_029378231.1 | *Staphylococcus xylosus* | No |
| P11 | WP_048310185.1 | *Alkalihalobacillus macyae* | No |
| P12 | WP_081745164.1 | *Eisenbergiella massiliensis* | No |
| P13 | WP_066861885.1 | *Neglectibacter timonensis* | No |
| P14 | WP_263041397.1 | *Acutalibacter* LFL 21 | No |
| P15 | WP_187031722.1 | *Oscillospiraceae* | No |
| P16 | WP_036125194.1 | *Lysinibacillus* | No |
| P17 | WP_031546286.1 | *Lacticaseibacillus rhamnosus* | No |
| P18 | WP_053359908.1 | *Clostridium butyricum* | No |
| P19 | GKH53471 | *Lachnospiraceae bacterium* | No |

**Table S13**: P6 variants are showing reduced defluorination vs. dechlorination activity in both experiment 1 and experiment 2.

| **Top P6 Protein variant influencing defluorination in both experiment 1 and 2** | | |
|---|---|---|
| V17A | E51A | A218H |
| V24A | Y53A | E223A |
| G28A | K55A | V225A |
| S38A | E59A | A232H |
| D40A | G63A | G236A |
| E43A | G129A | |
| N46A | F157A | |

**Table S14:** Top 48 P6 variants showing reduced defluorination activity in the alanine scanning experiment 1. The values correspond to the difference between dechlorination activity minus defluorination activity. The values were min-max normalized.

| value | P6 variant | value | P6 variant | value | P6 variant | value | P6 variant |
|---|---|---|---|---|---|---|---|
| 1.00 | gY53A | 0.70 | gG236A | 0.62 | gT224A | 0.55 | gA226H |
| 0.83 | gR229A | 0.69 | gA233H | 0.62 | gG63A | 0.54 | gD40A |
| 0.80 | gA209H | 0.69 | gA232H | 0.61 | gK55A | 0.53 | gg29A |
| 0.78 | gV225A | 0.68 | gV210A | 0.60 | gP204A | 0.53 | gK219A |
| 0.77 | gG28A | 0.67 | gE22A | 0.60 | gK35A | 0.53 | gR103A |
| 0.75 | gS38A | 0.66 | gR234A | 0.59 | gL199A | 0.53 | gE51A |
| 0.73 | gN46A | 0.65 | gE223A | 0.59 | gP106A | 0.52 | gF157A |
| 0.72 | gL117A | 0.65 | gT94A | 0.59 | gV24A | 0.52 | gV237A |
| 0.72 | gP125A | 0.63 | gP202A | 0.58 | gg27A | 0.52 | gg43A |
| 0.72 | gE217A | 0.63 | gA218H | 0.58 | gG129A | 0.52 | gP197A |
| 0.71 | gV214A | 0.63 | gD221A | 0.57 | gG52A | 0.51 | gE59A |
| 0.71 | gG36A | 0.63 | gL133A | 0.56 | gN212A | 0.51 | gD71A |

**Table S15:** Top 48 P6 variants showing reduced defluorination activity in the alanine screening experiment 2. The values correspond to dechlorination activity minus defluorination activity normalized with protein concentrations and to wild type protein (%).

| value | P6 variant | value | P6 variant | value | P6 variant | value | P6 variant |
|---|---|---|---|---|---|---|---|
| 1.314 | gY53A | 0.539 | gQ26A | 0.378 | gL87A | 0.304 | gV37A |
| 0.897 | gV24A | 0.516 | gG236A | 0.376 | gG129A | 0.296 | gM57A |
| 0.861 | gS38A | 0.506 | gS4A | 0.373 | gV3A | 0.291 | gA218H |
| 0.784 | gV17A | 0.469 | gD40A | 0.373 | gH213A | 0.274 | gE215A |
| 0.747 | gR75A | 0.461 | gG28A | 0.355 | gI10A | 0.269 | gL104A |

| | | | | | | | |
|---|---|---|---|---|---|---|---|
| 0.741 | gI39A | 0.456 | gE59A | 0.355 | gE91A | 0.253 | gT90A |
| 0.668 | gL89A | 0.432 | gK88A | 0.347 | gK55A | 0.251 | gF157A |
| 0.665 | gD2A | 0.426 | gV18A | 0.344 | gA232H | 0.251 | gN46A |
| 0.658 | gE93A | 0.408 | gV56A | 0.344 | gY220A | 0.242 | gV6A |
| 0.601 | gI194A | 0.403 | gK77A | 0.336 | gF228A | 0.240 | gE50A |
| 0.599 | gR7A | 0.385 | gA30H | 0.330 | gE43A | 0.228 | gV25A |
| 0.551 | gH42A | 0.382 | gE51A | 0.309 | gV225A | 0.220 | gG63A |

**Table S16:** Random forest classification accuracy based on 1000 distinct models with 80%/20% training test splits for full-length protein features and C-terminal features (final 41 amino acids). Experiment 1 was conducted on data without protein concentration normalization and without case weighting. Experiment 2 was conducted after dilution 1:3, with normalization for protein concentration and with case weighting. For this reason datasets were not combined however both yielded comparable results.

| | **Full-length protein model** | **C-terminus only model** |
|---|---|---|
| **Experiment 1 (min-max normalized)** | | |
| Training accuracy | 0.948 ± 0.008 | 0.832 ± 0.023 |
| Testing accuracy | 0.948 ± 0.020 | 0.832 ± 0.078 |
| Testing-F1 score | 0.928 ± 0.028 | 0.728 ± 0.129 |
| **Experiment 2** | **(diluted with protein concentration normalized)** | |
| | **Full-length protein model** | **C-terminus only model** |
| Training accuracy | 0.877 ± 0.003 | 0.800 ± 0.021 |
| Testing accuracy | 0.863 ± 0.006 | 0.791 ± 0.033 |

**Table S17:** Five HADs used for machine learning validation highlighting confident defluorination prediction for proteins with high defluorination prediction probabilities (P20) compared to non-defluorination proteins with low prediction probabilities (P21-P24). *Defluorination probability predicted by random forest (see Methods).

| **Protein id** | **NCBI identifier** | **Organism** | **Defluorination** | **Defluorination probability*** |
|---|---|---|---|---|
| P20 | WP_139652913.1 | *Raoultibacter phocaeensis* | Yes | 0.85 ± 0.19 |
| P21 | HTS94301.1 | *Stellaceae bacterium* | No | 0.54 ± 0.09 |
| P22 | PYN88313.1 | *Ca. Rokuibacteriota* | No | 0.43 ± 0.06 |
| P23 | MDH3205452.1 | *Gemmatimonadota bacterium* | No | 0.57 ± 0.10 |
| P24 | MBS6900484.1 | *Eubacterium sp.* | No | 0.38 ± 0.06 |

**Table S18:** Defluorination data from the alanine scanning library used for machine learning and Figure 4D. exp1 = Experiment 1, exp2 = Experiment 2, sd = standard deviation.

| label | mean exp1 | sd exp1 | mean exp2 | sd exp2 | label | mean exp1 | sd exp1 | mean exp2 | sd exp2 |
|---|---|---|---|---|---|---|---|---|---|
| gD2A | 0.4136 | 0.2258 | 0.1213 | 0.016 | gK120A | 0.5491 | 0.1361 | 0.2402 | 0.056 |
| gV3A | 0.1457 | 0.0639 | 0.0515 | 0.0324 | gY121A | 0.1511 | 0.1288 | 0.0021 | 0.0014 |
| gS4A | 0.1973 | 0.1272 | 0.2561 | 0.0631 | gS122A | 0.8013 | 0.0668 | 0.8107 | 0.1928 |
| gN5A | 0.2757 | 0.1397 | 0.1303 | 0.0359 | gI123A | 0.1079 | 0.0932 | 0.0048 | 0.003 |
| gV6A | 0.1922 | 0.0691 | 0.0346 | 0.0186 | gG124A | 0.1994 | 0.0819 | 0.0698 | 0.0314 |
| gR7A | 0.3105 | 0.0559 | 0.0788 | 0.0227 | gP125A | 0.3274 | 0.0329 | 0.1544 | 0.0486 |
| gI8A | 0.3276 | 0.1364 | 0.0355 | 0.0168 | gF126A | 0.1163 | 0.1645 | 0.0049 | 0.0032 |
| gV9A | 0.2828 | 0.1080 | 0.0114 | 0.0041 | gS127A | 0.1837 | 0.0787 | 0.0118 | 0.0076 |
| gI10A | 0.3694 | 0.1784 | 0.0862 | 0.0221 | gN128A | 0.2146 | 0.1137 | 0.0051 | 0.0012 |
| gF11A | 0.1792 | 0.0538 | 0.0063 | 0.0022 | gG129A | 0.3259 | 0.1466 | 0.097 | 0.006 |
| gD12A | 0.1713 | 0.1767 | 0.0222 | 0.0393 | gD130A | 0.2204 | 0.0153 | 0.0043 | 0.0036 |
| gT13A | 0.4883 | 0.3034 | 0.2666 | 0.0355 | gF131A | 0.1563 | 0.0305 | 0.0042 | 0.0048 |
| gF14A | 0.1239 | 0.0056 | 0.0026 | 0.0013 | gR132A | 0.2448 | 0.0534 | 0.0409 | 0.028 |
| gG15A | 0.1352 | 0.0208 | 0.0037 | 0.0022 | gL133A | 0.4262 | 0.0738 | 0.2423 | 0.1028 |
| gT16A | 0.2355 | 0.1875 | 0.0045 | 0.0029 | gL134A | 0.2597 | 0.0877 | 0.0027 | 0.0062 |
| gV17A | 0.4426 | 0.1063 | 0.0619 | 0.0051 | gL135A | 0.2652 | 0.0096 | 0.0211 | 0.0099 |
| gV18A | 0.5939 | 0.0323 | 0.3115 | 0.1565 | gN136A | 0.2927 | 0.0818 | 0.0124 | 0.0076 |
| gN19A | 0.1456 | 0.0108 | 0.0073 | 0.002 | gM137A | 0.2372 | 0.1135 | 0.0042 | 0.0056 |
| gW20A | 0.0821 | 0.0799 | 0.0029 | 0.0033 | gA138H | 0.2028 | 0.0312 | 0.0028 | 0.0033 |
| gH21A | 0.1920 | 0.0810 | 0.0023 | 0.0021 | gK139A | 0.3005 | 0.0589 | 0.1056 | 0.0386 |
| gE22A | 0.3321 | 0.1690 | 0.2808 | 0.0648 | gG140A | 1.0000 | 0.0000 | 1 | 0.069 |
| gS23A | 0.9337 | 0.0067 | 0.8136 | 0.2154 | gS141A | 0.9555 | 0.0000 | 0.6997 | 0.1393 |
| gV24A | 0.4228 | 0.0680 | 0.1697 | 0.0277 | gG142A | 0.6578 | 0.1714 | 0.2756 | 0.0375 |
| gV25A | 0.4338 | 0.0811 | 0.1404 | 0.0247 | gL143A | 0.2345 | 0.0521 | 0.0022 | 0.0021 |
| gQ26A | 0.5371 | 0.0273 | 0.3006 | 0.1335 | gP144A | 0.5462 | 0.1130 | 0.2055 | 0.0604 |
| gE27A | 0.3779 | 0.0339 | 0.21 | 0.0988 | gW145A | 0.1908 | 0.0948 | 0.003 | 0.0044 |
| gG28A | 0.2606 | 0.0716 | 0.0381 | 0.0041 | gD146A | 0.2377 | 0.0593 | 0.0081 | 0.0088 |
| gE29A | 0.8064 | 0.0000 | 0.2841 | 0.1177 | gF147A | 0.2124 | 0.0449 | 0.0025 | 0.0028 |
| gA30H | 0.1342 | 0.1485 | 0.1395 | 0.046 | gI148A | 0.2489 | 0.0631 | 0.002 | 0.0029 |
| gL31A | 0.1897 | 0.0142 | 0.0209 | 0.0009 | gL149A | 0.2147 | 0.0561 | 0.0044 | 0.0047 |
| gG32A | 0.8905 | 0.0000 | 0.2804 | 0.106 | gA150H | 0.2900 | 0.1177 | 0.0042 | 0.0042 |

| | | | | | | | | | |
|---|---|---|---|---|---|---|---|---|---|
| gR33A | 0.5721 | 0.1228 | 0.3307 | 0.0789 | gG151A | 0.9029 | 0.0240 | 0.4914 | 0.1592 |
| gA34H | 0.6273 | 0.0330 | 0.2813 | 0.1306 | gQ152A | 0.3513 | 0.1585 | 0.0242 | 0.0072 |
| gK35A | 0.4260 | 0.0418 | 0.2739 | 0.099 | gQ153A | 0.6693 | 0.0000 | 0.1983 | 0.0618 |
| gG36A | 0.3326 | 0.0699 | 0.3114 | 0.1285 | gF154A | 0.2262 | 0.0373 | 0.0043 | 0.0039 |
| gV37A | 0.5209 | 0.2900 | 0.2155 | 0.0322 | gQ155A | 0.4786 | 0.0862 | 0.1982 | 0.0906 |
| gS38A | 0.2779 | 0.0682 | 0.1922 | 0.0544 | gK156A | 0.2182 | 0.0210 | 0.008 | 0.0062 |
| gI39A | 0.2691 | 0.0499 | 0.0747 | 0.0057 | gF157A | 0.2733 | 0.0545 | 0.039 | 0.0238 |
| gD40A | 0.2873 | 0.0402 | 0.1178 | 0.0535 | gK158A | 0.2783 | 0.0679 | 0.0039 | 0.0048 |
| gW41A | 0.1173 | 0.0723 | 0.0042 | 0.0077 | gP159A | 0.2871 | 0.0266 | 0.0051 | 0.0027 |
| gH42A | 0.5786 | 0.1522 | 0.2726 | 0.1467 | gD160A | 0.3520 | 0.1605 | 0.0778 | 0.0206 |
| gE43A | 0.4584 | 0.0607 | 0.2291 | 0.0581 | gP161A | 0.5578 | 0.0000 | 0.0161 | 0.0067 |
| gF44A | 0.0754 | 0.0794 | 0.0039 | 0.0045 | gT162A | 0.4035 | 0.0104 | 0.1306 | 0.0331 |
| gA45H | 0.0601 | 0.0778 | 0.0026 | 0.0022 | gI163A | 0.2406 | 0.0261 | 0.0097 | 0.005 |
| gN46A | 0.2937 | 0.0613 | 0.1233 | 0.0301 | gY164A | 0.2500 | 0.0822 | 0.0023 | 0.0054 |
| gV47A | 0.4704 | 0.1406 | 0.2593 | 0.0668 | gE165A | 0.6613 | 0.0151 | 0.4182 | 0.0631 |
| gW48A | 0.0917 | 0.0805 | 0.0058 | 0.0064 | gD166A | 0.4622 | 0.2062 | 0.279 | 0.0799 |
| gR49A | 0.1931 | 0.0673 | 0.0054 | 0.0055 | gA167H | 0.2530 | 0.0436 | 0.0021 | 0.0067 |
| gE50A | 0.5223 | 0.0531 | 0.2676 | 0.1031 | gV168A | 0.3033 | 0.0612 | 0.0066 | 0.0069 |
| gE51A | 0.4349 | 0.0651 | 0.2505 | 0.0925 | gE169A | 0.7590 | 0.1002 | 0.5186 | 0.171 |
| gG52A | 0.3975 | 0.0503 | 0.3359 | 0.2261 | gL170A | 0.2713 | 0.0004 | 0.005 | 0.0033 |
| gY53A | 0.1848 | 0.0578 | 0.036 | 0.0188 | gL171A | 0.2433 | 0.0017 | 0.0044 | 0.0049 |
| gI54A | 0.5080 | 0.1128 | 0.329 | 0.1864 | gG172A | 0.2652 | 0.1067 | 0.003 | 0.0036 |
| gK55A | 0.3678 | 0.0505 | 0.2541 | 0.128 | gG173A | 0.2802 | 0.0894 | 0.0995 | 0.037 |
| gV56A | 0.2450 | 0.0666 | 0.0455 | 0.0201 | gR174A | 0.5235 | 0.1751 | 0.3157 | 0.0719 |
| gM57A | 0.5581 | 0.1231 | 0.2928 | 0.0608 | gP175A | 0.6953 | 0.0745 | 0.3307 | 0.0978 |
| gY58A | 0.4852 | 0.0452 | 0.1589 | 0.0702 | gE176A | 0.4983 | 0.0719 | 0.2514 | 0.1028 |
| gE59A | 0.4709 | 0.0504 | 0.2746 | 0.0514 | gE177A | 0.2446 | 0.1074 | 0.0092 | 0.0077 |
| gV60A | 0.2370 | 0.0310 | 0.1199 | 0.0404 | gV178A | 0.2768 | 0.0101 | 0.0058 | 0.0047 |
| gA61H | 0.2478 | 0.1728 | 0.0327 | 0.0225 | gL179A | 0.3916 | 0.0470 | 0.1547 | 0.0477 |
| gQ62A | 0.5896 | 0.0067 | 0.5254 | 0.2317 | gM180A | 0.2483 | 0.0757 | 0.0044 | 0.0013 |
| gG63A | 0.3292 | 0.0172 | 0.2661 | 0.0979 | gV181A | 0.2725 | 0.0413 | 0.0121 | 0.0099 |
| gL64A | 0.4749 | 0.0270 | 0.366 | 0.2342 | gA182H | 0.2373 | 0.1206 | 0.0063 | 0.0063 |
| gR65A | 0.5078 | 0.0515 | 0.3566 | 0.2015 | gA183H | 0.2509 | 0.0629 | 0.0093 | 0.0114 |
| gP66A | 0.6970 | 0.0617 | 0.329 | 0.16 | gH184A | 0.2921 | 0.0666 | 0.0027 | 0.0045 |

| | | | | | | | | | |
|---|---|---|---|---|---|---|---|---|---|
| gW67A | 0.2493 | 0.2401 | 0.0061 | 0.009 | gP185A | 0.5053 | 0.2037 | 0.2888 | 0.0741 |
| gE68A | 0.3236 | 0.2124 | 0.2107 | 0.0719 | gS186A | 0.6204 | 0.0540 | 0.2379 | 0.0088 |
| gP69A | 0.5015 | 0.2081 | 0.3838 | 0.1305 | gD187A | 0.2515 | 0.0242 | 0 | 0.0014 |
| gV70A | 0.1262 | 0.0059 | 0.0051 | 0.0066 | gL188A | 0.2600 | 0.1512 | 0.0082 | 0.0089 |
| gD71A | 0.2376 | 0.0675 | 0.0303 | 0.0183 | gD189A | 0.2884 | 0.0460 | 0.0046 | 0.0045 |
| gV72A | 0.2725 | 0.1621 | 0.008 | 0.0065 | gG190A | 0.5091 | 0.0852 | 0.1 | 0.0378 |
| gL73A | 0.1313 | 0.0161 | 0.0245 | 0.0129 | gA191H | 0.3394 | 0.1135 | 0.0063 | 0.0056 |
| gH74A | 0.3164 | 0.0074 | 0.0096 | 0.0148 | gH192A | 0.9673 | 0.0000 | 0.5221 | 0.1045 |
| gR75A | 0.3252 | 0.0015 | 0.0824 | 0.005 | gA193H | 0.5890 | 0.1499 | 0.3569 | 0.0832 |
| gR76A | 0.3043 | 0.0319 | 0.247 | 0.0861 | gI194A | 0.2272 | 0.0416 | 0.0409 | 0.0079 |
| gK77A | 0.3762 | 0.1182 | 0.1204 | 0.0269 | gG195A | 0.0892 | 0.0091 | 0.0075 | 0.0139 |
| gL78A | 0.1798 | NA | 0.0139 | 0.0188 | gC196A | 0.2361 | 0.0976 | 0.0327 | 0.0467 |
| gD79A | 0.4822 | 0.0604 | 0.5085 | 0.3439 | gP197A | 0.4561 | 0.0539 | 0.2688 | 0.124 |
| gE80A | 0.5469 | 0.0384 | 0.44 | 0.3203 | gT198A | 0.0788 | 0.0475 | 0.0078 | 0.0072 |
| gL81A | 0.2500 | 0.0174 | 0.0077 | 0.0095 | gL199A | 0.2781 | 0.0214 | 0.2234 | 0.0848 |
| gL82A | 0.6384 | 0.0547 | 0.2554 | 0.0728 | gY200A | 0.1339 | 0.1153 | 0.1539 | 0.0697 |
| gD83A | 0.5906 | 0.1441 | 0.3814 | 0.1799 | gV201A | 0.0952 | 0.0561 | 0.0122 | 0.0053 |
| gV84A | 0.5399 | 0.0607 | 0.2937 | 0.0837 | gP202A | 0.2310 | 0.0667 | 0.0586 | 0.0252 |
| gY85A | 0.4632 | 0.1722 | 0.2214 | 0.1356 | gR203A | 0.0714 | 0.0254 | 0.0047 | 0.0065 |
| gG86A | 0.4495 | 0.0882 | 0.4361 | 0.3551 | gP204A | 0.3489 | 0.0212 | 0.2112 | 0.0217 |
| gL87A | 0.2879 | 0.0689 | 0.0167 | 0.0096 | gL205A | 0.4286 | 0.0004 | 0.347 | 0.1157 |
| gK88A | 0.5715 | 0.0559 | 0.3104 | 0.1286 | gE206A | 0.1005 | 0.0161 | 0.005 | 0.008 |
| gL89A | 0.5111 | 0.0517 | 0.106 | 0.0398 | gY207A | 0.0644 | 0.0231 | 0.0012 | 0.0019 |
| gT90A | 0.7042 | 0.0915 | 0.3317 | 0.2088 | gG208A | 0.0968 | 0.0158 | 0.009 | 0.0138 |
| gE91A | 0.6371 | 0.0472 | 0.3696 | 0.198 | gA209H | 0.3107 | 0.0465 | 0.2968 | 0.1262 |
| gE92A | 0.6167 | 0.1010 | 0.4529 | 0.3318 | gV210A | 0.3492 | 0.0398 | 0.327 | 0.1268 |
| gE93A | 0.4263 | 0.1783 | 0.066 | 0.009 | gN211A | 0.3123 | 0.1716 | 0.269 | 0.0808 |
| gT94A | 0.4140 | 0.2512 | 0.4376 | 0.2855 | gN212A | 0.4144 | 0.0669 | 0.2726 | 0.0739 |
| gD95A | 0.6896 | 0.0104 | 0.4895 | 0.208 | gH213A | 0.1448 | 0.0490 | 0.0267 | 0.0058 |
| gH96A | 0.5300 | 0.0482 | 0.3414 | 0.0839 | gV214A | 0.3361 | 0.0061 | 0.2408 | 0.0085 |
| gF97A | 0.2515 | 0.0386 | 0.0118 | 0.0089 | gE215A | 0.1204 | 0.0215 | 0.0644 | 0.0229 |
| gN98A | 0.5823 | 0.0104 | 0.2869 | 0.0369 | gP216A | 0.3582 | 0.0325 | 0.2317 | 0.0922 |
| gR99A | 0.7416 | 0.0000 | 0.2851 | 0.009 | gE217A | 0.3649 | 0.0522 | 0.3208 | 0.1874 |
| gL100A | 0.7226 | 0.0000 | 0.277 | 0.0627 | gA218H | 0.2716 | 0.0761 | 0.1923 | 0.0739 |

| | | | | | | | | | |
|---|---|---|---|---|---|---|---|---|---|
| gW101A | 0.1031 | 0.0675 | 0.0052 | 0.0027 | gK219A | 0.2922 | 0.0481 | 0.2751 | 0.0909 |
| gH102A | 0.3459 | 0.1577 | 0.0128 | 0.0045 | gY220A | 0.1752 | 0.0176 | 0.0339 | 0.0088 |
| gR103A | 0.3532 | 0.0166 | 0.0248 | 0.0179 | gD221A | 0.3783 | 0.0992 | 0.3449 | 0.136 |
| gL104A | 0.5109 | 0.1236 | 0.2282 | 0.0208 | gH222A | 0.0822 | 0.0700 | 0.0064 | 0.0037 |
| gL105A | 0.8244 | 0.1746 | 0.4598 | 0.0579 | gE223A | 0.2384 | 0.0176 | 0.2549 | 0.0697 |
| gP106A | 0.3295 | 0.0338 | 0.0803 | 0.012 | gT224A | 0.3693 | 0.0012 | 0.3114 | 0.0795 |
| gW107A | 0.1160 | 0.1174 | 0.0006 | 0.0014 | gV225A | 0.2552 | 0.0614 | 0.169 | 0.0392 |
| gP108A | 0.5080 | 0.0603 | 0.2411 | 0.0653 | gA226H | 0.3996 | 0.1008 | 0.2902 | 0.0724 |
| gD109A | 0.2215 | 0.0000 | 0.0032 | 0.004 | gD227A | 0.0815 | 0.0107 | 0.0007 | 0.001 |
| gV110A | 0.4678 | 0.2630 | 0.3849 | 0.2342 | gF228A | 0.1122 | 0.0185 | 0.0016 | 0.0022 |
| gQ111A | 0.5112 | 0.0189 | 0.3602 | 0.1944 | gR229A | 0.2787 | 0.1243 | 0.2526 | 0.095 |
| gE112A | 0.4456 | 0.1612 | 0.2952 | 0.0103 | gE230A | 0.8656 | 0.0000 | 0.3191 | 0.1007 |
| gG113A | 0.7484 | 0.1985 | 0.5705 | 0.0881 | gL231A | 0.0313 | 0.0443 | 0.0077 | 0.006 |
| gL114A | 0.1624 | 0.0138 | 0.0047 | 0.0039 | gA232H | 0.2064 | 0.0488 | 0.0243 | 0.0154 |
| gR115A | 0.5597 | 0.1369 | 0.4441 | 0.0455 | gA233H | 0.3840 | 0.0031 | 0.3409 | 0.0939 |
| gR116A | 0.5653 | 0.0367 | 0.3634 | 0.0763 | gR234A | 0.3410 | 0.0257 | 0.3396 | 0.1373 |
| gL117A | 0.1191 | 0.0117 | 0.0037 | 0.004 | gL235A | 0.0708 | 0.0419 | 0.01 | 0.0091 |
| gK118A | 0.4547 | 0.2647 | 0.2779 | 0.0274 | gG236A | 0.3461 | 0.0036 | 0.0996 | 0.0735 |
| gT119A | 0.4841 | 0.0391 | 0.3245 | 0.0993 | gV237A | 0.4622 | 0.0193 | 0.3715 | 0.1677 |

**Table S19**: Organisms used in this study

| **Strains** | **Genotype and specifications** | **Origin** |
|---|---|---|
| *E. coli* DH5α | *fhuA2Δ(argF-lacZ)U169 phoA glnV44 Φ80Δ(lacZ)M15 gyrA96 recA1 relA1 endA1 thi-1 hsdR17* | New England BioLabs |
| T7 Express *E. coli* | *fhuA2 lacZ::T7 gene1 [lon] ompT gal sulA11 R(mcr-73::miniTn10*--Tet$^S$*)2 [dcm] R(zgb-210::Tn10*--Tet$^S$*) endA1 Δ(mcrC-mrr)114::IS10* | New England BioLabs |

| | | |
|---|---|---|
| *Gordonibacter pamelaeae* DSM 110924 | WT, isolated from human feces | DSMZ |

**Table S20:** Plasmids generated in this study

| Plasmid name | Parent plasmid | Description | Inducer |
|---|---|---|---|
| pCDFDuet-1-WP118709078 (P1) | pCDFDuet-1 | *spec*$^{R}$, *wp118709078.1* | IPTG |
| pCDFDuet-1-WP087189991 (P2) | pCDFDuet-1 | *spec*$^{R}$, *wp087189991.1* | IPTG |
| pCDFDuet-1-GKG91607 (P3) | pCDFDuet-1 | *spec*$^{R}$, *gkg91607* | IPTG |
| pCDFDuet-1-WP004693440 (P4) | pCDFDuet-1 | *spec*$^{R}$, *wp004693440.1* | IPTG |
| pCDFDuet-1-3UMG (P5-control) | pCDFDuet-1 | *spec*$^{R}$, *3UMG-1* | IPTG |
| pCDFDuet-1-WP178618037 (P6) | pCDFDuet-1 | *spec*$^{R}$, *wp178618037.1* | IPTG |
| pCDFDuet-1-WP006686258 (P7) | pCDFDuet-1 | *spec*$^{R}$, *wp006686258.1* | IPTG |
| pCDFDuet-1-WP053428405 (P8) | pCDFDuet-1 | *spec*$^{R}$, *wp053428405.1* | IPTG |
| pCDFDuet-1-WP050688410 (P9) | pCDFDuet-1 | *spec*$^{R}$, *wp*050688410.*1* | IPTG |
| pCDFDuet-1-WP029378231 (P10) | pCDFDuet-1 | *spec*$^{R}$, *wp*029378231.*1* | IPTG |
| pCDFDuet-1-WP048310185 (P11) | pCDFDuet-1 | *spec*$^{R}$, *wp048310185.1* | IPTG |
| pCDFDuet-1-WP081745164 (P12) | pCDFDuet-1 | *spec*$^{R}$, *wp*081745164.*1* | IPTG |
| pCDFDuet-1-WP066861885 (P13) | pCDFDuet-1 | *spec*$^{R}$, *wp066861885.1* | IPTG |
| pCDFDuet-1-WP263041397 (P14) | pCDFDuet-1 | *spec*$^{R}$, *wp*263041397.*1* | IPTG |
| pCDFDuet-1-WP187031722 (P15) | pCDFDuet-1 | *spec*$^{R}$, *wp*187031722.*1* | IPTG |
| pCDFDuet-1-WP036125194 (P16) | pCDFDuet-1 | *spec*$^{R}$, *wp036125194.1* | IPTG |
| pCDFDuet-1-WP031546286 (P17) | pCDFDuet-1 | *spec*$^{R}$, *wp031546286.1* | IPTG |
| pCDFDuet-1-WP053359908 (P18) | pCDFDuet-1 | *spec*$^{R}$, *wp053359908.1* | IPTG |
| pCDFDuet-1-GKH53471 (P19) | pCDFDuet-1 | *spec*$^{R}$, *gkh53471* | IPTG |

| | | | |
|---|---|---|---|
| pCDFDuet-1-WP13943301 (P20) | pCDFDuet-1 | *spec$^R$, wp13943301.1* | IPTG |
| pCDFDuet-1-HTS9340 (P21) | pCDFDuet-1 | *spec$^R$, hts9340* | IPTG |
| pCDFDuet-1-PYN88313 (P22) | pCDFDuet-1 | *spec$^R$, pyn88313* | IPTG |
| pCDFDuet-1-MDH3205452 (P23) | pCDFDuet-1 | *spec$^R$, mdh3205452* | IPTG |
| pCDFDuet-1-MBS6900484 (P24) | pCDFDuet-1 | *spec$^R$, mbs6900484* | IPTG |
| pCDFDuet-1-WP118709078-F54I | pCDFDuet-1 | *spec$^R$*, WP118709078-F54I | IPTG |
| pCDFDuet-1-WP118709078-T57M | pCDFDuet-1 | *spec$^R$*, WP118709078-T57M | IPTG |
| pCDFDuet-1-WP118709078-F74H | pCDFDuet-1 | *spec$^R$*, WP118709078-F74H | IPTG |
| pCDFDuet-1-WP118709078-F187S | pCDFDuet-1 | *spec$^R$*, WP118709078-F187S | IPTG |
| pCDFDuet-1-WP118709078-F54I-T57M | pCDFDuet-1 | *spec$^R$*, WP118709078-F54I-T57M | IPTG |
| pCDFDuet-1-WP118709078-F54I-F74H | pCDFDuet-1 | *spec$^R$*, WP118709078-F54I-F74H | IPTG |
| pCDFDuet-1-WP118709078-F54I-F187S | pCDFDuet-1 | *spec$^R$*, WP118709078-F54I-F187S | IPTG |
| pCDFDuet-1-WP118709078-F54I-T57M-F74H | pCDFDuet-1 | *spec$^R$*, WP118709078-F54I-T57M-F74H | IPTG |
| pCDFDuet-1-WP118709078-F54I-T57M-F187S | pCDFDuet-1 | *spec$^R$*, WP118709078-F54I-T57M-F187S | IPTG |
| pCDFDuet-1-WP118709078-F54I-F74H-F187S | pCDFDuet-1 | *spec$^R$*, WP118709078-F54I-F74H-F187S | IPTG |
| pCDFDuet-1-WP118709078-F54I-T57M-F74H-F187S | pCDFDuet-1 | *spec$^R$*, WP118709078-F54I-T57M-F74H-F187S | IPTG |
| pCDFDuet-1-WP1187090781-F54I-T57M-F74H-F187S-1-35-WP1786180371 | pCDFDuet-1 | *spec$^R$*, pCDFDuet-1-WP1187090781-F54I-T57M-F74H-F187S AA substitutes 1-35 with WP1786180371 | IPTG |
| pCDFDuet-1-WP-1187090781-F54I-T57M-F74H-F187S-36-69-WP1786180371 | pCDFDuet-1 | *spec$^R$*, pCDFDuet-1-WP1187090781-F54I-T57M-F74H-F187S AA substitutes 36-69 with WP1786180371 | IPTG |
| pCDFDuet-1-WP-1187090781-F54I-T57M-F74H-F187S-70-109-WP1786180371 | pCDFDuet-1 | *spec$^R$*, pCDFDuet-1-WP1187090781-F54I-T57M-F74H-F187S AA substitutes 70-109 with WP1786180371 | IPTG |

| | | | |
|---|---|---|---|
| pCDFDuet-1-WP-1187090781-F54I-T57M-F74H-F187S-110-150-WP1786180371 | pCDFDuet-1 | *spec*$^R$, pCDFDuet-1-WP1187090781-F54I-T57M-F74H-F187S AA substitutes 110-150 with WP1786180371 | IPTG |
| pCDFDuet-1-WP-1187090781-F54I-T57M-F74H-F187S-150-196-WP1786180371 | pCDFDuet-1 | *spec*$^R$, pCDFDuet-1-WP1187090781-F54I-T57M-F74H-F187S AA substitutes 150-196 with WP1786180371 | IPTG |
| pCDFDuet-1-WP-1187090781-F54I-T57M-F74H-F187S-197-end-WP1786180371 | pCDFDuet-1 | *spec*$^R$, pCDFDuet-1-WP1187090781-F54I-T57M-F74H-F187S AA substitutes 197-238 with WP1786180371 | IPTG |

**Table S21:** Primers used in this study

| Primer name | Sequence |
|---|---|
| **Site directed mutagenesis P1** | |
| FW-WP118709078-F54I | CGCAATGATGGCTATattAAGGCAACC |
| RV-WP118709078-F54I | CCATCCGTTGGCAAATTCATCCCAATC |
| FW-WP118709078-T57M | GCAatgTTTGATATCGCCCATGGTAAGC |
| RV-WP118709078-T57M | CTTAAAATAGCCATCATTGCGCCATCCG |
| FW-WP118709078-F54T-T57M | CTATattAAGGCAatgTTTGATATCG |
| RV-WP118709078-F54T-T57M | CCATCATTGCGCCATCCG |
| FW-WP118709078-F74H | CTGATACTGTGcatATGGAATACC |
| RV-WP118709078-F74H | CAGGCACCCATTCACGC |
| FW-WP118709078-F187S | CTCATCCTagtGACCTTGAC |
| RV-WP118709078-F187S | CGGCAACCATTGCAACCTC |
| **Chimeric protein P1/P6 segment AA1-36** | |
| FW-WP118709078-1-36 | GTGCCAAATCGTTAGATTTAGATTGGGATGAATTTGC |
| RV-WP118709078-1-36 | GCTTACATCCATGTATATCTCCTTATTAAAGTTAAACAAAATTATTTC |
| FW-WP1786180371-1-35 | GGAGATATACATGGATGTAAGCAATGTGCGTATTG |

| | |
|---|---|
| RV-WP1786180371-1-35 | CTAAATCTAACGATTTGGCACGCCCTAATGCTTC |

**Chimeric protein P1/P6 segment AA 36-69**

| | |
|---|---|
| FW-WP1786180371-36-69 | CGAAAAATACGGGGTTAGTATTGACTGGCAC |
| RV-WP1786180371-36-69 | GTATCAGCAGGTTCCCAAGGGCGC |
| RV-WP118709078-36-69 | CAATACTAACCCCGTATTTTTCGTTTAACGCTTGAC |
| FW-WP118709078-36-69 | CCTGCTGATACTGTGcatATGGAATACCTTG |

**Chimeric protein P1/P6 segment AA 70-109**

| | |
|---|---|
| FW-WP1786180371-70-109 | GTGCCTGTTGACGTGTTGCATCGTCGC |
| RV-WP1786180371-70-109 | CAGGACGTCCGGCCATGGAAGCAG |
| FW-WP118709078-70-109 | CCGGACGTCCTGGAGGGGCTGAATCG |
| RV-WP118709078-70-109 | CACGTCAACAGGCACCCATTCACGCTTACC |

**Chimeric protein P1/P6 segment AA 110-150**

| | |
|---|---|
| FW-WP1786180371-110-150 | CCGGATGTGCAGGAGGGGCTGCG |
| RV-WP1786180371-110-150 | GAATAAGTCCCCTGCTAAAATGAAGTCCCACGGTAAG |
| FW-WP118709078-110-150 | CATTTTAGCAGGGGACTTATTCCAGAAGTTC |
| RV-WP118709078-110-150 | CTGCACATCCGGCCACGGTGAAAG |

**Chimeric protein P1/P6 segment AA 151-196**

| | |
|---|---|
| FW-WP1786180371-151-196 | CGACCGGCCAGCAGTTCCAGAAATTTAAG |
| RV-WP1786180371-151-196 | GTGGTACATCCGATTGCGTGCGCG |
| FW-WP118709078-151-196 | CAATCGGATGTACCACCATCTTCGTGCCGC |
| RV-WP118709078-151-196 | GCTGGCCGGTCGTAATAAAATCCCACGGC |

**Chimeric protein P1/P6 segment AA 197-238**

| | |
|---|---|
| FW-WP1786180371-197-238 | GGCAGGGTGCCCAACCTTATATGTACCTCGC |
| RV-WP1786180371-197-238 | CAGATTCTCGACTCCAAGGCGAGCGG |

| | |
|---|---|
| FW-WP118709078-197-238 | CCTTGGAGTCGAGAATCTGTATTTTCAGAGCCATCACC |
| RV-WP118709078-197-238 | CATATAAGGTTGGGCACCCTGCCAATTTAGCTCCGTC |
